# Features that matter: evolutionary signatures that predict viral transmission routes

**DOI:** 10.1101/2023.11.22.568327

**Authors:** Maya Wardeh, Jack Pilgrim, Melody Hui, Aurelia Kotsiri, Matthew Baylis, Marcus SC Blagrove

**Affiliations:** Department of Computer Science, University of Liverpool, Liverpool, UK; Institute of Infection, Veterinary and Ecological Sciences, University of Liverpool, Liverpool, UK

## Abstract

Routes of virus transmission between hosts are key to understanding viral epidemiology. Different routes have large effects on viral ecology, and likelihood and rate of transmission. For example, respiratory and vector-borne viruses together encompass the majority of high-consequence animal and plant outbreaks. However, the specific transmission route(s) can take months to years to determine, undermining the efficiency of mitigation efforts. Here, we identify the vial features and evolutionary signatures which are predictive of viral transmission routes, and use them to predict potential routes for fully-sequenced viruses – we perform this for both viruses with no observed routes, as well as viruses with missing routes. This was achieved by compiling a dataset of 24,953 virus-host associations with 81 defined transmission routes, constructing a hierarchy of virus transmission encompassing those routes and 42 higher-order modes, and engineering 446 predictive features from three complementary perspectives. We integrated those data and features, to train 98 independent ensembles of LightGBM classifiers, each incorporating five different class-balancing approaches. Using our trained ensembles, we demonstrated that all features contributed to the prediction for at least one of routes and/or modes of transmission, demonstrating the utility of our multi-perspective approach. Our approach achieved ROC-AUC=0.991, and F1-score=0.855 across all modelled transmission mechanisms; and was able to achieve high levels of predictive performance for high-consequence respiratory (ROC-AUC=0.990, and F1-score=0.864) and vector-borne transmission (ROC-AUC=0.997, and F1-score=0.921). Our framework ranks the viral features in order of their contribution to prediction, per transmission route, and hence identifies the genomic evolutionary signatures associated with each route. Together with the more matured field of viral host-range prediction, our predictive framework could: provide early insights into the potential for, and pattern of viral spread; facilitate rapid response with appropriate measures; and significantly triage the time-consuming investigations to confirm the likely routes of transmission. Moreover, the performance of our approach in high-consequence transmission routes showcases that our methodology has direct utility to pandemic preparedness.

**AUTHORS SUMMARY:** Routes of virus transmission – the mechanism(s) by which a virus physically gets from an infected to an uninfected host, are crucial to understanding how viral diseases spread among animals and plants. Here, we uncover the evolutionary signatures which can predict the transmission routes a virus uses to move from one host to another, enabling us to identify any unobserved routes for known viruses and even predict potential routes of newly emerged viruses. We first compile a comprehensive dataset of virus-host associations. Leveraging this dataset, we employ a multi-perspective machine learning approach to achieve high predictive performance. Our framework ranks viral features by their significance in prediction, revealing genomic evolutionary signatures linked to each route. Our approach could provide early insights into viral spread patterns, facilitating prompt response efforts to new outbreaks and epidemics, and streamline laboratory investigations. Overall, our study represents a step forward in our ability to anticipate and mitigate the impact of emerging infectious diseases on human, animal, and plant health.

## INTRODUCTION

Mounting an effective response to an emerging virus requires establishing all critical information, as quickly as possible. In recent years, significant focus has been placed on understanding determinants of host range and potential for spillover [1–5]. However, the transmission route, i.e. the pathway a virus uses to physically get from an infected to an uninfected host, still entails months, or even years, to thoroughly investigate. This was most apparent during the initial phases of the SARS-CoV-2 pandemic, where the relative importance of lingering aerosols versus fomite transmission was still being determined [6,7]. Furthermore, secondary transmission routes, such as sexual transmission of both Zika [8] and Ebola [9] viruses, are often only ascertained during, or after, significant outbreaks. The ability to identify all epidemiologically significant transmission routes of a virus, with high accuracy, computationally, with minimal information, and as quickly as possible, is therefore of paramount importance to future emerging viruses.

The transmission routes of a virus are also fundamentally intertwined with its ecology, epidemiology [10], and its potential for host shifting and spillover [11]. They determine how the virus spreads within and between different host populations, therefore influencing the potential, severity, and geographical extent of outbreaks. In animals, transmission routes such as respiratory, via droplets or aerosol, can result in rapid virus spread through a dense population. Influenza A, many coronaviruses, and the more benign rhinoviruses, all benefit from this transmission mechanism to cause swift outbreaks worldwide [12]. Conversely, vector-borne viruses tend toward a more varied outbreak speed, for instance the gradual spread of Usutu across Europe [13], compared with the El Niño-driven rapid spread of Zika through South America [14]. But nonetheless, vector-borne routes can produce wide ranging and long-term establishment, for example dengue[15], bluetongue [16], and Tomato Yellow Leaf Curl [17] viruses. Different epidemiological patterns are also seen for other sets of transmission routes, such as vertical, sexual, and water-borne [10,18].

In the plant kingdom, most viruses are transmitted by vectors, particularly by hemipterans insects, such as aphids and whiteflies [19]. The dynamics of this transmission by the vector: non-persistent, semi-persistent, or persistent, determine the length of window to disseminate the virus to a new plant after feeding (seconds to minutes, hours to days, or days to weeks, respectively) [20]. Vertical transmission via seeds, on the other hand, enables the virus to persist for considerably long periods when hosts or vectors are not available [21], and may allow it to disseminate over long distances, including continental jumps [22].

To facilitate computational prediction of transmission routes, we first compiled a dataset of known transmission routes of the animal and plant viromes, to their hosts, from published literature, and used these data to construct a hierarchy of transmission mechanisms. We then established a field-bridging and uniform methodology to define routes of transmission based on each virus-host association rather than a ‘per virus’ definition, because, in some cases, the same species or strain of virus may utilise a different range of transmission routes to infect different hosts. For instance, Influenza A is faecal-orally transmitted in waterfowl [23], but undergoes respiratory transmission in humans [24]. In other cases, very closely related, but different, viruses may utilise a diverse set of transmission routes in different hosts. For instance, whilst many viruses in the family Flaviviridae are exclusively vector-borne, some are also vertically or sexually transmitted in both vertebrate and vector populations [25], further some, e.g. Hepatitis C virus, are blood-borne and do not replicate in arthropods [26].

Given the above two scenarios, we incorporated pair-wise association-level similarities into a unified framework, termed: ‘Virus-host integrated neighbourhoods’, to synthesise complementary predictive features. Furthermore, in order to enable parameterisation of transmission routes that are closely interlinked with host taxonomy (e.g. seed and pollen-borne routes are strictly limited to plants, milk-borne transmission in mammals, and egg related routes in non-mammalian animals), we incorporated similarity between hosts to differentiate between those categorically different routes.

Finally, as virus structure has been shown to constrain virus transmission [27,28], and biases in the virus genome composition can also inform the transmission mechanism deployed by the virus, as well as correlating with virus reservoirs and vectors [29], we synthesised a complementary array of variables from the full genome sequences of viruses.

We combined the above features and viral evolutionary signatures into lightGBM ensembles. We used those ensembles to identify which of our features are most predictive of transmission routes deployed by animal- and plant-infecting viruses to their known hosts; to predict which of those mechanisms are applicable to virus-host associations without observed transmission routes; and to establish potential gaps in our current knowledge of the transmission routes of known viruses to their animal or plant hosts.

This study is the most taxonomically broad study of its kind, to demonstrate the potential of sequence and morphological information, increasingly available within the first few days of an outbreak, to predict the transmission pathways of a novel virus, across the animal and plant viromes. Deployment of our suggested framework could provide early insights into the potential for, and pattern of spread of a virus; facilitate rapid response with appropriate measures; and significantly triage the time-consuming investigations to confirm the likely routes.

## RESULTS

### Hierarchy of virus transmission

We captured data on 81 *non-mutually exclusive routes* of 4,446 viruses to 5,317 animal and plant species (a total of 24,953 virus-host associations, Supplementary Table 3, Supplementary Datasets 1-2), by performing a series of complementary literature searches (see methods).

Where at least one *route* of transmission was identified, we used those data to populate higher levels (*modes*) in our hierarchy. Figure 1 illustrates the distribution of observed transmission routes and modes of virus-host associations across our suggested hierarchy (Figure 1.A), as well as between our viruses and hosts (Figures 1.B and 1.C).

**Figure 1.**
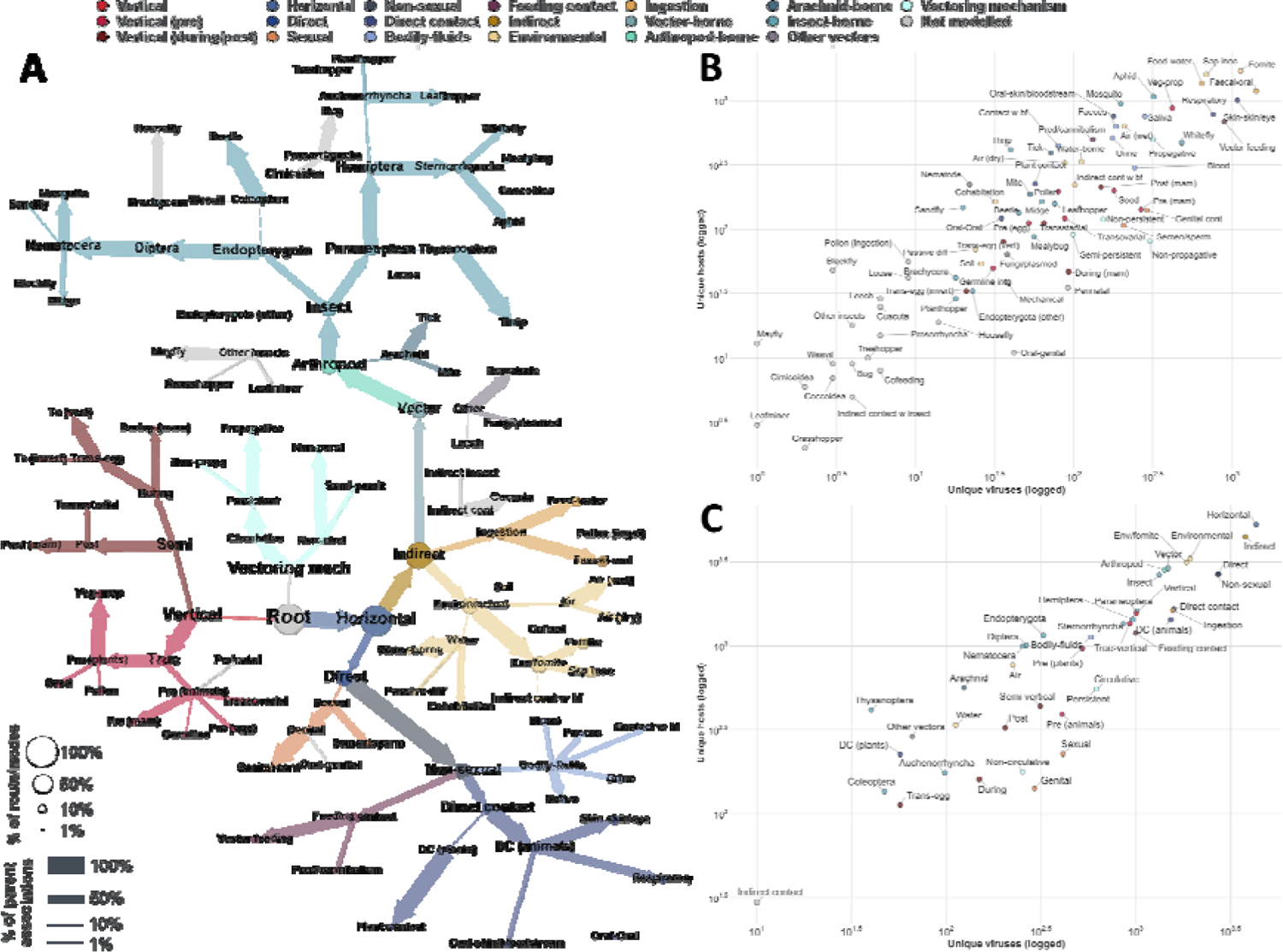
Overview of observed transmission routes/modes. **Panel A – Our suggested hierarchy.** Nodes represent transmission routes/modes identified in this study. Edges link parent modes (nodes with at least one child) with their offspring (e.g. indirect and direct transmission are two modes of horizontal transmission). Nodes and edges are coloured by the mode of transmission; routes not modelled in this study (due to insufficient data, n=18, modelled routes = 59), and conceptual nodes (root and vectoring mechanism) are coloured in light grey. Node size is proportional to percent of unique virus-host associations (of 24,953 associations) where the virus is transmitted to the host species via the corresponding route/mode. Thickness of edges is proportional to the percent of the parent associations identified to where the virus transmitted to the host species via the child route/mode (e.g. of 10,021 associations transmitted by arthropods [40.16% of included associations], 92.63% are insect-borne, and 9.5% are arachnid-borne). Supplementary Figure 5 visualises for each unique route/mode pair (no route/mode in the pair is an ancestor/offspring of the other), the percent of virus-host associations (of total included), whereby the virus is known to be transmitted to the host via both pathways. **Panel B – Transmission routes identified in this study.** Points represent transmission *routes* (Supplementary Table 3) and are coloured by transmission modes. X axis represents the number of observed unique viruses per route. Y axis represents the number of observed unique host species per route. **Panel C – Transmission modes identified in this study.** Given a virus-host association, we considered the virus to be transmitted to the host via a parent *mode* (e.g. Zika virus is insect-borne to humans), if it were transmitted by at least one *route* that is also an offspring/descendant of the parent mode (e.g. zika virus is mosquito-borne to humans), in our hierarchy (Figure 1.A).

For arthropod-borne viruses, the route is different from vector-to-host compared to host-to-vector, hence, we captured the mechanism of their transmission by the vector to the vertebrate or plant host, as well as from to the arthropod vector to the vertebrate/plant (and between vectors where relevant). For example: Zika virus is mosquito-borne to humans, but mosquitoes become infected with Zika via feeding on humans (arthropod feeding), additionally Zika is transovarially/sexually transmitted in some mosquito species. Additionally, we captured the dynamics of vector-transmission, e.g. whether the virus is transmitted mechanically, or if it replicates within the vector, which we termed vectoring mechanism (4 routes, 2 modes). For instance, Tomato Yellow Leaf Curl virus (TYLCV) is whitefly-borne to tomato (and other) plants, and is circulative, non-propagative in its whitefly vector.

To facilitate the construction of our transmission hierarchy, spanning viruses of humans, animals, and plants, we unified certain transmission pathways into route names that may not be widely used (Supplementary Table 3), these include “Air (dry)” and “Air (wet)” – which we used to describe transmission via inhalation of virus particles from the environment, but which may be confused with airborne transmission, commonly used to describe individual-to-individual transmission via droplets or airborne particles, termed respiratory in our study. We elected to separate environmental airborne transmission, from individual-to-individual respiratory transmission as these routes have different epidemiological implications, environmental viruses may persist in the environment for longer period, and do not require direct contact between the individual, for instance hantaviruses are transmissible by inhalation of virus particles from rodent urine (Air (wet)), and many avian viruses are transmitted via inhalation of dust (Air (dry)).

### Predictors of transmission routes/modes

Of a total of 446 features, our virus-host integrated neighbourhoods (‘MN4D’ and ‘MN3H’, see Methods and Supplementary Note 5) and hosts similarity (‘hosts’) features made the largest contributions to predictions across all routes/modes (Figure 2): ‘MN4D’ (top predictor of 51.02% of 98 routes/modes with sufficient data for modelling, see methods), ‘hosts’ (45.19%), and ‘MN3H’ (3.06%). These three features were in the top ten predictors of 96.94%, 98.98% and 72.45% of our route/modes, respectively (Figure 2, Supplementary Dataset 3). The remainder integrated neighbourhoods feature: ‘MN4C’ (see methods, Supplementary Note 5), was a top ten predictor in 45.92% of routes/modes. Supplementary Dataset 3 provides ranking, and local and global SHAP values of top ten predictors of each route/mode.

**Figure 2.**
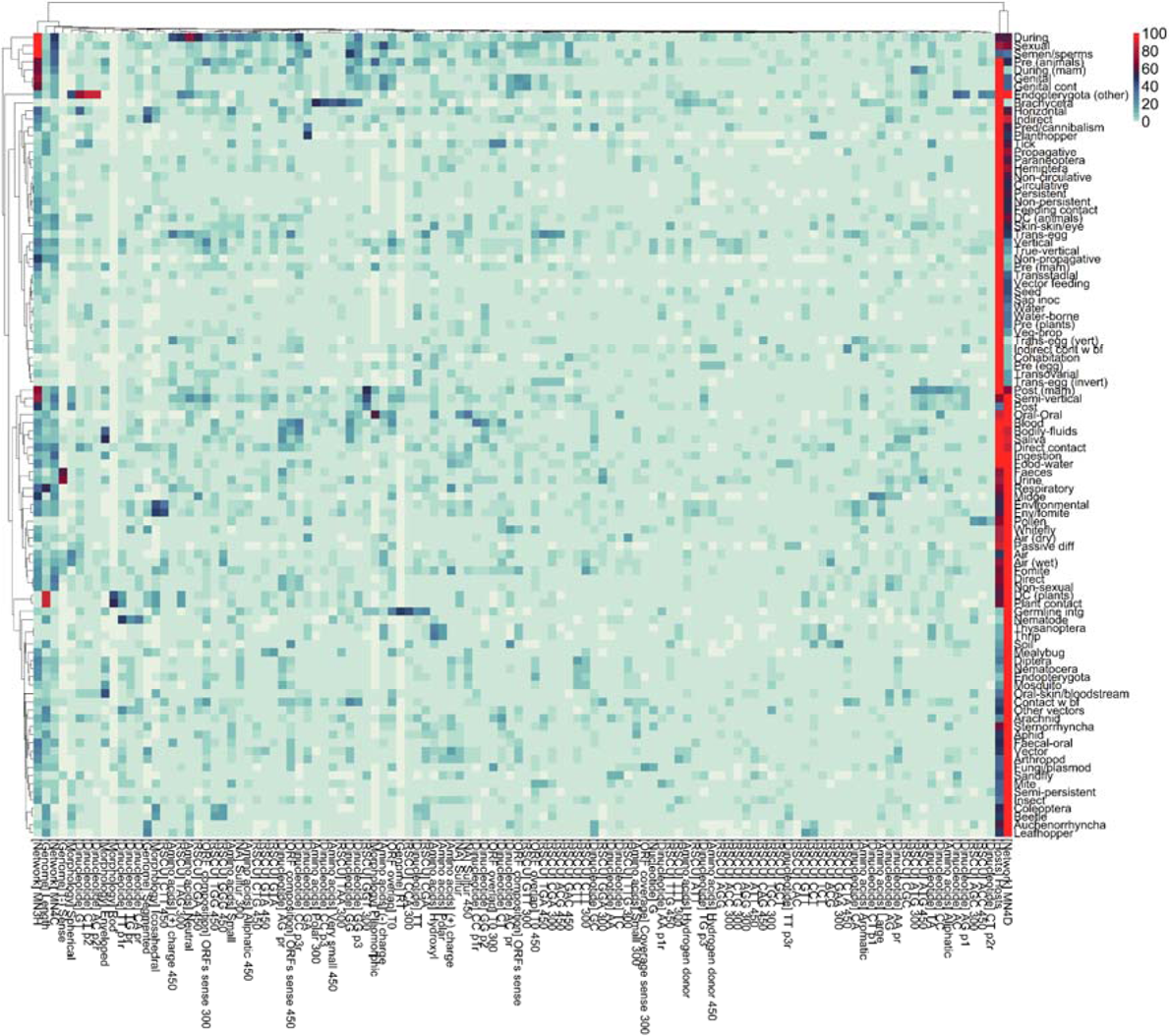
Top 10 predictors of modelled routes/modes. Mean absolute SHAP values were normalised, separately, for each route/mode modelled in this study (scale 1:100, formula = 100*mean SHAP value/max (SHAP value)). Features were ordered by the descending value of the locally normalised SHAP values and the top 10 were selected per each route/mode. The heatmap visualises the contribution (locally normalised SHAP values) the resulting 116 features (Y-axis) made to the predictions of the 98 route/mode modelled in this study (X-axis). We performed hierarchical clustering on both rows and columns, using the R package pheatmap. the resulting dendrogram is displayed (top and left).

We compiled nine genomic structure features (Supplementary Table 5 fully describes viral features utilised in this study), of which genome ‘length’, and whether the virus is ‘segmented’ or not were the most contributing, featuring in the top ten predictors of 39.8% and 13.3%, respectively. ‘Length’ made a significant contribution (locally normalised mean absolute SHAP value ≥20) to the predictions of five routes/modes (Figure 2, Supplementary Dataset 3) including respiratory (locally normalised absolute SHAP value = 85.92, vs globally normalised mean absolute SHAP value = 30.86) and Arachnid-borne transmission (24.27 vs global 9.91), where it ranked as 3^rd^ predictor. ‘Segmented’ made similar contribution to four routes/modes including indirect (40.73 vs 17.07) and horizontal (32.76, 7.34) transmission modes.

Of our 16 virus morphology features, five were in the top ten predictors of our routes/modes, with ‘icosahedral’ (17.35%), ‘enveloped’ (14.29%) and ‘spherical’ (10.2%) being top ten predictors of >10% of our routes/modes. These features contributed significantly to the predictions of 14 routes/modes, including: ‘enveloped’ (indicating if the virus is enveloped or not) for oral-skin/bloodstream (42 vs 20.44), and saliva transmission (50.81 vs 15.23); ‘pleomorphic’ capsid for oral-oral transmission (57.68 vs 13.98), and ‘spherical’ capsid for environmental transmission via inhalation of wet virus particles (e.g. urine) (24.21 vs 8.62).

Our framework incorporated features representative of the sizes of the ORFs (composition, 9 features), the proportion of genome utilised by ORFs (coverage, 21 features), and the proportion of overlapping ORFs (overlap, 21 features). Three of our ORF composition features: ‘ORFs sense 450’, ‘ORFs sense 300’, and ‘ORFs sense’ (number of predicted ORFs, of length ≥ 450, ≥ 300, and ≥ 36, respectively, divided by length of sequence) were top ten predictors for 14.29%, 12.24%, and 12.24% of our routes/modes, respectively. These features were important predictors for nine routes/modes, including 6 vertical transmission pathways (Supplementary Dataset 3). Of 21 ORF overlap features ‘T0’ (proportion of the sequence not featuring in any predicted ORF of length ≥ 36) was the only top ten predictors for >10% of routes/modes (15.31%), with most significant contribution to transmission by germline integration (25.83 vs 8.44), and by blood (23.34 vs 6.95). Finally, only ‘coverage sense 300’ (proportion of genome covered with predicted ORFs with length ≥300) featured as a top ten predictor (fungi/plasmodiophorids transmission) of our ORF coverage features.

We computed 57 biases in amino acids (from predicted ORFs, Supplementary Note 3), 19 of which were top ten predictors for at least one route/mode, and six were top ten predictors for >10% of our routes/modes: ‘sulphur’ (13.27%), ‘sulphur 450’ (13.27%), ‘(+) charge’ (12.24%), ‘(+) charge 450’ (12.24%), ‘hydroxyl’ (11.22%), and ‘(-) charge’ (10.2%). Our amino acid biases made significant contribution to 21 routes/modes, including: ‘(+) charge’ for fomite (ranked 3^rd^), and respiratory transmission (5^th^), and ‘hydroxyl’ for thrip-borne transmission (ranked 2^nd^), and ‘neutral’ for (vertical) ‘during’ transmission mode incorporating all transmission routes in which the virus is transmitted during birth or egg hatching (ranked 2^nd^, 72.86 vs 9.12).

Of four nucleotide biases, only ‘G’ bias was a top ten predictor (three routes/modes), whereas 32 of 128 dinucleotide biases were top ten predictors for at least one route/mode, with six biases ranked in the top ten predictors for >10% of our models: ‘AG pr’ (AG bias in the reverse complement of the sequence, 14.29%), ‘GT’ (13.27%), ‘GA’ (11.22%), ‘GG’ (11.22%), ‘TT’ (10.2%), and ‘GG p3’ (GG bias in position 3-1 within codon reading frames, 10.2%).

Finally, our framework incorporated 190 ‘RSCU’ (Relative Synonymous Codon Usage) features. 44 of 190 ‘RSCU’ features were top ten predictors of at least one route/mode. With ‘CGA’ (CGA bias in predicted ORFs of length ≥ 36, 10.2%) being the only top ten predictor for >10% of our routes/modes.

### Instance-level predictors of transmission routes/modes

Figure 3 visualises the instance-level contributions of top twenty features, by spread of variance in each sub-plot/category, for six categories: main transmission modes (3.A), direct (non-sexual) transmission modes (3.B), direct contact routes (3.C), indirect transmission modes (3.D), arthropod-borne routes (3.E), and environmental transmission routes (3.F). Overall, our virus-host integrated neighbourhoods (MN4D) and hosts similarity features had the most spread of variance across the six categories, whereas the spread of our viral features varied per category. ‘Segmented’ (indicating whether the virus genome is segmented or not) was the viral feature with most spread in variance of contribution to our main modes of transmission (Figure 3.A, Supplementary Dataset 3); ranked 3^rd^ predictor of ‘indirect’ mode of transmission (locally normalised mean absolute SHAP value = 40.74 vs globally normalised = 17.07), but only 18^th^, 42^nd^, and 87^th^ predictor of ‘sexual’, ‘non-sexual’, and ‘vertical’ transmission modes, respectively.

**Figure 3.**
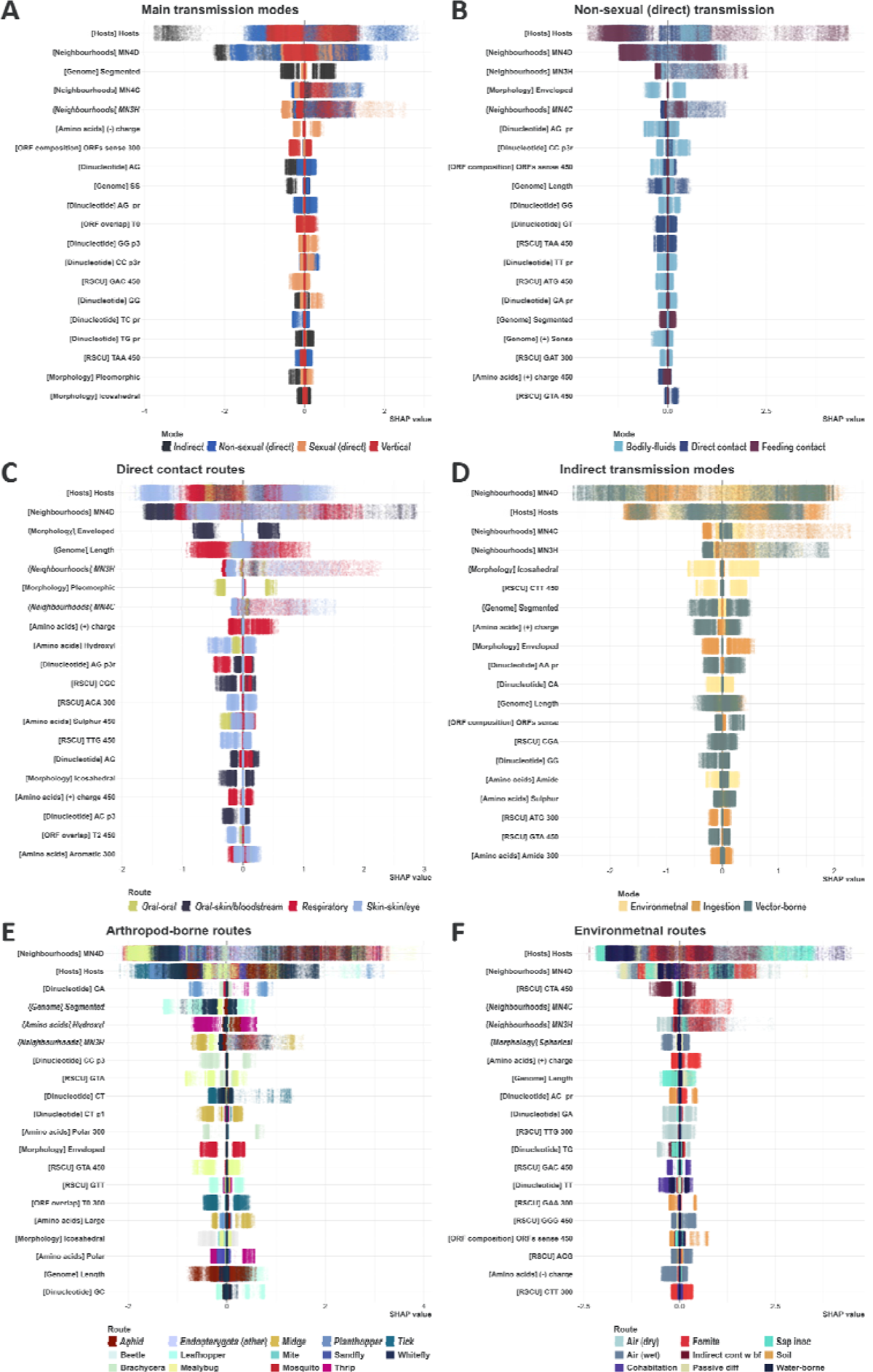
Instance-level feature-contribution to various transmission routes/modes. Instance-level SHAP values quantify the contribution each feature made to a particular (virus-host association) prediction. Here, we averaged instance-level SHAP values generated by all constituent models of each of our top-10 ensembles (50 models per route/mode). In each sub-plot, features were ordered by the spread of their variance (max(variance)-min(variance) across all routes/modes included in each sub-plot), and the top 20 features (from most to least spread) were selected. Points represent virus-host associations (instances) and are coloured by the underlying route/mode. The Y axes represent the selected features (category of each features between brackets). The X axes represent SHAP values. Positive SHAP values indicate that the feature has contributed towards a positive prediction for the virus-host association (the virus species/strain is transmitted to the host species via the given route/mode). Negative SHAP values indicate that the feature has contributed towards a negative prediction (the virus is not transmitted to the host via route/mode). Larger magnitudes indicate that the feature has had a stronger influence on the prediction for the particular instance.

Whether the virus is enveloped or not (‘Enveloped’) had the most spread in variance for both direct transmission modes (Figure 3.B), and direct contact routes (Figure 3.C). It ranked 3^rd^ predictor of ‘bodily-fluids’ transmission mode (local 37.16 vs global 12.82), but only 63^rd^ and 95^th^ predictor of ‘feeding contact, ‘direct contact’ transmission modes, respectively (Figure 3.B). For direct contact (between animals) routes, it ranked 2^nd^ for ‘oral-skin/bloodstream’ transmission route (42.00 vs 20.44) but had virtually no influence on the remaining routes (Figure 3.C).

GA biases was the viral feature with most spread for arthropod-borne transmission routes. It was 3^rd^ most contributing feature to planthopper-borne transmission (45.5 vs 25.1), but only 77^th^ for mealybug-borne and 86^th^ for whitefly-borne transmission (Figure 5.E). ‘Segmented’ also had a large spread in variance of contribution to arthropod-borne routes/modes (Figure 3.E, Supplementary Figures 11.A and 11.D); ranked as 3^rd^ predictor of ‘Auchenorrhyncha-borne’ and ‘leafhopper-borne’ transmission; and 4^th^ of three routes/modes including ‘whitefly-borne’ transmission, but made no contribution to the predictions of 13 to arthropod-borne routes/modes.

Icosahedral structure was the viral feature with most spread in variance in our main indirect transmission modes (Figure 3.D): ranked 3^rd^ predictor of ‘environmental’ transmission (40.31 vs 13.72), but only 27^th^ of ‘ingestion’ and 63^rd^ of ‘vector-borne’ transmission. ‘CTA 450’ bias and ‘Spherical’ morphology were the viral feature with most spread in variance in contribution to environmental transmission routes (Figure 3.F). ‘CTA 450’ bias made significant contribution to transmission via indirect contact with bodily-fluids (ranked 2^nd^), and ‘Spherical’ contributed the most to predictions of ‘air’ and ‘air (wet)’ (ranked 2^nd^), both features made but very small to insignificant contribution to the remainder routes. Supplementary Figures 9-13 visualise the remainder routes/modes not included in Figure 3.

### Prediction of potential routes/modes

We were able to predict (mean top-10 ensemble probability > 0.5) at least one route/mode for 3,108 out of 3,708 (83.82%) virus-host instances without known routes in our dataset (2,004 viruses (8.1.63%) to 1,300 host species (87.84%)) (Figure 4); Of those instances, 2,969 were predicted to be transmitted horizontally (80.07% vs 98.73% of observed associations), and 249 were predicted to be transmitted vertically (6.715% vs 15.15%). Indirect (57.50% vs 81.25%), direct (34.98% vs 41.77%), non-sexual (32.42% vs 41.65%), and ingestion (26.54% vs 26.76%) *modes* were the most predicted after horizontal transmission. The faecal-oral *route* was the most predicted route of transmission for unknown associations (23.33% vs 20.07%), followed by sap inoculation (9.47% vs 19.36% - a plant-only route), vector feeding (8.60% vs 15.81%), and fomite (7.15% vs 23.93%).

**Figure 4.**
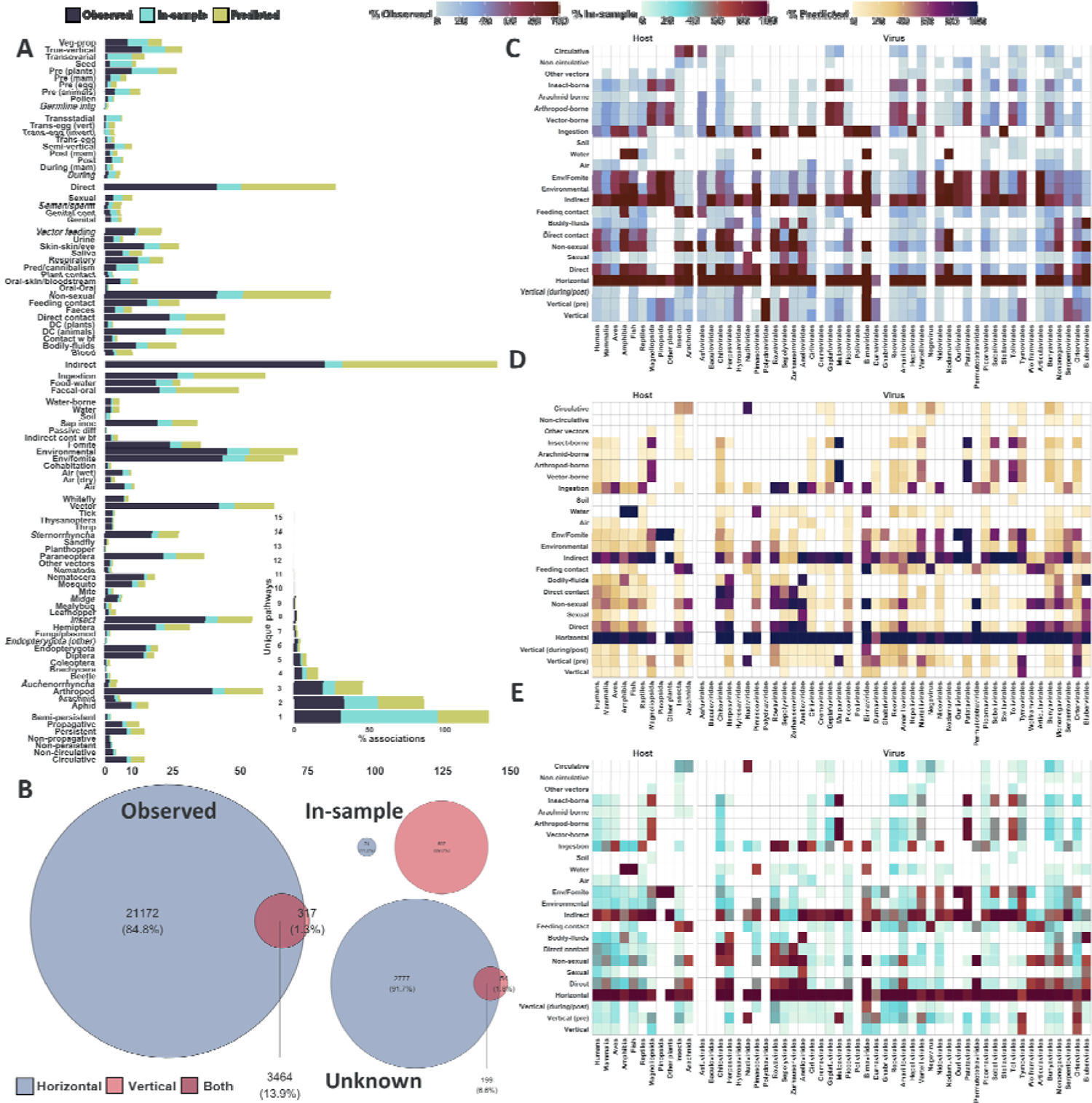
Predicted transmission routes and modes. Panel A – Proportion of predicted unknowns (yellow, n=3,108, no known transmission route, mean probability cut-off>0.5), in-sample predictions (cyan, n=6,701, hitherto unobserved routes predicted with mean probability cut-off>0.5 for associations with at least one observed route) and observed virus-host instances (dark blue, n= 24,953, at least one observed transmission route). Vertical and horizontal modes were removed from the bar plot for better visualisation (represented in panel B as Venn diagrams). The inset (bottom) represent the percent of unique pathways for unknowns and in-sample predictions, as well as observed for each virus-host association. Supplementary Figures 5, 6, and 7 visualise the percent of virus-host associations, whereby the virus is observed, predicted within sample, and predicted for unknowns, to be transmitted to the host via each pair of unique pathways, respectively. **Panel B – Horizontal and vertical transmission.** Venn diagrams represent horizontal transmission (blue) and vector transmission modes (red) for observed, in-sample predictions (with at least one previously observed route, but that is not observed to be transmitted via the corresponding mode), and unknown (out of sample) predictions (instances without observed routes), respectively. **Panel C – Proportion of host-virus instances transmitted by each main route/mode per each host group or virus order.** Rows represent main transmission modes/routes. Columns represent main host groups, and virus orders. Proportions are calculated by the number of instances known to be transmitted via given route/mode per each category (e.g. humans), divided by the total number of instances in the category. Some routes/modes were grouped together for better visualisation (e.g. Vertical (pre), insect-borne). **Panel D – Proportion of host-virus instances, without previously observed transmission route, predicted to be transmitted by each main route/mode per each host group or virus order.** Rows represent main transmission modes/routes. Columns represent main host groups, and virus orders. Proportions are calculated by the number of instances predicted to be transmitted via given route/mode per each category, divided by the total number of instances in the category. **Panel E – Proportion of host-virus instances, with at least one previously observed transmission route, predicted to be transmitted by each main route/mode per each host group or virus order.** Rows represent main transmission modes/routes. Columns represent main host groups, and virus orders. Proportions are calculated by the number of instances predicted to be transmitted via given route/mode per each category, divided by the total number of instances in the category.

Additionally, our models made in-sample predictions (routes/modes predicted with mean top-10 ensemble probability>0.5 for virus-host associations with at least one observed transmission route in our dataset) for 6,701 virus-host associations (26.85% of total associations). The top additional routes/modes predicted in-sample were as follows: true vertical transmission in plants - pre-plants (649 additional associations, 9.685% of total in-sample predictions vs 9.86% of observed and 6.96% of predicted for unknowns); non-sexual (direct) transmission (639, 9.535% vs 41.65% and 32.42%); transovarial transmission (611, 9.12% vs 0.82% and 4.64%); and vertical transmission (597, 8.91% vs 15.15% and 6.715%). Figure 4 visualises both in-sample as well as out-of-sample (unknowns) predicted routes/modes.

Given a virus-host association, we constructed a representative set of *unique transmission pathways* by traversing our hierarchy (Figure 1.A) from routes to root, and including all transmission *routes* of the given virus to the focal host, as well as any transmission *modes* that are not ancestors of any already included routes/modes. We predicted that 1,068 instances have a single unique transmission pathway (29.33% of unknown instances vs 30.89% of associations with at least one observed route and 64.525% for our in-sample predictions); 953 (26.17% vs 33.25% and 22.53%) to have two unique pathways; and 1,020 (28.01% vs 35.855% and 12.94%) to have three or more unique pathways.

### Prediction dependencies

We utilised Mutual Information (MI) to quantify the relationship between predictions (top-10 ensemble mean probability > 0.5) for a given route/mode and those of its sibling route(s)/modes(s) - children of the same parent node in our hierarchy (Figure 1). Figure 5.A visualises the resulting normalised MI estimates. Our normalised MI ranged between 0.00002 (predation/cannibalism) and 0.027 (Brachycera-borne), suggesting a very weak to weak correlation (very limited to limited relationship or dependency) between the predictions of the focal route and those of its siblings. The routes with highest normalised MI were: Brachycera-borne (0.027), air (wet) (0.025), mosquito-borne (0.022), and midge-borne (0.021).

**Figure 5.**
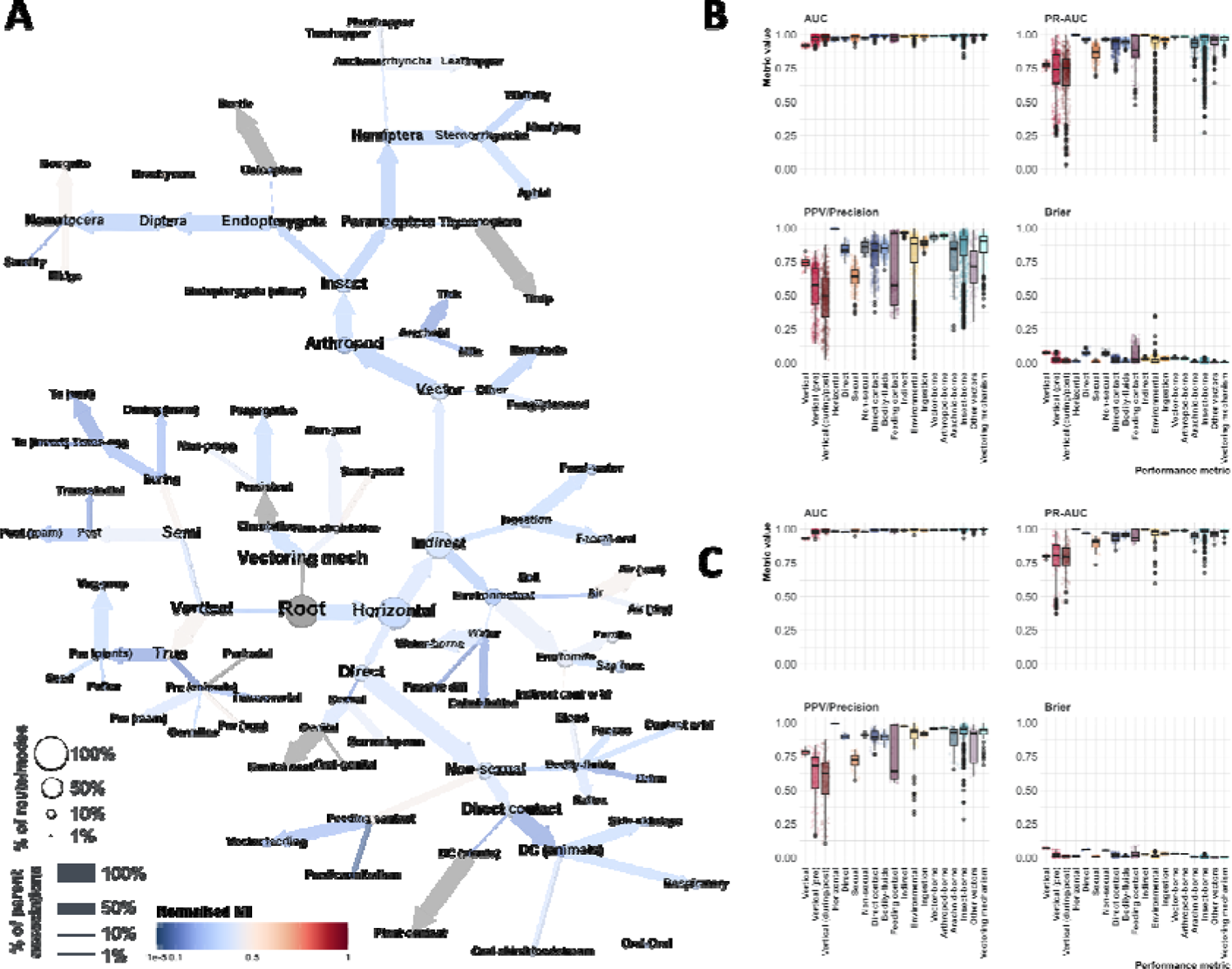
Prediction dependencies and performance assessment. **Panel A – Prediction dependencies**. Nodes represent transmission routes/modes modelled in this study. Edges indicate our hierarchy (Figure 1). Nodes are coloured by Normalised Mutual Information (MI) estimates between the mean probabilities (derived from our top-10 ensembles) of instances predicted by the route/mode represented by the node (mean probability > 0.5), and corresponding instance-wise probabilities of their siblings (nodes with the same parent node). Grey nodes indicate structural nodes (not transmission related), and only children (routes without modelled siblings). Nodes sizes and thickness of edges are same as Figure 1. MI quantifies how much knowing the value of one variable can tell us about the value of the other variable. If the resulting estimate is high, it indicates a strong relationship or dependency between the variables, meaning that knowing one variable provides useful information about the other. To assess the statistical significance of the MI estimates, we compared each estimate to a null distribution using bootstrapping (n=2,000). Supplementary Figure 13 visualises the resulting p-values. Supplementary Figures 14-16 visualises decencies between probabilities for routes/modes and knowledge of their siblings, predictions of their siblings, as well as knowledge of routes/modes and resulting probabilities of their siblings, respectively. **Panel B – Performance assessment of constituent class-balancing ensembles of against 50 held-out test sets at >0.5 probability threshold.** Points represent the class-balancing ensemble mean values for each performance metric (50 points per route/mode, 98 routes/modes in total). Supplementary Figures 20 and 22 provides results all included metrics/constituent models. **Panel C – Performance assessment of constituent class-balancing ensembles of our top-10 selected ensembles against 10 held-out test sets at >0.5 probability threshold.** Points represent the class-balancing ensemble mean values (5 models per each run) for each performance metric (20 points per route/mode, 98 routes/modes in total). Supplementary Figures 21 and 23 provides results all included metrics/constituent models. In panels B and C, Boxplots represent the interquartile range (IQR), of the data distribution per each category of transmission route/mode. Horizontal lines within the box represent the median of the data distribution. Whiskers extend from the edges of the box to the minimum and maximum values within a distance of 1.5 times the IQR from the nearest quartile, individual data points that fall outside the range covered by the whiskers are plotted as outliers. Supplementary Table 9 provides full definitions of included performance metrics. For Brier score values closer to 0 indicate better performance, and those closer to 1 indicate worse performance.

### Model performance

We evaluated the performance of our predictive framework against held-out test sets (see methods and Supplementary Note 7 for full details). Overall our framework achieved average ROC-AUC = 0.988±0.017 (values closer to 1 indicates high accuracy in distinguishing between positive and negative instances); and average F1-score (>0.5 probability cut-off, commonly used for imbalanced classes, not used in tuning or selection of models) = 0.806±0.169 (values closer to 1 indicates high accuracy in identifying positive instances while minimising potentially false positives).

We ranked our class-balancing ensembles (n=50, see methods), trained for each pathway (n=98), based on the average of four metrics: AUC, PR-AUC, Precision, and 1-Brier score, we selected the top-10 performing ensembles to produce final predictions. Our top-10 ensembles achieved ROC-AUC=0.991±0.012, and F1-score=0.855±0.143. Figures 5.B and 5.C visualise the performance of constituent models over 50 iterations, as well as for constituent models of our top-10 performing ensembles against our selection metrics, respectively. Table 1 lists the average performance against ten metrics. Supplementary Figures 20-23 visualise the performance of constituent ensembles/models against all ten metrics. Supplementary Figure 24 illustrates our post-hoc assessment of in-sample predictions of our top-10 selected ensembles of five class balancing techniques at >0.5 mean probability threshold. Supplementary Dataset 4 lists average performance metrics per each modelled route/mode for both all class-balancing ensembles, as well as our top-10 ensembles.

**Table 1.** Average performance and standard deviation of all ensembles, and top-10 selected ensembles, across all routes/modes. Brier scores range from 0 (best performance) to 1 (worst performance); and MCC values range from +1 (best performance) to −1 (worst performance). Values in square brackets indicate worst and best performing ensembles, respectively. Top-10 ensembles were selected based in the average of four metrics (AUC, PR-AUC, PPV/Precision and Brier (1-actual score)). Supplementary Figures 20-23 visualise performance metrics of constituent models. Supplementary Figure 24 provides post-hoc assessment of in-sample predictions of our top-10 selected ensembles. Supplementary Dataset 4 lists performance assessment results for each route/mode.

|  | All Ensembles (n=50) | Top-10 Ensembles |
| --- | --- | --- |
| **ROC-AUC** | 0.988±0.017 [Vertical = 0.917 (±0.011), Cohabitation = 0.999 (± 0.0004)] | 0.991±0.012 [Vertical = 0.931 (±0.008), Cohabitation = 0.999 (± 0.0001)] |
| **PR-AUC** | 0.891±0.147 [Trans-egg (invert)= 0.332 (±0.167), Horizontal= 0.999 (±0.0002)] | 0.922±0.109 [Transovarial= 0.446 (±0.052), Horizontal= 0.999 (±0.00004)] |
| **F1-score** | 0.806±0.169 [Trans-egg (invert)= 0.237 (±0.086), Horizontal= 0.994 (±0.001)] | 0.855±0.143 [Trans-egg (invert)= 0.293 (±0.073), Horizontal= 0.995 (±0.0007)] |
| **NPV** | 0.987±0.048 [Horizontal= 0.546 (±0.091), Mealybug-borne= 0.999 (± 0.0002)] | 0.987±0.042 [Horizontal= 0.601 (±0.059), Passive diffusion= 0.999 (± 0.0002)] |
| **PPV/**<br>**Precision** | 0.755±0.206 [Trans-egg (invert)= 0.149 (±0.06), Horizontal= 0.995 (±0.001)] | 0.829±0.179 [Trans-egg (invert)= 0.180 (±0.049), Horizontal= 0.996 (±0.0008)] |
| **Recall/**<br>**Sensitivity** | 0.896±0.105 [Seed= 0.605 (±0.072), Vector feeding= 0.995 (±0.004)] | 0.905±0.09 [Seed= 0.615 (±0.089), Vector feeding= 0.995 (±0.003)] |
| **Specificity** | 0.976±0.052 [Horizontal= 0.656 (±0.066); Sandfly-borne= 0.999 (±0.001)] | 0.984±0.037 [Horizontal= 0.672 (±0.061); Sandfly-borne= 0.999 (±0.001)] |
| **MCC** | 0.797±0.156 [Trans-egg (invert)= 0.305 (±0.101), Vector feeding= 0.982 (±0.007)] | 0.847±0.133 [Trans-egg (invert)= 0.384 (±0.079), Vector feeding= 0.990 (±0.0035)] |
| **Brier** | 0.018±0.024 [Feeding contact= 0.144 (±0.047), Sandfly-borne= 0.002 (±0.001)] | 0.014±0.016 [Vertical= 0.070 (±0.005), Passive diffusion= 0.001 (±0.0003)] |
| **TSS** | 0.872±0.111 [Seed= 0.578 (±0.069), Vector feeding= 0.992 (±0.004)] | 0.889±0.096 [Seed= 0.593 (±0.083), Vector feeding= 0.994 (±0.004)] |

## DISCUSSION

In this study, we constructed a computational framework that explored the landscape of viral transmission in the animal and plant kingdoms, with the aim to firstly uncover the specific viral features and evolutionary signatures predictive of the transmission routes viruses utilise between their respective hosts; secondly to assess the applicability of predictive approaches as means to triage the potential transmission routes of emerging viruses; and finally to quantify possible gaps in our knowledge of transmission pathways of existing viruses to their known hosts.

This was achieved by training lightGBM ensembles on a comprehensive dataset of transmission routes and modes of 4,446 viruses to 5,317 animal and plant species (Figure 1). Broadly, 112 of our 442 viral features were important predictors for at least one of 98 routes and modes of transmission analysed (Figures 2). Furthermore, analysing the differences in contribution our viral features made to individual predictions of hierarchically close routes and modes (Figure 3), enabled us to establish the different roles the same, or similar, viral features and evolutionary signatures play in influencing viral transmission dynamics. We further quantified the ability of our ensembles to discriminate between closely related routes/modes by examining dependencies of their predictions (Figure 5.A), and found that overall, our ensembles exhibited very limited to limited dependency between related routes/modes.

Following assessment of the predictive performance of our ensembles using a complementary array of metrics (Table 1, Supplementary Data 4), whereby our ensembles achieved high level of performance for routes commonly associated with high consequence human, animal, and plant viruses (e.g. vector-borne pathways, respiratory viruses), we applied our ensembles to predict the transmission routes of 2,004 virus species or strains for which there are no known transmission routes to 1,300 host species. Our models predicted at least one route/mode for ∼84% of those instances (Figure 4).

Furthermore, we identified an additional 19,396 transmission routes/modes potentially unobserved in 2,147 viruses and 2,355 host species with at least one route/mode observed (Figure 4). These predictions were made across a total of 4,076 animal and plant viruses.

This study, therefore, showcases the potential to provide early insights into the epidemiology of a newly emerging virus, and hence can be used to facilitate rapid response and significantly triage the time-consuming investigations to confirm the routes.

### Application of multiple perspectives in predicting transmission routes

We generated predictive features from three complementary perspectives: viruses, hosts, and our virus-host integrated neighbourhoods which depict the topology of the virus-host network in the phylogenetic neighbourhood [29] of a virus.

Our ensembles indicated that our integrated neighbourhoods and host similarity features were highly predictive of all transmission routes/modes (Figure 2). However, all of our viral features were also predictive, or improved the predictions, of at least one route/mode, enabling us to distinguish between closely related routes/modes. This highlights the advantage of our multi-perspective approach to investigating mechanisms of virus transmission, and further emphasises the applicability of multi-perspective approaches, in line with the significant promise they have shown in predicting virus-host associations [1,2,30,31].

Other existing approaches which aim to predict viral phenotypes solely from one perspective – such as the viral sequence – will miss key features from the host and network perspectives which would enhance their predictive performance. For example, our integrated neighbourhoods and host features were the most informative predictors across all routes/modes, providing the large-scale structure of the viral transmission landscape. Our host similarity was top-10 predictor of 97 routes (ranked 12^th^ for nematode-borne transmission).

Our 442 viral features (often referred to as viral evolutionary signatures [29]) further enhanced accuracy and explainability at higher resolutions of the individual association and route/mode levels, and therefore improving distinction between similar sister routes/modes. For instance, envelope status was an important predictor of mosquito-borne transmission (ranked 3^rd^) but has no effect on predicting sister routes midge- and sandfly-borne transmission. Conversely, ‘CT p1’ bias (refer to Supplementary Note 3 and Supplementary Table 5 for full definition) was predictive of midge-borne transmission (ranked 4^th^) but has no effect on predicting mosquito-borne transmission, and negligible effect on predicting sandfly-borne transmission. This is mirrored throughout our hierarchy, including routes that are often interlinked: ‘(+) charge 450’ bias ranked 4^th^ predictor of faecal-oral transmission, but only 9^th^ for transmission via ingestion of food/water. Furthermore, and in line with previous studies[27], envelop status was a top predictor (ranked 5^th^) of faecal-oral transmission, but has no effect on predicting transmission via ingestion of food/water (Supplementary Figure 11.B). Indeed, our viral features can differentiate between mechanistically similar routes, for instance, ‘spherical’ morphology was highly predictive (ranked 3^rd^) of transmission via inhalation of wet environmental particles (air (wet)) but had little effect on predicting respiratory transmission (ranked 110^th^).

### Framework deployment

We next deployed our trained ensembles to predict transmission mechanisms for viruses with no observed routes to their known hosts (3,108 virus-host associations). Proportionally, direct transmission routes were overall more likely to be predicted than indirect (Figure 4). However, ingestion and faecal-oral were the most likely unobserved individual routes to be predicted. We also noted a significant underestimation in the number of plant viruses potentially transmitted by insect.

Looking specifically at predictions for arthropod-borne transmission, there are 608 associations (16% of those without observed routes in our dataset). Of these 24% are to animals, and 76% to plants. The greatest proportion is within the Hemiptera-borne routes (354 associations); the Hemiptera being a superorder of insects which contains the majority of the plant vectors [32]. Given the relative difficulty, time, and expense of demonstrating arthropod-borne transmission, which requires vector competence studies [33], it is not surprising that many such routes remain undetermined. Our approach could readily be used to triage the vast numbers of potential competence studies into high-likelihood and high-priority combinations.

### Strength of predictions in potentially high-consequence transmission routes

Routes of transmission are fundamentally intertwined with virus ecology and epidemiology [10]; determining the virus spread within and between host species and populations, therefore having a large effect on the rate of spread (R_0_) and, for some routes, the geographical range of outbreaks [34,35]. Respiratory (in humans) and vector-borne (in animal and plants) transmission frequently results in high-consequence outbreaks[12,16,36], and our framework is particularly effective at identifying viruses with these two mechanisms. For example, for respiratory viruses, our top-10 selected ensembles approach achieved mean ROC-AUC=0.990, and mean F1-Score = 0.864; whereas our arthropod-borne classifiers (n=26, including animal and plant viruses) averaged ROC-AUC=0.997, and F1-score = 0.921 (Supplementary Dataset 4). Furthermore, at the level of individual vector-borne routes, our top-10 ensembles exhibited high predictive performance for all important routes of both plants (e.g. aphid-borne (ROC-AUC = 0.998, F1-score = 0.951), whitefly-borne (0.9997, 0.987), thrip-borne (0.999, 0.976)), and animal (e.g. mosquito-borne (0.999, 0.959), tick-borne (0.995, 0.944)) viruses (Supplementary Dataset 4). This strength is likely driven by the large amount of data (e.g. 40% of our virus-host associations had at least one arthropod-borne route of transmission), and the large number of human-virus data assigned to respiratory transmission (33% of virus-human associations). Given the importance of these two classes from previous high-consequence outbreaks, the high density of data serves to improve our pipeline’s predictive performance where it is needed most.

### Limitations

We acknowledge certain methodological limitations and shortcomings in our study. Firstly, in order to synthesise of meaningful features, the training of our framework has been restricted to fully sequenced viruses with at least one known animal or plant host species. While there are no theoretical limitations to deployment of our trained models, our assessment of our framework’s predictive performance cannot be extended to partially sequenced viruses. Our framework could be utilised to predict potential transmission routes of viruses without known hosts, to probable hosts as long as diversion times of those probable hosts, relative to other host species in our framework, are known.

Secondly, we could not integrate the full genome sequence of host species, as those data are lacking for the majority of species included. Similarly, we could not include life-history or other ecological traits of our hosts, as those data are not available for many species, and are not comparable between animals and plants. Thus, we had to rely on diversion times as the only proxy to differentiate between our hosts.

Finally, our method does not make assumptions, or use features, based on which specific parts of the virus genome, or which receptor or receptor binding proteins are commonly utilised in specific transmission routes. Instead, we synthesised a wide range of features (n=446) from three complementary perspectives. This ‘no-preconceptions’ approach enables us to analyse transmission routes/modes of viruses to their known hosts without being restricted by our current, and highly incomplete, knowledge of the specific biological and molecular mechanisms which govern mechanism of transmission [1,2]. Whilst some of these details are known for a very limited number of well-studied viruses and hosts, they are unknown for the vast majority. Therefore, a machine learning study aiming for breadth of understanding across all transmission routes, viruses, and hosts cannot use these incomplete data. Despite this ‘no-preconceptions’ approach having this distinct advantage, it is also a limitation of the predictions, and may result in less accurate predictions for those well studied transmission routes/viruses/hosts for which important factors are well known.

## Conclusions

This study is the most taxonomically broad study of its kind, and is the first to demonstrate that viral sequence, morphology, and host information, increasingly available in the first few days of an outbreak, can be used to accurately identify the transmission routes of a novel virus, across the animal and plant viromes. Importantly, we have showcased that predictions can be achieved with high accuracy, including for respiratory and vector-borne routes/modes, which together encompass the majority of high-consequence outbreaks across animals and plants. Together with the more matured field of viral host-range prediction, much of the key information which is needed to assess the potential for a virus to cause a high-consequence outbreak can be predicted in the first few days, enabling rapid and targeted mitigation procedures and triage of the time-consuming confirmatory investigative.

## METHODS

### Data sources and unification

#### Viruses

We downloaded complete and references virus sequences from Genbank [37]. Sequences without known vertebrate or plant host were excluded, and those labelled with the terms: ‘vector’, ‘construct’, ‘vaccine’, or ‘clone’ were removed, as they are mainly laboratory-derived and/or manipulated. The number of ambiguous bases was identified for each sequence, and those requiring more than 1,024 permutations to resolve were excluded. Segmented viruses were included only if all corresponding individual segments met these criteria. This resulted in a total of 6,803 virus species or strains that were included in further analyses. Supplementary Table 1 illustrates the distribution of these viruses.

#### Virus-host associations

We compiled a comprehensive set of virus-host associations from relevant databases [38–42] and literature (e.g. [43]). Those sources were manually verified for accuracy, and any association where the underlying evidence (e.g. publication) only concurrently cite the virus and host, or specifically indicate an absence of interaction, were removed. This resulted in 28,661 associations between the above viruses and 5,750 host species (animals = 3,649, and plants = 2,101). Supplementary Table 2 lists the distribution of these associations by virus Baltimore classification and host taxa. Supplementary Dataset 1 lists all included associations and their sources.

#### Transmission routes

We identified 81 <u>non-mutually-exclusive</u> routes of virus transmission, in animals and plants, by searching relevant literature (Supplementary Table 3). The breakdown of these routes was as follows: vertical (14 routes), sexual (3), transmission via direct contact with bodily-fluids (5), feeding contact (2), direct contact (5), ingestion (3), indirect contact (2, minor routes), environmental transmission (10), arachnid-borne (2), insect-borne (28), and transmission via other-vectors (3).

We adopted a two-fold strategy to search the literature for whether our viruses are known to be transmissible (to their hosts) by one or more of our routes as follows: Firstly, we identified Title and Abstract (TIABs) of PubMed papers linked to single virus species (i.e. excluding TIABs with multiple viruses), and subsequently matched each TIAB, via keyword searches, to the transmission routes described above. The resulting routes/TIABS matches were verified, and erroneous associations removed. Secondly, we manually captured routes of transmission of viruses for which no papers were identified by the previous step, as well as for routes not detected in the TIABs, by searching through textbooks and virus sources (e.g. [43–46], Supplementary Dataset 2 lists all sources).

Following a further manual check for accuracy, and to remove erroneous routes, we were able to identify at least one transmission *route* (of total = 77 routes, Supplementary Table 3) for 4,446 viruses (65.35% of total – Supplementary Table 4) to 5,317 hosts. Overall, we identified a total of 24,953 virus-host associations. Supplementary Dataset 2 lists identified routes and their sources for all associations.

#### Hierarchy construction

Given a virus-host association, we considered the virus to be transmitted to the host via a parent mode (e.g. dengue virus is insect-borne to humans), if we found it to be transmissible by at least one route that is also a child node of the parent node (e.g. dengue virus is mosquito-borne to humans) in our hierarchy (Figure 1).

### Predictive features

We engineered 446 features, in three complementary perspectives, as follows (For full description see Supplementary Notes 3-5).

#### Viral features (Supplementary Note 3)

In order to facilitate the identification of the unique evolutionary signatures associated with specific transmission routes/modes, we synthesised 442 features from the virus genome (Supplementary Table 5), categorised as:

a. Basic genomic features (9 total): including length, GC content, and genomic structure (e.g. segmentation, linearity, and sense). These features proxy virus stability, replication mechanisms, tropism, and evasion strategies, which may determine the range of transmission routes/modes deployed by the virus.
b. Genome biases: including nucleotide (4 features), dinucleotide (128 features), and amino acid biases (57 features), as well as Relative Synonymous Codon Usage[47] (RSCU, 204 features). These features may influence the routes of virus transmission through their impact on viral fitness and host interactions. Additionally, they reflect underlying selective pressures acting on the virus during transmission, replication, and co-adaptation processes.
c. Open Reading Frames (ORF) specific features: including ORF composition (12 features), genome coverage (6 features) and proportion of total ORF length with overlaps (21 features). Those features proxy the genetic architecture, complexity, and functional potential of the virus.
d. Morphological (capsid) features (16) may influence how the virus interacts with environment, host cells, intracellular compartments, tissues, and bodily fluids, ultimately impacting the transmission routes/modes it may utilise.
e. Replication site (2 features, cytoplasm/nucleus) may influence the range and diversity of transmission routes exploited by a given virus.

#### Host similarity

In order to parameterise transmission routes/modes that are closely interlinked with host taxonomy, as well as those restricted to certain taxa (e.g. plant-only or mammalian-only routes), we obtained a time tree of 4,342 plant and animal species from the Time Tree of Life[48] (timetree.org), and computed diversion time distance between 99.98% of all included host species pairs (Supplementary Note 4). We utilised the resulting distances to calculate similarity between the host species of the focal virus-host association, and all hosts (excluding the focal hosts), for which at least one virus is known to be transmitted by the focal route/mode.

#### Virus-host integrated neighbourhoods (Supplementary Note 5)

Given that closely related viruses may utilise similar transmission routes in taxonomically close hosts (e.g. majority of orthoflaviviruses are mosquito-borne in mammals and birds), as well as a diverse set in taxonomically distant hosts (e.g. orthoflaviviruses may exploit sexual and vertical routes in some of their vertebrate hosts, as well as some of their arthropod vectors), we incorporated pair-wise association-level similarities in our predictive pipeline.

This was achieved by expanding the concept of phylogenetic neighbourhoods [29], so that for any given virus-host association and a transmission route/mode, we firstly identified the set of viruses most closely related to the focal virus, that are known to be transmitted to some of their hosts via the focal route/mode – the virus phylogenetic neighbourhood (PN) [29]. We then included the hosts their viruses infect via the focal route/mode to compute three complementary features (Supplementary Figure 1):

1. *MN3H* indicates whether the focal host is susceptible, via the given route/mode, to viruses which exhibit high sequence similarity to the focal virus.
2. *MN4D* measures the average similarity between the focal association, and associations between PN viruses, not known to be transmissible to the focal host via the given route/mode, and hosts other than the focal host.
3. *MN4C* measures the average similarity of the focal association to associations between PN viruses, known to be transmissible to the focal host via the given route/mode, and hosts other than the focal host.

### Basic components of the predictive framework

#### Binary Relevance multi-label classification

As the same virus species/strain may deploy multiple routes/modes in the same host species (Supplementary Figure 6), we developed a binary relevance multi-label classification framework to enable prediction of transmission routes/modes at the virus-host instance level. Unlike multi-class approaches, multi-label classification allows each virus-host instance to belong to multiple categories or classes (i.e. routes/modes), rather than just one. Binary relevance [49], where each label is treated as an independent binary classification problem, is one of the simplest, best-performing, and most commonly implemented, approaches to multi-label classification.

We adopted this approach for three reasons: 1) Binary relevance models each label independently, thus simplifying the classification task into a suite of binary decisions that can be trained and run in parallel. This modularity makes binary relevance highly scalable and computationally efficient. 2) Importantly, the independent nature of the training process, allows us to accommodate imbalanced input datasets of varying label frequencies, as is the case with our transmission routes/modes (see below). 3) Binary relevance is highly interpretable, as predictions for each label are independent, thus allowing us to interpret predictions, and compare the contributions made by our input features, for individual routes/modes.

#### LightGBM

We trained a suite of LightGBM (Lightweight Gradient Boosting Machines)[50] models per every route/mode sufficient data (n=98, 57 routes, 41 modes). LightGBM utilises boosting and gradient descent to build decision trees (weak learners) sequentially, based on the error obtained from previous iterations. The gradient-based approach allows LightGBM to handle large-scale datasets efficiently, minimising memory usage and acceleration training speed. Tree-based learning enables it to capture complex patterns in the data effectively. Unlike other implementations (e.g., XGBoost) which grow trees in level-wise manner, LightGBM implements a leaf-wise approach to tree growth, which enhances training and prediction speed.

#### Class balancing

The fraction of observed virus-host instances varied greatly per route/mode (Figure 1, Supplementary Figure 3), ranging from 0.16% (vertical trans-egg transmission in invertebrates) to 98.71% (horizontal transmission) of the 24,953 virus-host associations with at least one observed transmission route/mode. This presented a varied and bi-directional imbalance between observed (positive class), and unknown (negative class) transmission routes/modes for our associations.

We compared the performance of 22 class-balancing techniques (Supplementary Table 6), as well as that obtained by tunning a lightGBM specific hyperparameter used to address class imbalance in binary classification tasks, across all modelled transmission routes/modes (n=98), over a single iteration of our pipeline (below, steps 1-5). Following an assessment against the corresponding held-out test-set (stratified, 10%), using a comprehensive set of performance metrics (Supplementary Table 8, Supplementary Figure 4), we incorporated the following five class balancing into our multi-label classification framework:

1. Two over-sampling techniques - SL-SMOTE (25%, minority class = 25% of resulting total), and MWMOTE (25%). Safe-Level-Synthetic Minority Over-Sampling (SL-SMOTE) [51] synthesises new minority class instances from existing cases in a manner that preserves the “safe” minority instances with a higher nearest neighbour density; by focusing on the safe minority instances and generating synthetic samples around them, SL-SMOTE helps prevent overfitting by avoiding the creation of outliers or noise in the minority class distribution. Majority Weighted Minority Over-Sampling Technique (MWMOTE) [52] generates synthetic minority class samples based on the weighted majority instances in their vicinity using a hierarchical clustering approach. This targeted approach ensures that synthetic samples are generated in regions where the minority class is underrepresented but surrounded by a majority of instances. MWMOTE is less susceptible to the effects of noisy instances than SMOTE and its extensions.
2. One over-sampling and noise reduction hybrid technique - SMOTE (NRAS, minority class = 50% of resulting total). Noise Reduction A Priori Synthetic Over-Sampling (NRAS) [53] is applied to “clean” training data prior to synthesising new instances using SMOTE (Synthetic Minority Over-Sampling Technique [54]). This hybrid approach synthesises instances for the minority class while reducing noise through the removal of majority class instances that are likely to be misclassified. It combines the principles of SMOTE with noise reduction strategies to create a more balanced and noise-resistant dataset for improved classification performance.
3. Two over- and under-sampling hybrid techniques - SMOTE-ENN (25%), and SMOTE-TL (25%).

In SMOTE-ENN (SMOTE [54] followed by Edited Nearest Neighbours (ENN) [55]), minority instances are first synthesised using SMOTE, and then ENN iteratively removes instances whose class label differs from the majority class label of its nearest neighbours from the training set, thus creating a smoother decision surface. Whereas in SMOTE [54] followed by Tomek Links[56] (SMOTE-TL), the borderline instance resulting from the application of SMOTE are removed using Tomek Links to enhance the separation between classes in the resulting training set, thus allowing for the retention of informative samples while still balancing the class distribution.

### Predictive framework workflow

#### Training, optimisation, and validation of class-balancing ensembles

In order to accommodate the above selection of class-balancing methods, and to incorporate the uncertainty arising from the stochastic elements in machine-learning, as well as from the variations in class-balancing resampling techniques and resulting training sets, we carried model training and optimisation over 50 replicate class-balancing ensembles per each route/mode (comprising five LightGBM models each, corresponding to the selected class-balancing methods). Our framework encompassed the following steps (Supplementary Figure 4):

<u>Step 1 – Initialisation.</u> We first initialised an input set of virus-host associations for each of our routes/modes comprising all associations whereby the virus is known to be transmitted to the host species via the given route/mode (positive class), and the remainder of our 24,953 virus-host associations with at least one observed transmission route/mode after excluding all positive-class associations (negative class).

<u>Step 2 – Splitting</u>. We split the above set into training (80%), optimisation (10%), and validation (held-out test, 10%) sets per each iteration (n = 50). Sets were drawn using stratified random sampling to maintain class ratio in the resulting splits.

<u>Step 3 – Generating association derived features.</u> We recomputed association-derived feature (host similarity, virus-host integrated neighbourhoods) for each set as follows: 1) training - only associations within the training set (∼80% of total) were used; 2) optimisation - associations within both training and optimisation sets were used (∼90%); and 3) validation - all available associations were used.

<u>Step 4 – Class balancing.</u> We passed each training set through our class balancing pipeline, thus generating five training sets, as described above (and Supplementary Note 6). Integer and binary variables were adjusted, post balancing, by rounding.

<u>Step 5 – Optimisation.</u> A LightGBM model was then optimised for each of the resulting balanced training sets (n=5) against the corresponding optimisation set (10%, not balanced, same set was used for all five models). We performed Bayesian (model-based) optimisation, using two metrics: AUC and PRAUC, against the optimisation set, to tune nine hyperparameters (Supplementary Table 7). We set the tuning stopping to stagnation over 15 rounds. Optimisation was performed using the R packages: mlr3tuning and mlrintermbo.

<u>Step 6 – Bagging of class-balancing techniques.</u> We generated predictions by averaging probability outputs of the five constituent models (i.e. bagging). This approach allowed us to factor in uncertainties arising from our class-balancing processes.

<u>Step 7 – Validation.</u> Performance of resulting bagged ensembles and their constituent models was assessed against the stratified held-out test sets (10% of virus-host interactions per iteration, not balanced), using a comprehensive set of performance metrics (Supplementary Table 8). Steps 2-7 above were performed using the R Packages: mlr3, mlr3extralearners, mlr3tuning, mlrintermbo, and lightgbm.

#### Construction of final ensembles

We ranked the class-balancing ensembles (n=50), trained for each route/mode (n=98), based on the average of four metrics: AUC, PR-AUC, Precision, and 1-Brier score, computed against the corresponding held-out test sets. We then excluded underperforming models by averaging predictions from the top-10 class-balancing ensembles to generate finale predictions.

A value closer to 1 of the first three metrics indicates a better performance. The Area Under the ROC Curve (AUC) is widely used to evaluate the discrimination ability of binary classifiers. For imbalanced datasets, Precision-Recall AUC (PR-AUC) is preferred, as it focuses on the positive class and is more sensitive to class imbalances. Precision (Positive Predictive Value) further assesses the accuracy of positive predictions. Conversely, Brier scores closer to zero indicate more reliable predictions. Brier score measures the accuracy of probabilities generated by the models, thus assessing the confidence of a model. A more confident model would generate probabilities closer to 1 for instances of the positive class, and closer to 0 for instances of the negative class.

By employing a combination of these metrics, we could comprehensively assess the performance of our ensembles, factoring in discrimination, class imbalances, positive prediction accuracy, and calibration aspects for a more informed decision-making.

#### Model interpretability

We quantified the contribution our features made to local (instance-level) predictions using SHAP (SHapley Additive exPlanations) values [57]. Instance-level SHAP values employs cooperative game theory principles to quantify the marginal contribution of each feature to the divergence between the actual prediction and the average prediction across all instances. We first computed local SHAP values for each constituent model (n=50) of our top-10 ensembles, and then averaged them to generate an aggregate SHAP value for each instance/feature combination. Instance-level SHAP values can be either positive or negative. A positive SHAP value indicates that the corresponding feature increases the resulting prediction (probability) compared to the average prediction across all possible instances. Conversely, a negative SHAP value suggests the feature has a suppressive or inhibitory effect on the prediction. The absolute magnitude of the SHAP value for a feature measures the importance or influence the corresponding feature has on the prediction for a specific instance.

To facilitate the comparison of the contribution our features made to the predictions, and to assess their overall importance, across all modelled routes/modes, we normalised SHAP values at two levels as follows:

<u>Local (locally normalised) SHAPs</u> - Provides relative importance within each route/mode and allows us to compare the relative importance of features within that specific route/mode. This is turn enable us to understand the contribution each of our feature made to the prediction of each modelled route/mode. Additionally, local normalisation facilitates model-specific interpretation, such as feature importance within the context of each individual route/mode, thus enhancing interpretability of our predictions.

<u>Global (Globally normalised) SHAPs</u> - Provides relative importance across all models, which allows us to compare the relative importance of features across different routes/modes, which in turn enables us to identify consistent patterns of feature importance across multiple routes/modes. Additionally, normalising globally simplifies the comparison process by aligning the scales of SHAP values across all routes/modes. Thus, allowing for straightforward comparisons without the need to consider individual model characteristics.

Additionally, we examined the stability of our SHAP values of our top-10 ensembles (Supplementary Figure 17), as well as in relation to correlation between our predictive features (Supplementary Figure 18). Supplementary Results 4 provides further details.

#### Prediction dependencies

To maximise the utilisation of our models in mitigating against future emerging viruses, it was essential to measure their ability to differentiate between closely related routes/modes (e.g. mosquito-borne vs midge- or sandfly-borne). To this end, we evaluated the extent by which our predictions for a given route/mode depended on predictions of its sibling route(s)/modes(s) - children of the same parent node in our hierarchy (Figure 1). This was achieved by utilising Mutual Information (MI) to determine if there exists a dependency between the mean probabilities (top-10 ensembles) of instances predicted to be transmitted by the focal route/mode (mean probability >0.5), and the corresponding mean probabilities of its siblings. Where the focal route/mode has more than one sibling (e.g. mosquito-borne has two siblings: midge-borne and sandfly-borne), we used the maximum mean probability for each instance in our calculations.

To assess the statistical significance of the MI estimates, we compared each estimate, obtained using mi.empirical function in the R package entropy, to a null distribution using bootstrapping (n=2,000). The null hypothesis assumes there is no dependency between the predictions of focal route and those of its siblings. The resulting p-value, calculated using empPvals function in the R package qvalue, indicates the probability of obtaining an MI value as extreme or more extreme than the observed value, assuming the null hypothesis is true.

We normalised MI estimates by dividing each by the maximum possible MI given the underlying sample size. Normalised MI ranges between 0 and, where 0 indicate there is no information shared between the focal route/mode and its siblings, and therefore there is no relationship between the two; whereas a Normalised MI ≥ 0.7 suggests a strong correlation between the focal route/mode and its siblings, and changes in one are likely to be reflected in the other.

## DATA AND CODE

All data and codes accompanying the manuscript can be found at: https://doi.org/10.6084/m9.figshare.24581421.v1.

## ACKNOWLEDGMENTS

MW acknowledges support from BBSRC and MRC for the National Productivity Investment Fund (NPIF) fellowship (MR/R024898/1) which funded this research. MSCB, MB, and MW acknowledge BBSRC and NERC grants BB/W00402X/1 and NE/W002302/1 which funded parts of this research.

## AUTHOR CONTRIBUTIONS

Conceptualisation: MW, MSCB; Data Curation: AK, MH, MW; Formal Analysis: MW, MSCB, JP; Funding Acquisition: MW, MB, MSCB; Investigation: MW, MSCB; Methodology: MW, MSCB; Writing – Original Draft Preparation: MW, MSCB; Writing – Review & Editing: All authors

## REFERENCES

1. Wardeh M, Blagrove MSC, Sharkey KJ, Baylis M. Divide-and-conquer: machine-learning integrates mammalian and viral traits with network features to predict virus-mammal associations. Nat Commun 2021 121. 2021;12: 1–15. doi:10.1038/s41467-021-24085-w

2. Wardeh M, Baylis M, Blagrove MSC. Predicting mammalian hosts in which novel coronaviruses can be generated. Nat Commun 2021 121. 2021;12: 1–12. doi:10.1038/s41467-021-21034-5

3. Becker DJ, Albery GF, Sjodin AR, Poisot T, Bergner LM, Chen B, et al. Optimising predictive models to prioritise viral discovery in zoonotic reservoirs. The Lancet Microbe. 2022;0. doi:10.1016/S2666-5247(21)00245-7/ATTACHMENT/926C4B33-05E2-4ADE-B493-301656D967F8/MMC1.PDF

4. Gibb R, Albery GF, Mollentze N, Eskew EA, Brierley L, Ryan SJ, et al. Mammal virus diversity estimates are unstable due to accelerating discovery effort. Biol Lett. 2022;18: 20210427. doi:10.1098/RSBL.2021.0427

5. Brierley L, Fowler A. Predicting the animal hosts of coronaviruses from compositional biases of spike protein and whole genome sequences through machine learning. PLoS Pathog. 2021;17. doi:10.1371/JOURNAL.PPAT.1009149

6. Greenhalgh T, Jimenez JL, Prather KA, Tufekci Z, Fisman D, Schooley R. Ten scientific reasons in support of airborne transmission of SARS-CoV-2. Lancet (London, England). 2021;397: 1603–1605. doi:10.1016/S0140-6736(21)00869-2

7. Pastorino B, Touret F, Gilles M, de Lamballerie X, Charrel RN. Prolonged Infectivity of SARS-CoV-2 in Fomites. Emerg Infect Dis. 2020;26. doi:10.3201/EID2609.201788

8. Moreira J, Peixoto TM, Siqueira AM, Lamas CC. Sexually acquired Zika virus: a systematic review. Clin Microbiol Infect. 2017;23: 296–305. doi:10.1016/J.CMI.2016.12.027

9. Thorson A, Formenty P, Lofthouse C, Broutet N. Systematic review of the literature on viral persistence and sexual transmission from recovered Ebola survivors: Evidence and recommendations. BMJ Open. 2016;6: e008859. doi:10.1136/BMJOPEN-2015-008859/-/DC1

10. Cortez MH, Weitz JS. Distinguishing between indirect and direct modes of transmission using epidemiological time series. Am Nat. 2013;181. doi:10.1086/668826/ASSET/IMAGES/LARGE/FG5.JPEG

11. Plowright RK, Parrish CR, McCallum H, Hudson PJ, Ko AI, Graham AL, et al. Pathways to zoonotic spillover. Nat Rev Microbiol 2017 158. 2017;15: 502–510. doi:10.1038/nrmicro.2017.45

12. Johnston S, Holgate S. Epidemiology of Viral Respiratory Tract Infections. Viral Other Infect Hum Respir Tract. 1996; 1–38. doi:10.1007/978-94-011-7930-0_1

13. Folly AJ, Sewgobind S, HernándezlJTriana LM, Mansfield KL, Lean FZX, Lawson B, et al. Evidence for overwintering and autochthonous transmission of Usutu virus to wild birds following its redetection in the United Kingdom. Transbound Emerg Dis. 2022 [cited 25 Oct 2022]. doi:10.1111/TBED.14738

14. Caminade C, Turner J, Metelmann S, Hesson JC, Blagrove MSC, Solomon T, et al. Global risk model for vector-borne transmission of Zika virus reveals the role of El Niño 2015. Proc Natl Acad Sci U S A. 2017;114: 119–124. doi:10.1073/PNAS.1614303114/SUPPL_FILE/PNAS.1614303114.SAPP.PDF

15. Bhatt S, Gething PW, Brady OJ, Messina JP, Farlow AW, Moyes CL, et al. The global distribution and burden of dengue. Nature. 2013;496: 504. doi:10.1038/NATURE12060

16. Alkhamis MA, Aguilar-Vega C, Fountain-Jones NM, Lin K, Perez AM, Sánchez-Vizcaíno JM. Global emergence and evolutionary dynamics of bluetongue virus. Sci Rep. 2020;10. doi:10.1038/S41598-020-78673-9

17. Yan Z, Wolters AMA, Navas castillo J, Bai Y. The Global Dimension of Tomato Yellow Leaf Curl Disease: Current Status and Breeding Perspectives. Microorg 2021, Vol 9, Page 740. 2021;9: 740. doi:10.3390/MICROORGANISMS9040740

18. Burrell CJ, Howard CR, Murphy FA. Epidemiology of Viral Infections. Fenner White’s Med Virol. 2017; 185. doi:10.1016/B978-0-12-375156-0.00013-8

19. Bragard C, Caciagli P, Lemaire O, Lopez-Moya JJ, Macfarlane S, Peters D, et al. Status and prospects of plant virus control through interference with vector transmission. Annu Rev Phytopathol. 2013;51: 177–201. doi:10.1146/ANNUREV-PHYTO-082712-102346/CITE/REFWORKS

20. Whitfield AE, Falk BW, Rotenberg D. Insect vector-mediated transmission of plant viruses. Virology. 2015;479–480: 278–289. doi:10.1016/J.VIROL.2015.03.026

21. Pagán I. Transmission through seeds: The unknown life of plant viruses. PLOS Pathog. 2022;18: e1010707. doi:10.1371/JOURNAL.PPAT.1010707

22. Dwyer GI, Gibbs MJ, Gibbs AJ, Jones RAC. Wheat streak mosaic virus in Australia: Relationship to Isolates from the Pacific Northwest of the USA and Its Dispersion Via Seed Transmission. 2007;91: 164–170. doi:10.1094/PDIS-91-2-0164

23. Wille M, Bröjer C, Lundkvist Å, Järhult JD. Alternate routes of influenza A virus infection in Mallard (Anas platyrhynchos). Vet Res. 2018;49: 1–9. doi:10.1186/S13567-018-0604-0/FIGURES/3

24. Krammer F, Smith GJD, Fouchier RAM, Peiris M, Kedzierska K, Doherty PC, et al. Influenza. Nat Rev Dis Prim 2018 41. 2018;4: 1–21. doi:10.1038/s41572-018-0002-y

25. Pierson TC, Diamond MS. The continued threat of emerging flaviviruses. Nat Microbiol. 2020;5: 796– 812. doi:10.1038/S41564-020-0714-0

26. Blitvich BJ, Firth AE. A Review of Flaviviruses that Have No Known Arthropod Vector. Viruses. 2017;9. doi:10.3390/V9060154

27. Bushman FD, McCormick K, Sherrill-Mix S. Virus structures constrain transmission modes. Nat Microbiol. 2019;4: 1778–1780. doi:10.1038/S41564-019-0523-5

28. Whitfield AE, Falk BW, Rotenberg D. Insect vector-mediated transmission of plant viruses. Virology. 2015;479–480<otherinfo>: 278–289. doi:10.1016/J.VIROL.2015.03.026</otherinfo>

29. Babayan SA, Orton RJ, Streicker DG. Predicting reservoir hosts and arthropod vectors from evolutionary signatures in RNA virus genomes. Science (80-). 2018;362: 577–580. doi:10.1126/science.aap9072

30. Tseng KK, Koehler H, Becker DJ, Gibb R, Carlson CJ, Del Pilar Fernandez M, et al. Viral genomic features predict orthopoxvirus reservoir hosts. bioRxiv. 2023; 2023.10.26.564211. doi:10.1101/2023.10.26.564211

31. Blagrove MS, Pilgrim J, Kotsiri A, Hui M, Baylis M, Wardeh M. Monkeypox virus shows potential to infect a diverse range of native animal species across Europe, indicating high risk of becoming endemic in the region. bioRxiv. 2022; 2022.08.13.503846. doi:10.1101/2022.08.13.503846

32. Heck M. Insect Transmission of Plant Pathogens: a Systems Biology Perspective. mSystems. 2018;3. doi:10.1128/MSYSTEMS.00168-17

33. Wu VY, Chen B, Christofferson R, Ebel G, Fagre AC, Gallichotte EN, et al. A minimum data standard for vector competence experiments. Sci Data 2022 91. 2022;9: 1–6. doi:10.1038/s41597-022-01741-4

34. Kraemer MUG, Sinka ME, Duda KA, Mylne AQN, Shearer FM, Barker CM, et al. The global distribution of the arbovirus vectors Aedes aegypti and Ae. albopictus. Elife. 2015;4. doi:10.7554/ELIFE.08347

35. Leung NHL. Transmissibility and transmission of respiratory viruses. Nat Rev Microbiol 2021 198. 2021;19: 528–545. doi:10.1038/s41579-021-00535-6

36. Jones RAC, Janssen D. Global Plant Virus Disease Pandemics and Epidemics. Plants 2021, Vol 10, Page 233. 2021;10: 233. doi:10.3390/PLANTS10020233

37. Sayers EW, Agarwala R, Bolton EE, Brister JR, Canese K, Clark K, et al. Database resources of the National Center for Biotechnology Information. Nucleic Acids Res. 2018 [cited 15 Nov 2018]. doi:10.1093/nar/gky1069

38. Gibb R, Albery GF, Becker DJ, Brierley L, Connor R, Dallas TA, et al. Data proliferation, reconciliation, and synthesis in viral ecology. bioRxiv. 2021; 2021.01.14.426572. doi:10.1101/2021.01.14.426572

39. Olival KJ, Hosseini PR, Zambrana-Torrelio C, Ross N, Bogich TL, Daszak P. Host and viral traits predict zoonotic spillover from mammals. Nature. 2017;546: 646–650. doi:10.1038/nature22975

40. Shaw LP, Wang AD, Dylus D, Meier M, Pogacnik G, Dessimoz C, et al. The phylogenetic range of bacterial and viral pathogens of vertebrates. Mol Ecol. 2020;29: 3361–3379. doi:10.1111/MEC.15463

41. Stephens PR, Pappalardo P, Huang S, Byers JE, Farrell MJ, Gehman A, et al. Global Mammal Parasite Database version 2.0. Ecology. 2017;98: 1476. doi:10.1002/ECY.1799/SUPPINFO

42. Wardeh M, Risley C, Mcintyre MK, Setzkorn C, Baylis M. Database of host-pathogen and related species interactions, and their global distribution. Sci Data. 2015;2. doi:10.1038/sdata.2015.49

43. Sastry KS, Mandal B, Hammond J, Scott SW, Briddon RW. Encyclopedia of Plant Viruses and Viroids. Encycl Plant Viruses Viroids. 2019. doi:10.1007/978-81-322-3912-3

44. Lefkowitz EJ, Dempsey DM, Hendrickson RC, Orton RJ, Siddell SG, Smith DB. Virus taxonomy: The database of the International Committee on Taxonomy of Viruses (ICTV). Nucleic Acids Res. 2018;46: D708–D717. doi:10.1093/nar/gkx932

45. Hulo C, De Castro E, Masson P, Bougueleret L, Bairoch A, Xenarios I, et al. ViralZone: A knowledge resource to understand virus diversity. Nucleic Acids Res. 2011;39: D576. doi:10.1093/nar/gkq901

46. Woolhouse MEJ, Brierley L. Epidemiological characteristics of human-infective RNA viruses. Sci Data. 2018;5. doi:10.1038/sdata.2018.17

47. Sharp PM, Li WH. The codon Adaptation Index--a measure of directional synonymous codon usage bias, and its potential applications. Nucleic Acids Res. 1987;15: 1281. doi:10.1093/NAR/15.3.1281

48. Kumar S, Suleski M, Craig JM, Kasprowicz AE, Sanderford M, Li M, et al. TimeTree 5: An Expanded Resource for Species Divergence Times. Mol Biol Evol. 2022;39. doi:10.1093/MOLBEV/MSAC174

49. Zhang ML, Li YK, Liu XY, Geng X. Binary relevance for multi-label learning: an overview. Front Comput Sci. 2018;12: 191–202. doi:10.1007/S11704-017-7031-7/METRICS

50. Ke G, Meng Q, Finley T, Wang T, Chen W, Ma W, et al. LightGBM: A Highly Efficient Gradient Boosting Decision Tree. [cited 13 Aug 2022]. Available: https://github.com/Microsoft/LightGBM.

51. Bunkhumpornpat C, Sinapiromsaran K, Lursinsap C. Safe-level-SMOTE: Safe-level-synthetic minority over-sampling technique for handling the class imbalanced problem. Lect Notes Comput Sci (including Subser Lect Notes Artif Intell Lect Notes Bioinformatics). 2009;5476 LNAI: 475–482. doi:10.1007/978-3-642-01307-2_43/COVER

52. Barua S, Islam MM, Yao X, Murase K. MWMOTE - Majority weighted minority oversampling technique for imbalanced data set learning. IEEE Trans Knowl Data Eng. 2014;26: 405–425. doi:10.1109/TKDE.2012.232

53. Rivera WA. Noise Reduction A Priori Synthetic Over-Sampling for class imbalanced data sets. Inf Sci (Ny). 2017;408: 146–161. doi:10.1016/J.INS.2017.04.046

54. Chawla N V, Bowyer KW, Hall LO, Kegelmeyer WP. SMOTE: Synthetic Minority Over-sampling Technique. J Artif Intell Res. 2002. Available: https://arxiv.org/pdf/1106.1813.pdf

55. Wilson DL. Asymptotic Properties of Nearest Neighbor Rules Using Edited Data. IEEE Trans Syst Man Cybern. 1972;2: 408–421. doi:10.1109/TSMC.1972.4309137

56. Tomek I. EXPERIMENT WITH THE EDITED NEAREST-NEIGHBOR RULE. IEEE Trans Syst Man Cybern. 1976;SMC-6: 448–452. doi:10.1109/TSMC.1976.4309523

57. Lundberg SM, Lee SI. A Unified Approach to Interpreting Model Predictions. Adv Neural Inf Process Syst. 2017; 2017–December: 4766–4775. doi:10.48550/arxiv.1705.07874

