## Supplementary Information File for "Features that matter: evolutionary signatures that predict viral transmission routes"

*Supplementary Materials*

Maya Wardeh*[1,2], Jack Pilgrim[2], Melody Hui[2], Aurelia Kotsiri[2], Matthew Baylis[2], Marcus SC Blagrove*[2]
*1) Department of Computer Science, University of Liverpool, Liverpool, UK*
*2) Institute of Infection, Veterinary and Ecological Sciences, University of Liverpool, Liverpool, UK*

### Supplementary Note 1 – Virus-host associations

**Supplementary Table 1 – Full sequenced viruses considered in this study.** Virus classification followed NCBI taxonomy [1].

| | Baltimore | Orders | Families | Genres | Species | Viruses | Description |
|---|---|---|---|---|---|---|---|
| **DNA** | **Group I** | 8 | 16 | 116 | 838 | 1,279 | Double-stranded DNA viruses (e.g. herpesviruses) |
| | **Group II** | 8 | 10 | 61 | 1,224 | 1,455 | Single-stranded DNA viruses (e.g. circoviruses) |
| **RNA** | **Group III** | 4 | 6 | 23 | 198 | 269 | Double-stranded RNA viruses (e.g. rotaviruses) |
| | **Group IV** | 13 | 42 | 190 | 1844 | 2,543 | Positive-sense single-stranded RNA viruses (e.g. flaviviruses) |
| | **Group V** | 6 | 24 | 103 | 789 | 966 | Negative-sense single-stranded RNA viruses (e.g. Influenza A virus) |
| **Retro-transcribing** | **Group VI** | 1 | 4 | 22 | 210 | 248 | RNA with DNA intermediate in life cycle (e.g. HIV-1) |
| | **Group VII** | 1 | 1 | 5 | 21 | 43 | DNA with RNA intermediate in life cycle (e.g. Hepatitis B) |

**Supplementary Table 2 – Summary of virus-host associations included in the study.** Virus and host classification followed NCBI taxonomy [1]; 28,661 associations between 5,750 host species (animals = 3,649, and plants = 2,101), and 6,803 viruses.

| | Baltimore | Associations | Viruses | Hosts | Vertebrates | Invertebrates | Plants |
|---|---|---|---|---|---|---|---|
| **DNA** | **Group I** | 2,864 (9.99%) | 1,279 (18.80%) | 997 (17.34%) | 879 (15.29%) | 118 (2.05%) | 0 (0%) |
| | **Group II** | 3,628 (12.66%) | 1,455 (21.39%) | 1,060 (18.43%) | 398 (6.92%) | 78 (1.36%) | 584 (10.16%) |
| **RNA** | **Group III** | 2,228 (7.77%) | 269 (3.95%) | 597 (10.38%) | 387 (6.73%) | 140 (2.43%) | 70 (1.22%) |
| | **Group IV** | 12,334 (43.02%) | 2,543 (37.38%) | 3,673 (63.88%) | 1,427 (24.82%) | 614 (10.68%) | 1632 (28.38%) |
| | **Group V** | 6541 (22.82%) | 966 (14.2%) | 2,285 (39.74%) | 1,473 (25.62%) | 433 (7.53%) | 379 (6.59%) |
| **Retro-transcribing** | **Group VI** | 932 (3.25%) | 248 (3.64%) | 448 (7.79%) | 236 (4.1%) | 44 (0.77%) | 168 (2.92%) |
| | **Group VII** | 134 (0.47%) | 43 (0.63%) | 101 (1.76%) | 100 (1.74%) | 1 (0.02%) | 0 (0%) |

### Supplementary Note 2 – Hierarchy of transmission modes and routes

**Supplementary Table 3 – Transmission routes included in this study.** The M column indicates if suites of models were trained for the given routes or due to insufficient data, the routes were incorporated in parent nodes along the transmission hierarchy.

| Mode | M | Routes | Definition |
|---|---|---|---|
| **Vertical** | 1 | Vertical pre (mammalian); vertical pre (egg) | Transmission to foetus within uterus; to embryo within vertebrate egg. |
| | | Transovarial | Transmission from an infected female arthropod to its offspring through the eggs during reproduction. |
| | | Germline integration | Transmission via integrated viral genomes which are present in the germline. |
| | | Pollen; Seed | Transmission of plant viruses via pollen or seed. |
| | | Vegetative propagation | Transmission via asexual plant reproduction, in which an off-spring plant is grown from a fragment/cutting of the parent plant. |
| | | During (mammalian birth) | Transmission from parent to offspring during mammalian birth. |
| | | Trans-egg (vertebrates); trans-egg (invertebrates) | Transmission during egg-hatching, where the virus has contaminated the surface of the eggs. |
| | 0 | Perinatal/peri-hatching | Transmission from (maternal) parent to offspring within the perinatal/peri-hatch period |
| | | Oviposition | Transmission from arthropod parent to offspring during oviposition. |
| | 1 | Transstadial | Transmission (in arthropods) from one developmental stage (e.g., larvae or nymphs) to the subsequent life stage (e.g., adults). |
| | | Vertical (post) | Transmission from (maternal) parent to offspring via breast-milk, colostrum etc. |
| **Sexual** | 1 | Genital-genital contact; Semen/Sperm | Transmission via sexual contact. |
| | 0 | Oral-genital contact | |
| **Bodily-fluids** | 1 | Blood; Faeces; Saliva; Urine | Transmission via direct contact with bodily fluids |
| | | contact with bodily-fluids | |
| **Feeding contact** | | Arthropod feeding | Transmission via feeding contact such as arthropod feeding (from plant or vertebrate to the arthropod), or predation and cannibalism |
| | | Predation/cannibalism | |
| **Direct contact** | | Oral-skin/bloodstream contact | Transmission through broken skin via bites or scratching. |
| | | Oral-oral contact | Transmission via oral contact (e.g. kissing, grooming) |
| | | Respiratory | Droplet/airborne transmission from one individual to another (e.g. via coughing, sneezing, breathing). |
| | | Skin-skin/eye contact | Transmission via direct physical contact with skin or skin to eye (of animals) |
| | | Plant contact | Transmission via direct contact between plants. |
| **Ingestion** | 1 | Food/water | Transmission via ingestion of food/water contaminated with virus particles or containing faecal matter with virus particles. |
| | | Faecal-oral | |
| | 0 | Pollen (food) | |
| **Indirect contact** | 0 | Cuscuta | Minor indirect contact routes. |
| | | Indirect contact (insects) | |
| **Environmental** | 1 | Air (dry); Air (wet) | Environmental transmission via inhalation of wet (e.g. rodent urine) or dry (e.g., dust) virus particles from the environment. |
| | | Soil | Transmission of virus in the soil (excluding soil-dwelling organisms such as fungi/nematodes) |
| | | Water-borne | Transmission via contact with water (excluding ingestion). |
| | | Passive diffusion | Transmission through gills/spread through mucus by passive diffusion (from water). |
| | | Cohabitation | Transmission of virus horizontally (in the water) between cohabitating aquatic animals. |
| | | Fomite | Transmission via contact with inanimate objects (e.g., gloves, tools) contaminated with the virus. |
| | | Indirect contact with bodily-fluids | Transmission via inanimate objects contaminated with bodily-fluids (such as needles, surgical tools). |
| | | sap inoculation | Controlled mechanical transmission process, which typically involves extracting plant sap from an infected plant and then introducing this sap into a healthy plant. |

| | | | |
|---|---|---|---|
| | 0 | Co-feeding | Transmission occurs when infected and uninfected arthropods feed in proximity to each other on the same reservoir host. |
| **Arachnid-borne** | 1 | Mite-borne; tick-borne | Transmission by arthropod vectors (vector-borne, including mechanical transmission). These routes indicate the mechanism of viral transmission to the vertebrate or plant host (e.g. Zika virus is mosquito-borne to humans; Tomato yellow leaf curl virus is whitefly-borne to tomatoes).<br>The routes/modes of transmission to the arthropod vector are categorised under the mechanism by which the vector obtains the virus (e.g. arthropod feeding, transovarial transmission, sexual transmission, etc). the replication of virus (if any) in the arthropod is captured via various vectoring mechanisms such as: circulative, non-circulative, non-persistent, semi-persistent, non-propagative, and propagative transmission (below). |
| **Insect-borne** | 1 | beetle-borne; thrip-borne; leafhopper-borne; planthopper-borne; midge-borne; mosquito-borne; sandfly-borne; aphid-borne; mealybug-borne; whitefly-borne | |
| | 0 | grasshopper-borne; leafminer-borne; mayfly-borne; louse-borne; weevil-borne; fly-borne; treehopper-borne; horsefly-borne; housefly-borne; tabanidae-borne; blackfly-borne; bug-borne; cimicoidea-borne; coccoidea-borne | |
| **Other-vectors** | 1 | Fungi/plasmodiophorids; nematode-borne | Transmission by non-arthropod vectors (excluding vertebrate hosts/reservoirs). |
| | 0 | leech-borne | |
| **Non-circulative** | 1 | Non-persistent | Non-persistent viruses are transmitted mechanically by the vector. Transmission typically occurs within a short time frame, often within seconds to minutes, as the virus is carried on the surface of the vector's mouthparts or stylets. |
| | 1 | Semi-persistent | Semi-persistent viruses are retained within the vector for a longer duration compared to non-persistent viruses, typically ranging from hours to days. While the virus is retained, it does not undergo replication or systemic infection within the vector; the extended retention period allows for a greater potential for transmission to new hosts. |
| **Circulative (persistent)** | 1 | Non-propagative | Non-propagative viruses do not replicate within the vector. These viruses are acquired by the vector during feeding on an infected host but do not undergo replication or amplification within the vector's cells. Transmission occurs solely through the transfer of virions from the infected host to a susceptible host during subsequent feeding by the vector |
| | 1 | Propagative | Propagative viruses are capable of replicating and multiplying within the vector. After acquisition by the vector, these viruses infect and replicate within specific tissues or cells of the vector (e.g. midgut or salivary glands) |

**Supplementary Table 4 – Summary of virus-host associations for which at least one transmission route was identified.** Virus and host classification followed NCBI taxonomy[1].

| | **Baltimore** | **Associations** | **Viruses** | **Hosts** | **Vertebrates** | **Invertebrates** | **Plants** |
|---|---|---|---|---|---|---|---|
| **DNA** | **Group I** | 2,229 (8.93%) | 890 (20.02%) | 825 (15.5%) | 714 (86.55%) | 111 (13.45%) | 0 (0%) |
| | **Group II** | 2,942 (11.79%) | 899 (20.22%) | 899 (16.9%) | 327 (36.37%) | 55 (6.12%) | 517 (57.51%) |
| **RNA** | **Group III** | 2,106 (8.44%) | 165 (3.71%) | 531 (9.99%) | 376 (70.81%) | 115 (21.66%) | 40 (7.53%) |
| | **Group IV** | 11,059 (44.32%) | 1,771 (39.83%) | 3,427 (64.5%) | 1,308 (38.17%) | 548 (15.99%) | 1,571 (45.84%) |
| | **Group V** | 5,783 (23.175%) | 529 (11.90%) | 2,091 (39.3%) | 1,375 (65.76%) | 307 (14.68%) | 409 (19.56%) |
| **Retro-transcribing** | **Group VI** | 756 (3.03%) | 161 3.62%) | 386 (7.26%) | 210 (54.4%) | 41 (10.62%) | 135 (34.97%) |
| | **Group VII** | 78 (0.31%) | 31 (0.697%) | 60 (1.13%) | 60 (100%) | 0 (0%) | 0 (0%) |

### Supplementary Note 3 – Viral Features

**Supplementary Table 5 – Viral features groups.** Features were first calculated from sequences and then averaged for each included virus (strain or species). F indicates the number of features computed per group.

| Category | Group | F | Details |
|---|---|---|---|
| **Genome** | **Length** | 1 | Affects available space for protein encoding and gene expression regulation, number of mutations/genome/replication, and physical size of virion [2]. |
| | **GC content** | 1 | GC content affects thermal (and other stressor) stability of genome (and resultant RNA) and genomic regions [3]. |
| | **Structure** | 7 | 1. RNA (binary, 0 = DNA); RNA viruses generally have a higher mutation rate [4], and are generally more fragile (cannot survive as long outside of the cell).<br>2. Retro-transcribing (binary, 1=yes, NCBI taxonomy[1]); Retroviruses are often very conserved[5], and have to enter the nucleus[6] and insert into the genome, these additional steps may require specificity and limit range, hence adaption to direct transmission is advantageous.<br>3. (-/+) Sense (binary, NCBI taxonomy [1]); Sense affects replication cycle and range of host cellular machinery that needs to be recruited.<br>4. Linear (binary, 0 = circular, ViralZone [7] and ICTV [8]); This attribute affects replication and translation. Rolling circle replication and translation are common with circular genomes, negating the need to re-enlist host enzymes [9]. Linear genomes are often tightly bound with nucleocapsid proteins which affect environmental stability.<br>5. Single stranded(binary, 0= double, ViralZone [7] and ICTV [8]); single stranded viral genomes are often small, mutate and recombine readily, and on evolutionary timescales show more frequent evidence of horizontal gene transfer [10]. Hence, ss genomes readily adapt and evolve to different niches.<br>6. Segmented (binary, 0 = monopartite, ViralZone [7] and ICTV [8]); we noted if the virus was monopartite (has a single nucleic acid molecule protected in a shell made of proteins) or segmented (divided into two or more nucleic acid segment) (binary factor, no=monopartite, obtained from ViralZone [7] and ICTV [8]). Segmented viruses can undergo reassortment if two strains of the same virus infect a cell (e.g. influenza hemagglutinin & neuraminidase recombination [11]). This in turn can lead to host range changes of segments of the genome. In practice, there exists a third class of viral architecture - multipartite viruses. These viruses have their genome divided into two or more nucleic acid segment (similarly to segment viruses), but these segments are each packaged into separate virus particles. We ignored multipartite viruses in this study due to them being very rare and poorly understood [12]. |
| **Morphology (capsid)** | | 16 | We indicated if the virus is enveloped or not (binary, ViralZone [7]). Envelopes are usually derived from the host cell membrane; this can help them avoid host immune system. The envelopes can be very sensitive to the external environment, and enveloped viruses often require to be directly transferred between hosts; finally, because the envelope is made from the current host's cell membrane, it will change upon infection of a new host, making the virus rapidly adaptable [13].<br>We included 12 binary variables indicating the morphology of the virus; and three binary variables indicating further characteristics of the virus structure. Viral structure has been found to correlate with routes of transmission [14]. |
| **Replication** | | 2 | We collated information on the replication site of the virus (ViralZone [7] and ICTV [8]). We expressed these data as a binary factor indicating if the virus replicates in the cytoplasm and/or in the nucleus. Replication site is linked to RNA/DNA genome – if a virus has a DNA |

| | | | |
|---|---|---|---|
| | | | stage it usually replicates in the nucleus. This creates extra barriers to overcome for entry to the nucleus and may restrict host range. |
| **Biases** | **Nucleotide bias** | 4 | Nucleotide bias can result in genome-wide biases, amino acid composition, and function [15]. |
| | **Dinucleotide bias** | 128 | Dinucleotide bias is a direct precursor to codon pair bias[16]. Dinucleotide composition is influenced by more distant virus family relationships than closer species-level differentiation [17]. |
| | **Relative Synonymous Codon Usage (RSCU).** | 204 | Codon biases in viral genome affect translation in host cell. Similarities/differences affect tRNA recruitment, and production speed [18]. Can also affect virulence in host [19]. In many mammals, different tissues have different codon biases [20], therefore, tissue tropism and hence transmission routes may reflect different biases. |
| | **Amino acids bias** | 57 | Amino acid categories affect protein structure (e.g. proline), pH, solubility, etc. and hence proteins' properties in cellular or extracellular environments [21]. |
| **ORFs** | **Composition** | 9 | Proportion of genome utilised by ORFs, the sizes of the ORFs, and the proportion of overlapping ORFs (between different frames) describes genome utilisation for protein production, genome density, and constraints on AA sequence (from overlap). |
| | **Coverage** | 6 | |
| | **ORF Overlap** | 21 | |

**Pre-processing.** Sequences with ambiguous bases that could be resolved in fewer than or equal to 1,024 permutations were expanded to resolve ambiguity using Disambiguate function in the R package Decipher [22]. This process resulted in a total = 139,474 sequences (127,602 with known transmission route to at least one host species, 11,872 sequences without known transmission route).

**ORF generation.** Non-overlapping set containing longest ORFs was predicted for included sequence (n=139,474), and its reverse complement, using predORF function in the R package systemPipeR (parameters were set to n='all' and longest_disjoint=TRUE to subset to non-overlapping ORF set containing longest ORF). Predicted ORFs of length <36 were dropped from the analyses. Remainder ORFs were grouped into three overlapping categories: length≥36; length≥300; length≥450. RSCU and ORFs-derived features (Supplementary Table 5, below), were computed for each category.

**Genome wide features.** GC content, nucleotide biases (proportion of each nucleotide in the sequence), dinucleotide biases, and genome length were calculated for each of the inambiguous/disambiguated sequences.

In additions, we computed dinucleotide biases as follows [23]:

$$D_{xy} = \frac{\frac{f_{xy}}{D}}{\left(\frac{f_x}{N} \times \frac{f_y}{N}\right)}$$

Where $f_{xy}$ denotes the frequency of dinucleotide $xy$, $f_x$ and $f_y$ denote frequency of individual nucleotides $x$ and $y$, and $D$ and $N$ denote the total number of dinucleotides and nucleotides in the given sequence, respectively. We also quantified dinucleotide biases at each position within codon reading frames (i.e. positions 1-2, 2-3, or 3-1, p1, p2, and p3 respectively) for each sequence, and for the reverse complement of the sequence (pr, p1r, p2r, and p3r, respectively), thus resulting in total of 128 features depicting dinucleotide biases of each included sequence.

**ORF Composition.** We computed the following features to express ORF composition for each sequence:

| | | |
|---|---|---|
| **Sense bias** | $\frac{n_{sense_c}}{length_{sequence}}$ | $n_{sense_c}$ is the number of ORFs predicted from the sequence, of length≥ $c$ (c=36, 300, 450). |
| **Asense bias** | $\frac{n_{asense_c}}{length_{sequence}}$ | $n_{asense_c}$ is the number of ORFs predicted from the reverse complement of the sequence, of length≥ $c$ (c=36, 300, 450). |
| **Sense-Probability** | $\frac{n_{sense_c}}{n_{sense_c} + n_{asense_c}}$ | |
| **Asense-proportion** | $\frac{n_{asense_c}}{n_{sense_c}}$ | |

**ORF coverage.** For each sequence, we computed a coverage vector (length = sequence length, initialised with 0s). For each (non-overlapping) predicted $ORF_i$, coverage vector is updated such that:

$$Coverage(start_{ORF_i}, end_{ORF_i}) = inframe2end_{ORF_i} \quad \textbf{(S.1)}$$

Where $inframe2end_{ORF_i}$ is frame of identified ORF/CDS relative to 3' end of query sequence. For ORFs predicted from each sequence (sense), $inframe2end_{ORF_i} \in \{1,2,3\}$, where value 1 stands for in-frame with downstream ORF, whereas 2 or 3 indicates a shift of one or two bases, respectively. For ORFs predicted from reverse complements (asense), $inframe2end_{ORF_i} \in \{4,5,6\}$, where value 4 stands for in-frame with downstream ORF, whereas 5 or 6 indicates a shift of one or two bases, respectively.

1. Sense coverage = proportion of elements in coverage with value ∈ {1,2,3},
2. Asense coverage = proportion of elements in coverage with value ∈ {4,5,6},

We computed the above features for each of our ORF length cut-offs (≥36, ≥300, ≥450), thus resulting in total of six ORF coverage features.

**ORF overlap.** We expressed ORF overlap, at each ORF length cut-off, in seven features ($T_{0_c}$ to $T_{6_c}$, where $c$ is our cut-off (c=36, 300, 450)) whereby, for each possible frame value $k$ (0 = no predicted ORFs, 1 = in-frame with downstream ORF, 2 or 3 a shift of one or two bases, 5 = in-frame with downstream ORF (reverse complement), 5 or 6, a shift of one or two bases), $T_k$ = proportion of elements in coverage = $k$.

**Relative Synonymous Codon Usage (RSCU).** Frequency of each codon (n = 64) was calculated for each predicted ORF at each cut-off, and then summed across all ORFs obtained from the same sequence, at each cut-off. RSCU was then computed for each codon (including stop codons)[24], for each cut-off ($c$ ≥36, ≥300, ≥450) as follows:

Let $n_i$ be the number of codons synonymous for amino acid $AA_i$ and $C_{ij_c}$ the frequency of the $j^{th}$ codon encoding for $AA_i$, for cut-off $c$, across all ORFs obtained from the sequence whose length≥ $c$, then:

$$RSCU(C_{ij}, c) = \frac{C_{ij_c}}{\frac{1}{n_i} \times \sum_j^{n_i} C_{ij_c}} \quad \textbf{(S.2)}$$

**Amino Acids biases.** Frequency for each amino acid was computed for each predicted ORF, for each cut-off ($c$ ≥36, ≥300, ≥450), and then summed across all ORFs obtained from the same sequence, at each cut-off.

Amino acids were categorised into 19 overlapping binary categories expressing:

1. Hydropathy: neutral, hydrophilic, hydrophobic.

2. Volume: very large, large, medium, small, very small.
3. Charge: negative, positive.
4. Polar.
5. Hydrogen donor or acceptor.
6. Chemical: acidic, aliphatic, amide, aromatic, basic, hydroxyl, and sulphur.

Bias for each amino acid category was then computed as frequency of all amino acids encoding for each category, in the sequence, divided by the total number of amino acids in the sequence. This resulted in total of 57 features per sequence.

**Segmented viruses.** Pre-processing and ORF generation were applied to individual segments. Features were calculated for the full genome - all segments belonging to same strain of virus, identified from via sequence meta-data, and all ORFs derived from these segments.

**Post processing**. Computed feature values were averaged per all sequences for each included virus (n=7,853) to generate final features.

**Supplementary Note 4 – Hosts similarity**

We obtained a time tree of 4,342 animal and plant species from the Time Tree of Life [25] (timetree.org). This enabled us to directly calculate the diversion time between 9,428,653 pairs of species. We adopted the following two-step routine to establish a diversion time between species pairs for which we could not obtain direct values for from the Time Tree of Life [25] (1,571 species, total = 8,056,088 pairs):

1. **Distances between taxonomically distant species**: We precomputed distances between species from different kingdoms/clades/phylum/…/class as the distance between their respective kingdoms/clades/phylum/…/classes. For instance, distance between any animal species and any plant species was computed as: 2991.528; the distance between any vertebrate and any arthropod species was computed as: 1593.111; and the distance between any mammalian and any avian species was computed as: 623.8078.
2. **Distances between taxonomically close species**: For species within the same class (e.g. mammals), we calculated distances as follows:
   **a.** Distance between families: distance between any two species from different families was fixed as the distance between their respective families.
   **b.** Distance between genres: distance between any two species from different genres was fixed as the distance between their respective genres.
   **c.** Distances between species within the same genus: distance between any two species within the same genus was fixed as the average distance between species (with known diversion times) within the genus.

Following the above process, distances were computed for 99.98% of all included species pairs. Given a focal route/mode $r$, and a focal association $v_i h_m$, we utilised the resulting diversion time distances to compute hosts similarity metric as the average (diversion) distance between $h_m$ and $\forall h_n \in v_j h_n$, where $v_j$ is transmitted to $h_n$ via $r$, and $h_m \neq h_n$.

### Supplementary Note 5 – Virus-host integrated neighbourhoods

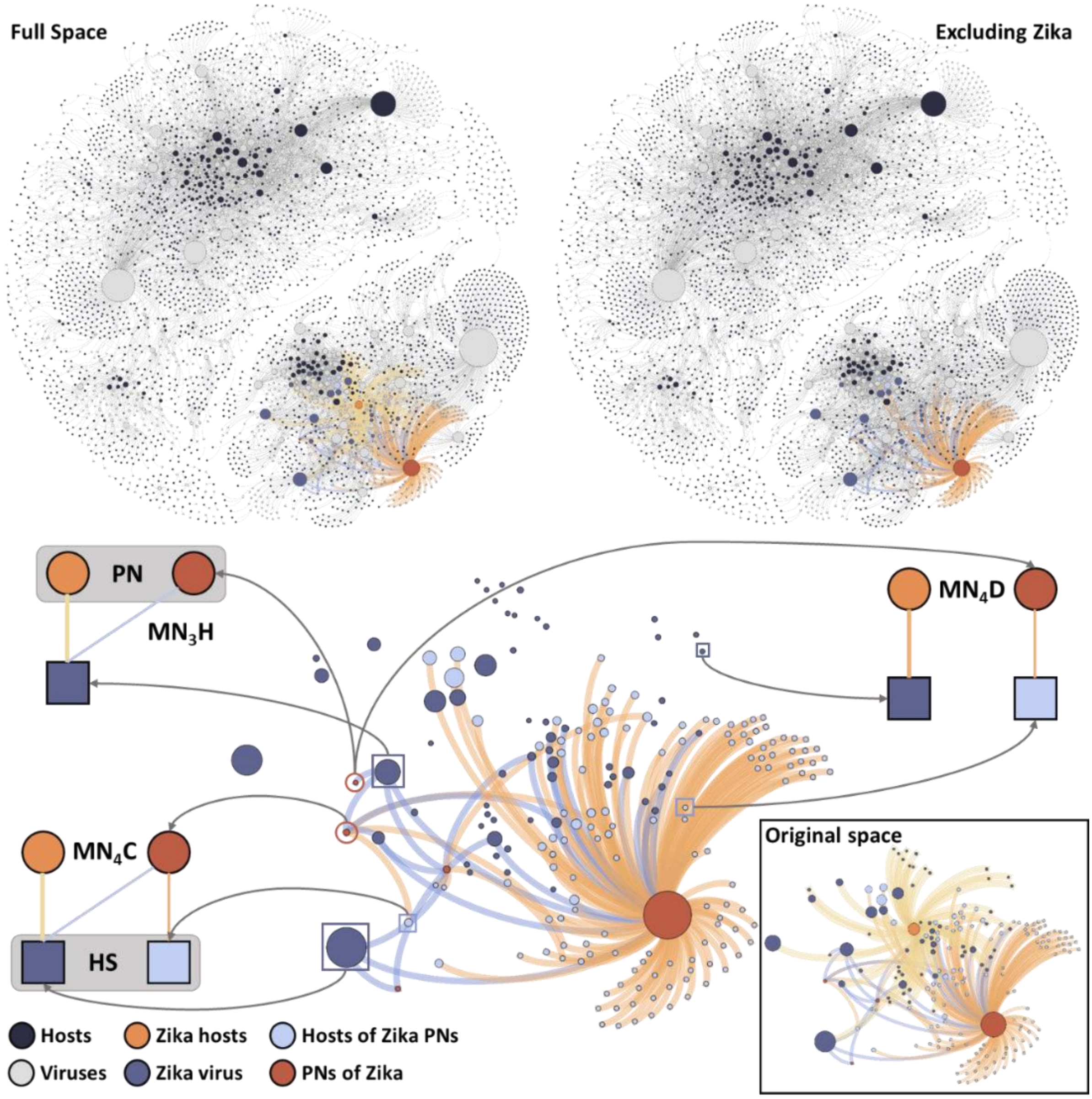


**Supplementary Figure 1 – Graphical abstraction of virus-host integrated neighbourhoods**. Here, the focal route/mode is insect-borne transmission, and the focal virus is Zika virus. Firstly, the focal virus is removed from the set of virus-host associations where by the virus is known to be transmitted to the host by the focal route/mode. Secondly, the Phylogenetic Neighbourhood (PN) is identified for the focal virus (by blasting against local database). Finally, the virus-host integrated neighbourhoods (VHINs) are constructed for each association between the focal virus, and each of its known hosts. Three similarity-based features are computer from the resulting VHINs.

As the same species or strain of virus may employ a varied range of transmission routes to infect different hosts, coupled with the fact that closely related viruses may utilise a diverse set of transmission routes in different hosts, we expanded the concept of phylogenetic neighbourhoods [23], so that for a virus-host association, similarities between both the focal virus and viruses within its phylogenetic neighbourhood, as well as those between the focal host and hosts of those neighbouring viruses, are incorporated in to a unified framework, as follows (Supplementary Figure 1):

**A. Local database construction**. In each iteration of our pipelines (n = 50, Supplementary Note 7), we randomly sampled the set of complete and unambiguous/disambiguated sequences (Supplementary Note 1), as follows:

1. Monopartite viruses: a single representative sequence was randomly sampled for each unique taxid (NCBI organism identifier).
2. Segmented viruses: a single representative sequence was randomly sampled for each unique segment (e.g., S, M or L), and taxid combination.

We then built customised local databases of sampled virus sequences, known to be transmitted via focal route/mode $r$ to at least one host species, using makeblastdb in the R package rBLAST (BLAST version 2.7.1+).

**B. Virus Phylogenetic Neighbourhood (PN).** Per each iteration, given a focal virus ($v_i$), a set of representative sample sequences of $v_i$ ($seq_{v_i}$), and a local database of focal route/mode $r$ ($db_r$), we used blastn to find the top hits of $seq_{v_i}$ (excluding those belonging to $v_i$) in $db_r$, based on e-values. We used the function predict in the R package rBLAST to apply blastn as follows:

predict($db_r$, $seq_{v_i}$, BLAST_args="-num_threads 8 -max_target_seqs=6 -max_hsps 1 -reward 2 -task blastn -evalue 10 -word_size 8 -gapopen 2 -gapextend 2")

Hits were aggregated at the level of virus species or strain (termed group in Supplementary Dataset 1), and the resulting unique top five hits were considered the PN of the focal virus $v_i$, for a give route $r$ (${PN_{v_i}}^r$).

**C. Virus-host integrated neighbourhoods (VHINs)**. Given a focal route $r$, and a focal association: $v_i h_m$, we constructed the neighbourhood network of $v_i h_m$, via $r$ ($\mathrm{NN}^r_{v_i h_m}$), as follows:

1. Nodes comprised viruses in $PN_{v_i}$, hosts they are known to infect via $r$, as well as known hosts of the focal virus.
2. Nodes were linked via an edge if the virus is known to infect the host via $r$.

We then computed a set of three features for each focal association $v_i h_m$:

| | |
|---|---|
| | $\boldsymbol{{PN_{v_i}}^r}$ is the phylogenetic neighbourhood of virus $v_i$ , a set of 5 (maximum) viruses representing the top five hits resulting from applying *blastn* of representative sequence against $r$ local database. |
| | $\boldsymbol{S_{h_m h_n}} = 1 - \frac{DT_{mn}}{DT_{max}}$ is the normalised similarity between two host species $h_n$ and $h_m$. $DT_{mn}$ is diversion time between $h_n$ and $h_m$, and $DT_{max}$ is the maximum diversion time between all included species. |
| | ${V_{h_m}}^r$ is the set of viruses known to infect the focal host $h_m$ (excluding focal virus $v_i$) via $r$. |
| | ${H_{v_j}}^r$ is the set of host species known to be susceptible to virus $v_j \in {PN_{vi}}^r$ (excluding focal host $h_m$) via $r$. |
| | $P_{v_i v_j}$ is the pairwise genetic identity between focal virus $v_i$ and hit $v_j \in {PN_{v_i}}^r$ |

| Feature | Formula | Relevance |
|---|---|---|
| $\mathbf{MN_3H}_{v_i h_m}$ | $\frac{\sum_{v_j \in {PN_{vi}}^r \wedge v_j \in {V_{h_m}}^r} P_{v_i v_j}}{\lvert v_j {h_m}^r \rvert_{v_j \in {PN_{vi}}^r \wedge v_j \in {V_{h_m}}^r}}$ | Indicates whether the focal host $h_m$ is susceptible, via route/mode $r$, to viruses that exhibit high sequence similarity (closely related) to the focal virus $v_i$. Higher values might indicate higher likelihood $v_i$ is transmitted to $h_m$ via $r$. |
| $\mathbf{MN_4C}_{v_i h_m}$ | $\frac{\sum_{v_j \in {PN_{vi}}^r \wedge v_j \in {V_{h_m}}^r} \sum_{h_n \in {H_{v_j}}^r} P_{v_i v_j} \times S_{h_m h_n}}{\lvert v_j {h_m}^r \rvert_{v_j \in {PN_{vi}}^r \wedge v_j \in {V_{h_m}}^r} \times \lvert v_j {h_n}^r \rvert_{v_j \in {PN_{vi}}^r \wedge v_j \in {V_{h_m}}^r \wedge h_n \in {H_{v_j}}^r}}$ | Measures the average similarity between the focal association ($v_i h_m$) and each association ($v_j h_n$ - $h_m \neq h_n$), where $v_j$ is known to be transmitted via $r$ to both hosts: $h_n$ and $h_m$. |
| $\mathbf{MN_4D}_{v_i h_m}$ | $\frac{\sum_{v_j \in {PN_{vi}}^r \wedge v_j \in {V_{h_m}}^r} \sum_{h_n \in {H_{v_j}}^r} P_{v_i v_j} \times S_{h_m h_n}}{\lvert v_j {h_m}^r \rvert_{v_j \in {PN_{vi}}^r \wedge v_j \notin {V_{h_m}}^r} \times \lvert v_j {h_n}^r \rvert_{h_n \in {H_{v_j}}^r}}$ | Measures the average similarity between the focal association ($v_i h_m$) and each association ($v_j h_n$ - $h_m \neq h_n$), where $v_j$ is known to be transmitted via $r$ to $h_n$ but not to $h_m$. |

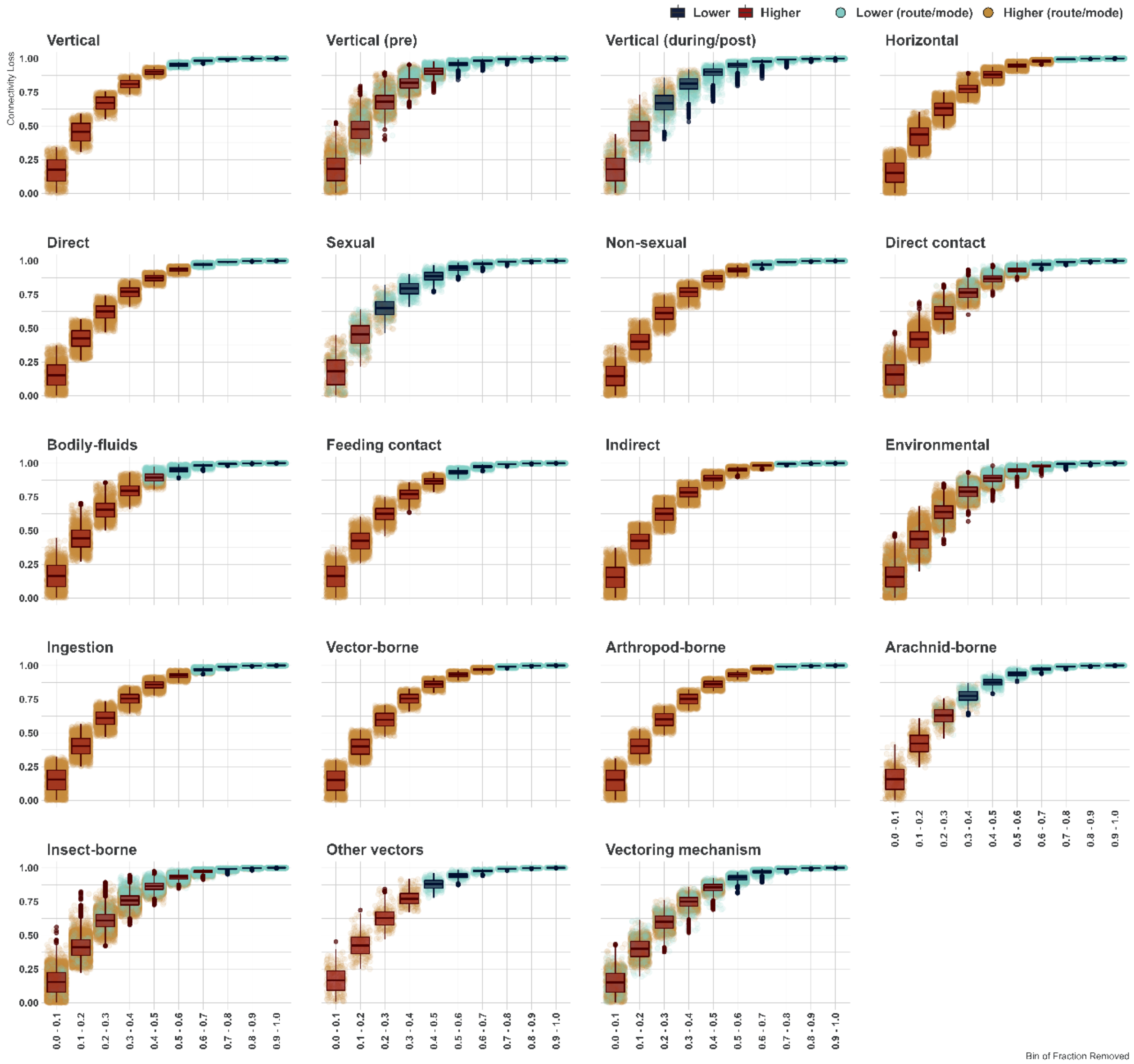


**Supplementary Figure 2 – Stability of virus-host association networks by category of transmission.** We explored the stability virus-host association networks, in which nodes represent viruses and their hosts, and edges (links) indicate that the virus is transmitted to the host via a given route/mode. Stability indicate the ability of a network to withstand perturbations without significant degradation in connectivity or efficiency. We quantified this stability by measuring connectivity loss per fraction of nodes removed at random from each network. We also compared the loss of connectivity exhibited by our networks against random networks generated using the Erdős-Rényi (ER) Random Graph Model, and exhibiting the same number of nodes and edges are our original networks. The stability analyses were conducted using the R Package NetSwan, and results were grouped into 19 categories (mode) of transmission. Fraction of nodes removed were binned into 10 equal bins, and results are summarised as boxplots representing the interquartile range (IQR), of the data distribution per bin. Horizontal lines within the box represent the median of the data distribution. Whiskers extend from the edges of the box to the minimum and maximum values within a distance of 1.5 times the IQR from the nearest quartile, individual data points that fall outside the range covered by the whiskers are plotted as outliers. Points represent the outcome from individual networks (each representing one route/mode). Boxplots are coloured to indicate if the connectivity loss is higher (red) or lower (blue), on average, than the loss of connectivity measured in the corresponding random networks per each category. Points are coloured to indicate if the connectivity loss is higher (orange) or lower (turquoise), on average, than the loss of connectivity measured in the corresponding random networks per each route/mode.

### Supplementary Note 6 – Class balancing

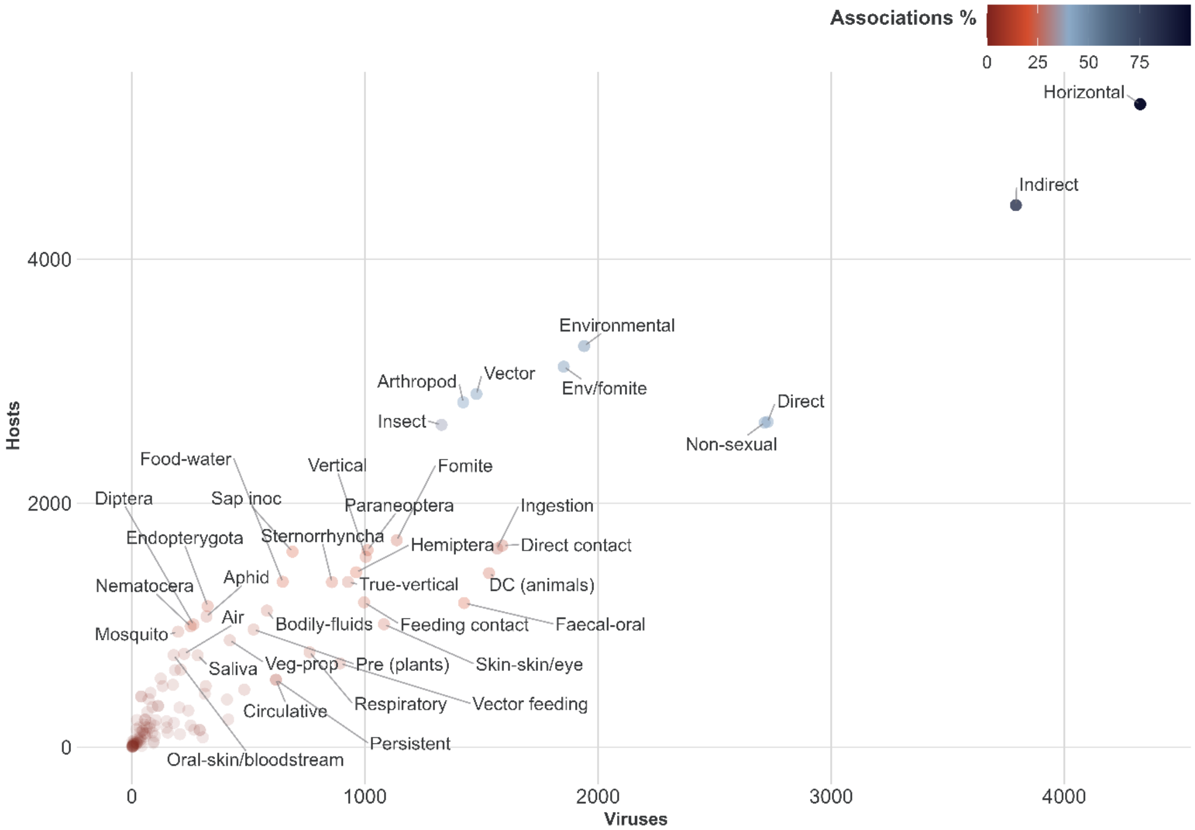


**Supplementary Figure 3 – Class Bias in transmission routes/modes included in this study.** Points represent individual routes/modes modelled in this study (n=98, Figure 1). Points are coloured by % of virus-host associations observed per route/mode. Transparency (alpha) is in relation to the percentage of virus-host associations observed per route/mode (n= 24,953). X-axis represent number of observed unique viruses per route/mode. Y-axis represent number of observed unique hosts per route/mode.

**Supplementary Table 6 – Class balancing techniques used in this study.** % refers to percent of minority class instances in the resulting training set. All balancing was applied to training sets prior to training and was performed using R packages: bimba and imbalance (ENN and TL only), and lightGBM (SPW only). Due to the small proportion of observed associations per majority of routes/modes (median = 763 associations, 3.06% of total), we elected to correct for class imbalance using a range of over-sampling and hybrid methods, rather than strict under-sampling.

| Technique | Sampling | Method | % |
|---|---|---|---|
| **SMOTE** | **Over-sampling** | SMOTE (Synthetic Minority Over-Sampling Technique [26]) synthesises new minority class instances from existing cases using a k-nearest neighbour algorithm. SMOTE then over-samples from the minority instances (original and synthesised) and under-samples from the majority class to create a balanced training set. | 50% |
| **SMOTE (25%)** | | | 25% |
| **BL-SMOTE** | | Borderline-SMOTE [27] classified each instance of the minority class based on the number of its minority class neighbours. Instances are categorised as noise (when the number of minority neighbours is 0) or safe (when the ratio of minority neighbours is greater than 50%). These two sets are excluded from the synthetic data generation process. Instead, new instances are created along the decision boundary between minority and majority classes, unlike SMOTE, which generates minority instances randomly. Borderline-SMOTE offers two options; we have adopted the second option, wherein only the minority class is oversampled, as opposed to the first option, which oversamples from both classes. | 50% |
| **BL-SMOTE (25%)** | | | 25% |
| **SL-SMOTE** | | Safe-Level-SMOTE [28] introduces the concept of a 'safe level,' which represents the number of majority instances among the k nearest neighbours of a minority class instance. Instances with a safe level close | 50% |
| **SL-SMOTE (25%)** | | | 25% |

| | | | |
|---|---|---|---|
| | | to 0 are classified as noise, while those close to k are considered safe. Unlike Borderline-SMOTE, Safe-Level-SMOTE synthesises minority instances in 'safe positions' by considering the safe level ratio of these instances. This approach avoids using outlier minority instances during the generation of synthetic instances. | |
| **ADASYN** | | Adaptive Synthetic Sampling (ADASYN) [29] dynamically synthesises minority instances in inverse proportion to the density of minority instances in their neighbourhood. Essentially, ADASYN generates more instances in regions of the feature space where the density of minority instances is low, and fewer (or none) where the density is high. Unlike SMOTE, which generates the same number of synthetic instances for each original minority instance, ADASYN automatically adjusts the number of new instances generated for each minority instance to compensate for skewed distributions. This adaptive approach helps address class imbalance more effectively. | 50% |
| **ADASYN (25%)** | | | 25% |
| **MWMOTE** | | Majority Weighted Minority Over-Sampling Technique (MWMOTE) [30] begins by identifying informative minority class instances that are difficult to learn. These instances are weighted based on their Euclidean distance from the nearest majority class neighbours (with varying k values such as k=1, k=2, k=3, etc.). Subsequently, MWMOTE synthesises minority instances from these selected weighted informative minority class instances using a hierarchical clustering approach. MWMOTE is less susceptible to the effects of noisy instances than SMOTE. | 50% |
| **MWMOTE (25%)** | | | 25% |
| **RWO** | | Random Walk Over-Sampling (RWO) [31] leverages the Central Limit Theorem to balance the minority class while maintaining the data distribution. It achieves this by perturbing existing minority instances to synthesise new ones. Moreover, RWO extends the minority class boundary after generating synthetic samples. | 50% |
| **RWO (25%)** | | | 25% |
| **SMOTE (NRAS)** | **Noise reduction & over-sampling hybrid** | Noise Reduction A Priori Synthetic Over-Sampling (NRAS) [32] is employed to pre-process training data by removing minority instances with a proportion of minority examples among their k nearest neighbours below a specified threshold (default = 50%). We empoloyed NRAS to pre-process training data, as implemented in the R package bimba (which has been uncoupled from the SMOTE step in the original implementation [32]).<br>Following pre-processing, the cleaned training data are subjected to various oversampling algorithms, including SMOTE, Borderline-SMOTE, Safe-Level-SMOTE, ADASYN, MWMOTE, and RWO, to balance classes. | 50% |
| **BL-SMOTE (NRAS)** | | | |
| **SL-SMOTE (NRAS)** | | | |
| **ADASYN (NRAS)** | | | |
| **MWMOTE (NRAS)** | | | |
| **RWO (NRAS)** | | | |
| **SMOTE-ENN** | **Over- & under-sampling hybrid** | Combines SMOTE [26] with Edited Nearest Neighbours (ENN)[33]. First, SMOTE synthesises new minority instances to balance the training data. Then, ENN is applied to remove instances whose class labels differ from the majority class of at least two of their three nearest neighbours. This process helps to eliminate noisy instances and create a smoother decision surface for classification. | 50% |
| **SMOTE-ENN (25%)** | | | 25% |
| **SMOTE-TL** | | Combines SMOTE [26] with the Tomek Links (TL) algorithm [34]. Initially, SMOTE is applied to balance the classes in the training data. Subsequently, the TL algorithm identifies pairs of instances with opposite classes that are each other's nearest neighbours. These pairs are then used to remove majority instances, clarifying the border between minority and majority instances and making the minority region more distinct. | 50% |
| **SMOTE- TL (25%)** | | | 25% |
| **SPW** | **Algorithm specific** | We tuned the scale_pos_weight parameter of the underlying lightGBM algorithm (R package lightgbm). This parameter defines the ratio of the negative class to the positive class, enabling the assignment of a configurable weight to the minority class. | 50% |

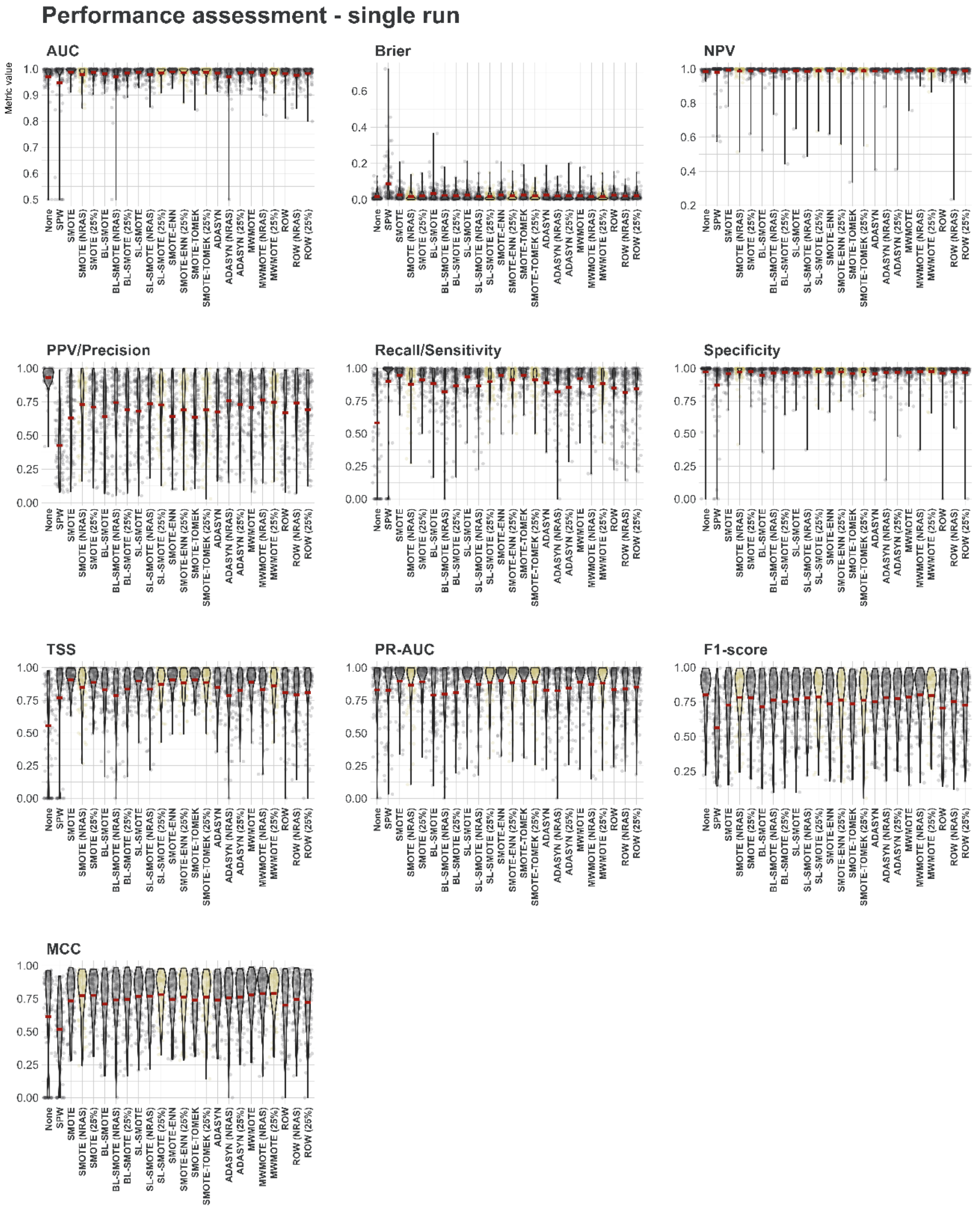


**Supplementary Figure 4 – Performance assessment of class balancing techniques over single held-out test set (run = 1) at >0.5 probability threshold**. Points represent results from individual transmission route/mode models (n=98). Violin plots show the kernel probability density of the data at different values. Yellow points and violin plots represent selected class balancing techniques, grey points and violin plots represent discarded class balancing techniques. Supplementary Table 9 provides full definitions of included performance metrics. Please note that for Brier score values closer to 0 indicate better performance, and those closer to 1 indicate worse performance.

### Supplementary Note 7 – Model training, optimisation, and validation

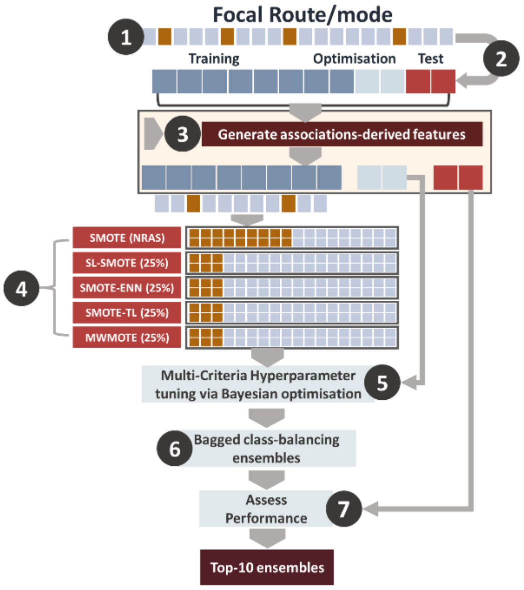


**Supplementary Figure 5 – Model training, optimisation, and validation over a single iteration.**

**Supplementary Table 7. LightGBM hyperparameters tuned per iteration.**

| Parameter | Range | |
|---|---|---|
| num_leaves | [7:4095] | Defines maximum number of leaves per weak learner (tree). Larger values increase accuracy on the training set but might lead to overfitting. |
| max_depth | [2:63] | Controls the maximum depth of each tree within the model, it is trained in tandem with num_leaves. Larger values increase accuracy on the training set but might lead to overfitting. |
| min_data_in_leaf | [200:10000] | Defines the minimum number of instances that must be contained in a leaf to be added to the tree. It controls for overfitting, so that the model does not become too specific. |
| Lambda_l1 | [0:100] | Regularisation parameters used to control overfitting. |
| Lambda_l2 | [0:100] | |
| min_gain_to_split | [0:15] | Regularisation parameter. When adding a new tree node, LightGBM chooses the split point that has the largest gain. Simply put, gain is the reduction in training loss that results from adding a split point. Larger values decrease training time. |
| min_sum_hessian_in_leaf | [0.01: \|train\|/1000] | The sum of hessians of the instances contained in the leaf. Hessian of a data point is the second order derivative of the loss function evaluated for each instance. It takes instance weights into consideration, as hessians are multiplied by those weights prior to passing them to the weak learner (tree). Tuning this hyperparameters allows weak learners to perform a more flexible split for those data instances with low confidence, and less flexible split for those with high confidence, thus providing an adaptive regularisation. |
| bagging_fraction | [0.4:1] | Controls the size of sample used in constructing each weak learner. For instance, a value = 0.7, means that each tree will be constructed using 70% of training data, randomly sampled without replacement. Lower values decrease training time. |
| feature_fraction | [0.4:1] | LightGBM randomly selects a subset of features per each tree (weak learner), feature_fraction defines % of feature included in each tree. For instance, when set to 0.4 LightGBM will sample 40% of features to be included in training each tree. This hyperparameter has two uses: speeding up training and avoiding overfitting. |

**Supplementary Table 8 - Measures utilised to assess the performance of our ensembles and their constituent models.** Bolded measures were used in ranking and selecting top 10 performing ensembles per route/mode.

| **Confusion matrix** | **Observed** | |
|---|---|---|
| **Predicted** | **Yes** | **No** |
| **Yes** | TP (true positives) | FP (False positives) |
| **No** | FN (False negatives) | TP (true negatives) |

| **Measure** | **Formula** | **Meaning** |
|---|---|---|
| Sensitivity (recall) – True Positive Rate (TPR) | $\frac{\text{TP}}{\text{TP} + \text{FN}}$ | Sensitivity is the percentage of actual positives (observed associations) that were correctly predicted. It indicates the percentage of 1s that was covered by the model. |
| False Positive Rate (FPR) | $\frac{\text{FP}}{\text{FP} + \text{TN}}$ | Percentage of false positive predictions of the model. |
| Specificity | $\frac{TN}{\text{TN} + \text{FP}}$ | Specificity is the percentage of negatives (here unknown associations, not necessarily true negative) that were correctly predicted |
| **<u>Precision (PPV (Positive Predictive Value))</u>** | $\frac{\text{TP}}{\text{TP} + \text{FP}}$ | Percentage of accurate positive predictions of the model. |
| NPV (Negative Predictive Value)) | $\frac{TN}{\text{TN} + \text{FN}}$ | Percentage of accurate negative predictions of the model. |
| **<u>AUC</u>** | **Area Under the ROC Curve** | Threshold-independent measure of model predictive performance that is commonly used as a validation metric for host-pathogen predictive models[23,35]. AUC favours both classes (negative/positive) equally. AUC captures how well the model separates the positive and negative examples and is calculated based on the TPR and FPR values. |
| TSS | $\text{Sensitivity} + \text{Specificity} - 1$ | Use of AUC has been criticised for its insensitivity to absolute predicted probability and its inclusion of a priori untenable prediction [36,37], we also calculated the True Skill Statistic (TSS)[38]. |
| **<u>PR-AUC</u>** | **Area Under the (precision-recall) Curve** | Threshold-independent measure, widely used with uneven class distribution. PR-AUC favours the positive (minority) class, and is calculated based on the TPR and PPV values. |
| F1-score | $2 \times \frac{\text{Precision} \times \text{Recall}}{\text{Precision} + \text{Recall}}$ | Captures the harmonic mean of the precision and recall. F1-score (F-score for short) is often used with uneven class distribution. |
| **<u>Brier</u>** | $\frac{1}{N}\sum_{i=1}^{N}(f_i - o_i)^2$ | Brier score measures the accuracy of probabilities generated by the models. It enables assessment of the confidence of a model (or ensemble |

| | | |
|---|---|---|
| | Where $N$ is the number of instances, $f_i$ is resulting probability, and $o_i$ the observed class of instance $i$ (negative =0, and positive = 1) | of models). A more confident model would generate probabilities closer to 1 for instances of the positive class, and closer to 0 for instances of the negative class.<br>The value of Briers sores ranges between [0,+1], with 0 being the best value (perfect classification), and 1 the worst value (perfect misclassification). |
| Matthews Correlation Coefficient (MCC) | $$\frac{(TP \times TN) - (FP \times FN)}{\sqrt{(TP + FP) \times (TP + FN) \times (TN + FP) \times (TN + FN)}}$$ | MCC is a special case of the Pearson Correlation Coefficient and measures the correlation of the true classes with the predicted labels. MCC ranges between [−1,+1], with –1 meaning perfect misclassification and +1 perfect classification. MCC produces a high score only when good results are obtained in all categories of the confusion matrix (TP, FP, TN, FN), proportionally both to the size of positive and negative class. |

### Supplementary References


1. Federhen S. The NCBI Taxonomy database. Nucleic Acids Res. 2012;40: D136-43. doi:10.1093/nar/gkr1178
2. Hu Y, Zandi R, Anavitarte A, Knobler CM, Gelbart WM. Packaging of a Polymer by a Viral Capsid: The Interplay between Polymer Length and Capsid Size. Biophys J. 2008;94: 1428. doi:10.1529/BIOPHYSJ.107.117473
3. Galtier N, Lobry JR. Relationships Between Genomic G+C Content, RNA Secondary Structures, and Optimal Growth Temperature in Prokaryotes. J Mol Evol 1997 446. 1997;44: 632–636. doi:10.1007/PL00006186
4. Sanjuán R, Nebot MR, Chirico N, Louis M, Belshaw R, Sanjua R, et al. Viral Mutation Rates Viral Mutation Rates □. J Virol. 2010;84: 9733–9748. doi:10.1128/JVI.00694-10
5. Coffin JM. Structure and Classification of Retroviruses. The Retroviridae. Springer US; 1992. pp. 19–49. doi:10.1007/978-1-4615-3372-6_2
6. Nisole S, Saïb A. Early steps of retrovirus replicative cycle. Retrovirology. 2004. doi:10.1186/1742-4690-1-9
7. Hulo C, De Castro E, Masson P, Bougueleret L, Bairoch A, Xenarios I, et al. ViralZone: A knowledge resource to understand virus diversity. Nucleic Acids Res. 2011;39: D576. doi:10.1093/nar/gkq901
8. Lefkowitz EJ, Dempsey DM, Hendrickson RC, Orton RJ, Siddell SG, Smith DB. Virus taxonomy: The database of the International Committee on Taxonomy of Viruses (ICTV). Nucleic Acids Res. 2018;46: D708–D717. doi:10.1093/nar/gkx932
9. Wawrzyniak P, Plucienniczak G, Bartosik D. The different faces of rolling-circle replication and its multifunctional initiator proteins. Frontiers in Microbiology. Frontiers Media S.A.; 2017. doi:10.3389/fmicb.2017.02353
10. Malathi VG, Renuka Devi P. ssDNA viruses: key players in global virome. VirusDisease. 2019;30: 3. doi:10.1007/S13337-019-00519-4
11. Lin X, Eddy NR, Noel JK, Whitford PC, Wang Q, Ma J, et al. Order and disorder control the functional rearrangement of influenza hemagglutinin. Proc Natl Acad Sci U S A. 2014;111: 12049–54. doi:10.1073/pnas.1412849111
12. Sicard A, Michalakis Y, Gutiérrez S, Blanc S. The Strange Lifestyle of Multipartite Viruses. PLoS Pathogens. Public Library of Science; 2016. doi:10.1371/journal.ppat.1005819
13. Rey FA, Lok SM. Common Features of Enveloped Viruses and Implications for Immunogen Design for Next-Generation Vaccines. Cell. Cell Press; 2018. pp. 1319–1334. doi:10.1016/j.cell.2018.02.054
14. Bushman FD, McCormick K, Sherrill-Mix S. Virus structures constrain transmission modes. Nat Microbiol. 2019;4: 1778–1780. doi:10.1038/S41564-019-0523-5
15. Singer GAC, Hickey DA. Nucleotide Bias Causes a Genomewide Bias in the Amino Acid Composition of Proteins. Mol Biol Evol. 2000;17: 1581–1588.

doi:10.1093/OXFORDJOURNALS.MOLBEV.A026257
16. Kunec D, Osterrieder N. Codon Pair Bias Is a Direct Consequence of Dinucleotide Bias. Cell Rep. 2016;14: 55–67. doi:10.1016/J.CELREP.2015.12.011
17. Giallonardo F Di, Schlub TE, Shi M, Holmes EC. Dinucleotide Composition in Animal RNA Viruses Is Shaped More by Virus Family than by Host Species. J Virol. 2017;91. doi:10.1128/JVI.02381-16
18. Quax TEF, Claassens NJ, Söll D, van der Oost J. Codon Bias as a Means to Fine-Tune Gene Expression. Mol Cell. 2015;59: 149–161. doi:10.1016/J.MOLCEL.2015.05.035
19. Groenke N, Trimpert J, Merz S, Conradie AM, Wyler E, Zhang H, et al. Mechanism of Virus Attenuation by Codon Pair Deoptimization. Cell Rep. 2020;31. doi:10.1016/J.CELREP.2020.107586
20. Plotkin JB, Robins H, Levine AJ. Tissue-specific codon usage and the expression of human genes. Proc Natl Acad Sci U S A. 2004;101: 12588. doi:10.1073/PNAS.0404957101
21. Creighton TH. “Chapter 1”. Proteins: structures and molecular properties. San Francisco: W. H. Freeman; 1993.
22. Wright ES. DECIPHER: Harnessing local sequence context to improve protein multiple sequence alignment. BMC Bioinformatics. 2015;16: 322. doi:10.1186/s12859-015-0749-z
23. Babayan SA, Orton RJ, Streicker DG. Predicting reservoir hosts and arthropod vectors from evolutionary signatures in RNA virus genomes. Science (80- ). 2018;362: 577–580. doi:10.1126/science.aap9072
24. Sharp PM, Li WH. The codon Adaptation Index--a measure of directional synonymous codon usage bias, and its potential applications. Nucleic Acids Res. 1987;15: 1281. doi:10.1093/NAR/15.3.1281
25. Kumar S, Suleski M, Craig JM, Kasprowicz AE, Sanderford M, Li M, et al. TimeTree 5: An Expanded Resource for Species Divergence Times. Mol Biol Evol. 2022;39. doi:10.1093/MOLBEV/MSAC174
26. Chawla N V, Bowyer KW, Hall LO, Kegelmeyer WP. SMOTE: Synthetic Minority Over-sampling Technique. J Artif Intell Res. 2002. Available: https://arxiv.org/pdf/1106.1813.pdf
27. Han H, Wang WY, Mao BH. Borderline-SMOTE: A new over-sampling method in imbalanced data sets learning. Lect Notes Comput Sci. 2005;3644: 878–887. doi:10.1007/11538059_91
28. Bunkhumpornpat C, Sinapiromsaran K, Lursinsap C. Safe-level-SMOTE: Safe-level-synthetic minority over-sampling technique for handling the class imbalanced problem. Lect Notes Comput Sci (including Subser Lect Notes Artif Intell Lect Notes Bioinformatics). 2009;5476 LNAI: 475–482. doi:10.1007/978-3-642-01307-2_43/COVER
29. He H, Bai Y, Garcia EA, Li S. ADASYN: Adaptive synthetic sampling approach for imbalanced learning. Proc Int Jt Conf Neural Networks. 2008; 1322–1328. doi:10.1109/IJCNN.2008.4633969
30. Barua S, Islam MM, Yao X, Murase K. MWMOTE - Majority weighted minority oversampling technique for imbalanced data set learning. IEEE Trans Knowl Data Eng. 2014;26: 405–425. doi:10.1109/TKDE.2012.232
31. Zhang H, Li M. RWO-Sampling: A random walk over-sampling approach to imbalanced data classification. Inf Fusion. 2014;20: 99–116. doi:10.1016/J.INFFUS.2013.12.003
32. Rivera WA. Noise Reduction A Priori Synthetic Over-Sampling for class imbalanced data sets. Inf Sci (Ny). 2017;408: 146–161. doi:10.1016/J.INS.2017.04.046
33. Wilson DL. Asymptotic Properties of Nearest Neighbor Rules Using Edited Data. IEEE Trans Syst Man Cybern. 1972;2: 408–421. doi:10.1109/TSMC.1972.4309137
34. Tomek I. EXPERIMENT WITH THE EDITED NEAREST-NEIGHBOR RULE. IEEE Trans Syst Man Cybern. 1976;SMC-6: 448–452. doi:10.1109/TSMC.1976.4309523
35. Dallas T, Park AW, Drake JM. Predicting cryptic links in host-parasite networks. Koella J, editor. PLOS Comput Biol. 2017;13: e1005557. doi:10.1371/journal.pcbi.1005557
36. Lobo JM, Jiménez-Valverde A, Real R. AUC: a misleading measure of the performance of predictive distribution models. Glob Ecol Biogeogr. 2008;17: 145–151. doi:10.1111/j.1466-8238.2007.00358.x
37. Allen T, Murray KA, Zambrana-Torrelio C, Morse SS, Rondinini C, Di Marco M, et al. Global hotspots and correlates of emerging zoonotic diseases. Nat Commun. 2017;8: 1124. doi:10.1038/s41467-017-00923-8
38. Barbet-Massin M, Jiguet F, Albert CH, Thuiller W. Selecting pseudo-absences for species distribution models: how, where and how many? Methods Ecol Evol. 2012;3: 327–338. doi:10.1111/j.2041-210X.2011.00172.x

### Supplementary Results 1 – Transmission by multiple unique pathways

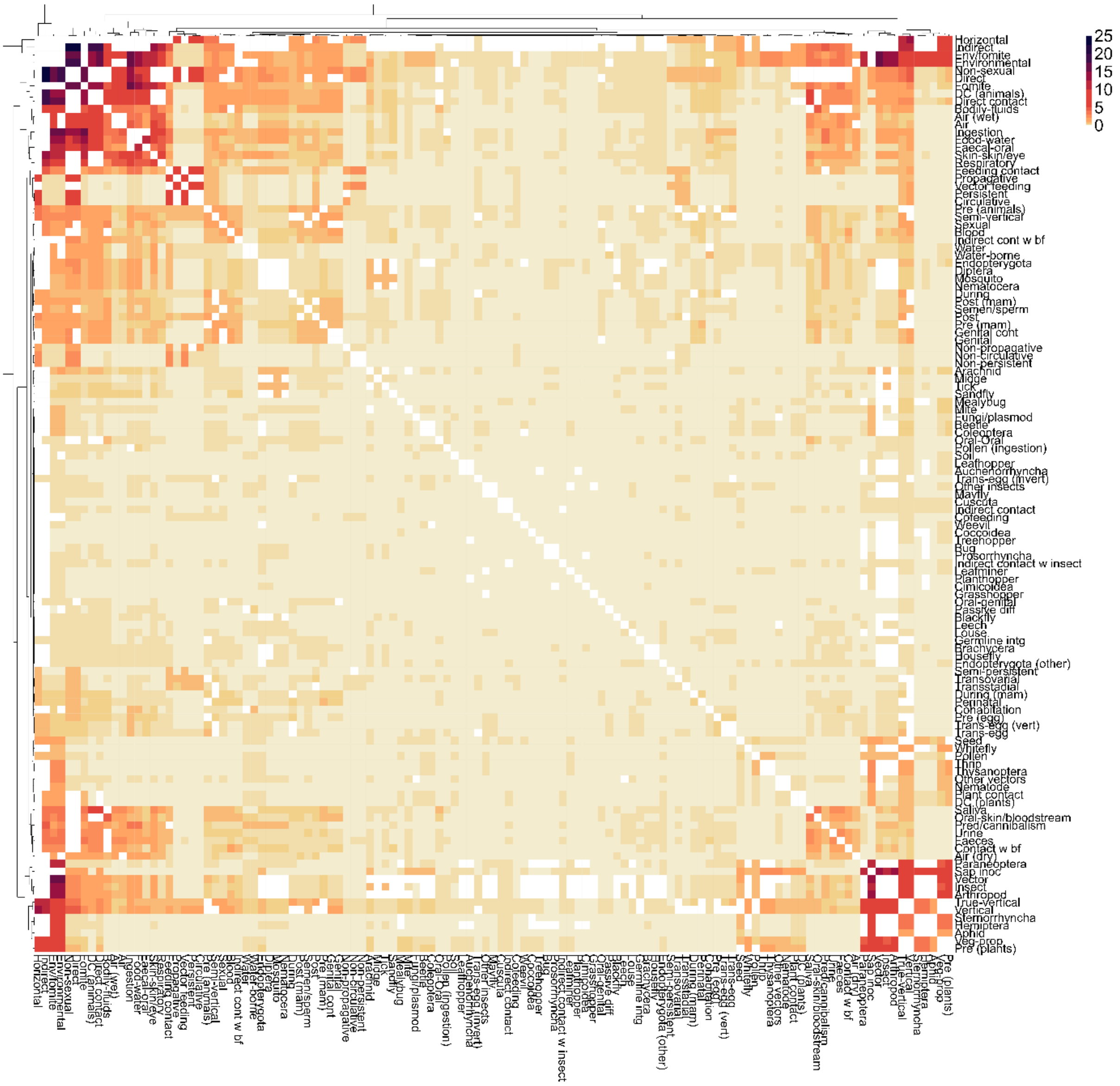


**Supplementary Figure 6 – Viruses known to be transmitted by unique pathway pairs to the same host species.** Rows and column represent the transmission pathways identified in this study (**n=120**). Heatmap represent the percent of virus-host associations (of total = 24,953) observed to be transmitted by each unique pathway pair. Unique pathway pairs are defined as any pair of nodes in our hierarchy (Figure 1), whereby no one node of the pair is a direct ancestor of the other. We performed hierarchical clustering on both rows and columns, using the R package pheatmap. the resulting dendrogram is displayed (top and left). White cells represent NAs = non-unique pathways.

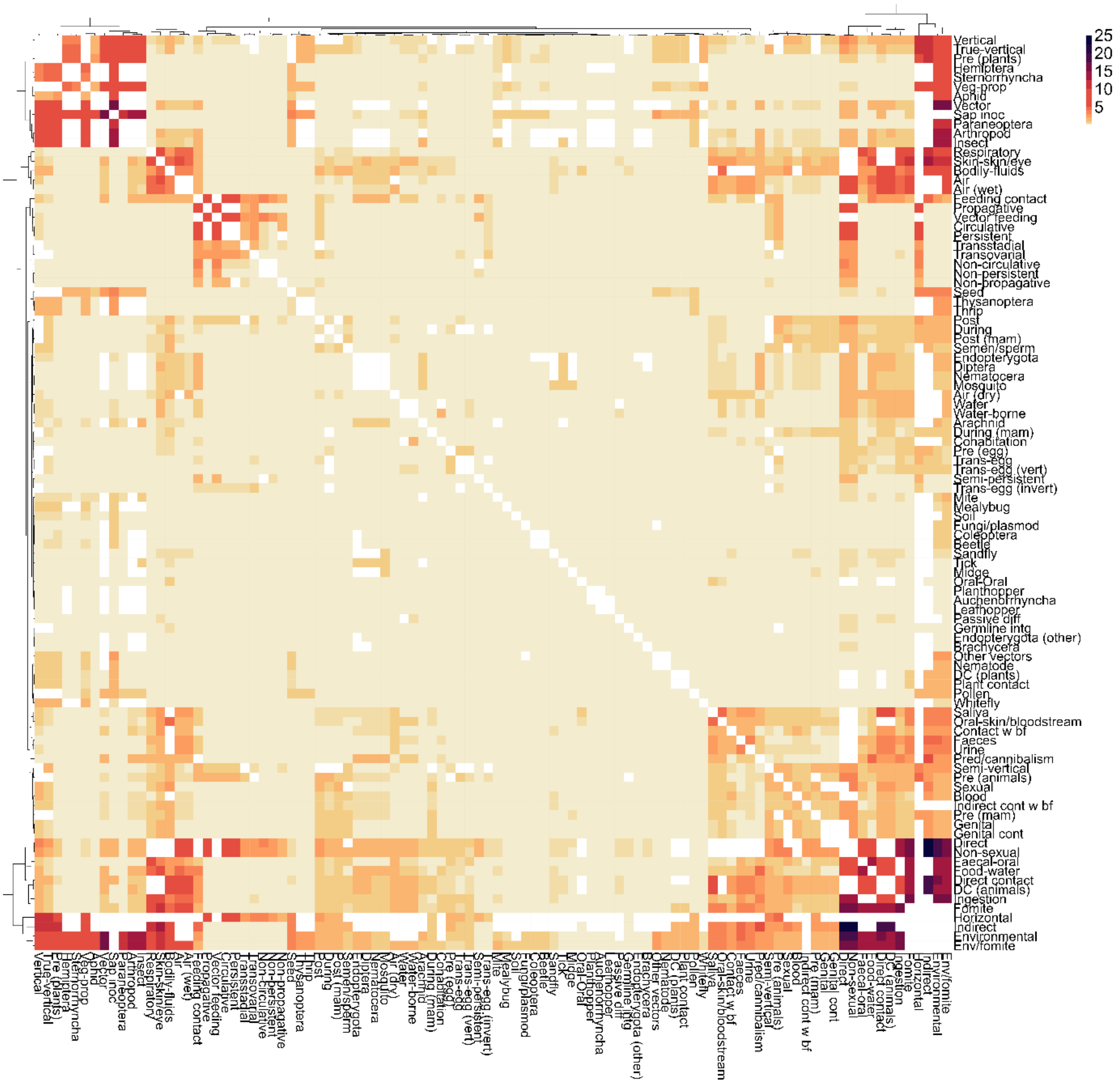


**Supplementary Figure 7 – Viruses predicted (within-sample) to be transmitted by unique pathway pairs to the same host species.** Rows and column represent transmission pathways modelled in this study (n=98). Heatmap represent the percent of virus-host associations predicted (mean top-10 ensemble probability >0.5, for each route/mode) to be transmitted by each unique pathway pair, from within sample (associations used to train and test our models, for which at least one known route was identified, and for which our models have made at least one positive prediction, n=24,862). Unique pathway pairs are defined as any pair of nodes in our hierarchy (Figure 1), whereby no one node of the pair is a direct ancestor of the other. We performed hierarchical clustering on both rows and columns, using the R package `pheatmap`. the resulting dendrogram is displayed (top and left). White cells represent NAs = non-unique pathways.

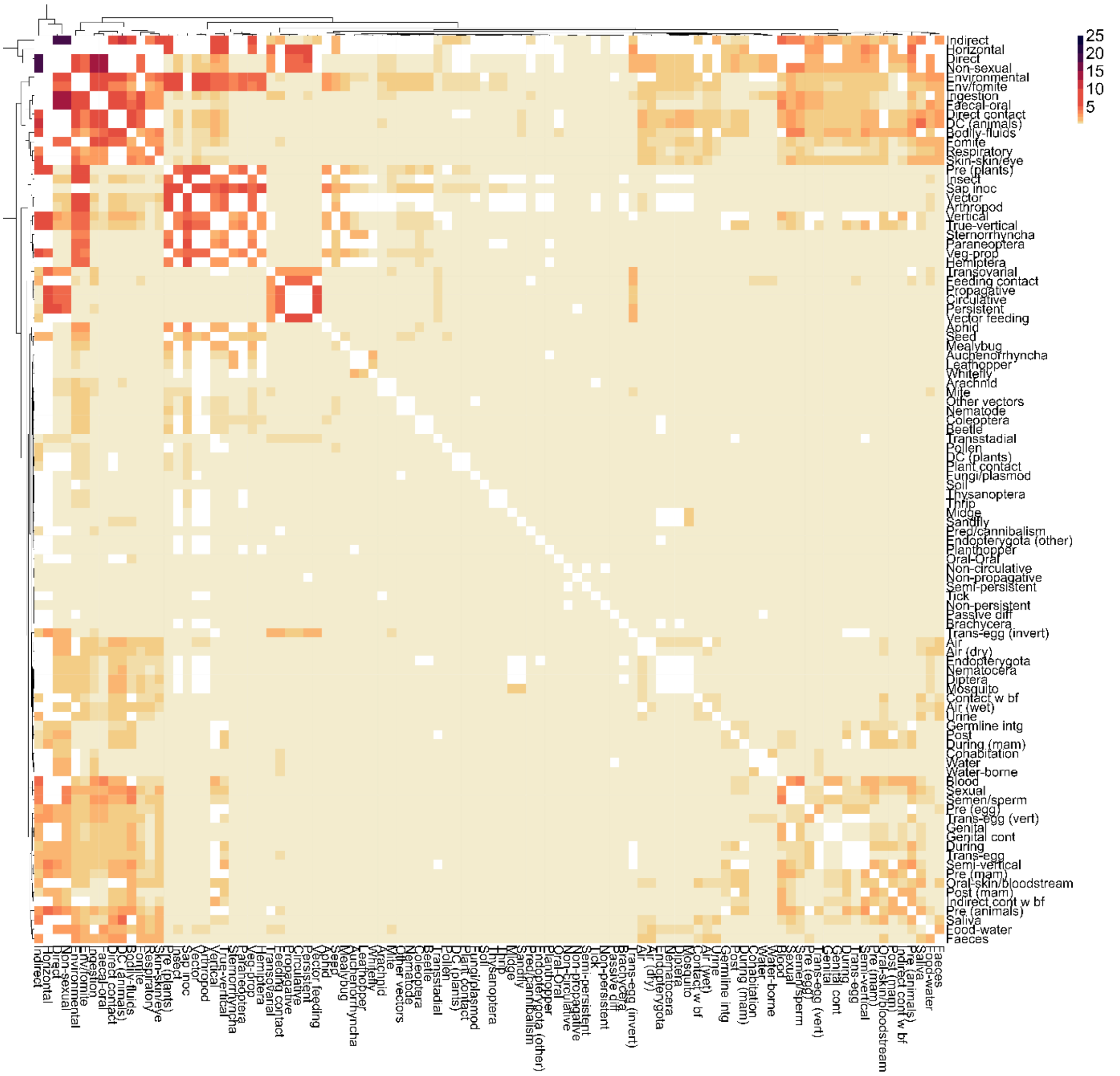


**Supplementary Figure 8 – Viruses predicted (out-of-sample) to be transmitted by unique pathway pairs to the same host species.** Rows and column represent the routes/modes modelled in this study (n=98). Heatmap represent the percent of virus-host associations predicted (mean top-10 ensemble probability >0.5, for each route/mode) to be transmitted by each unique pathway pair from out-of-sample associations (virus-host associations without known routes, did not enter our training and testing pipelines, with at least one route/mode predicted by our models, n=3,108). Unique pathway pairs are defined as any pair of nodes in our hierarchy (Figure 1), whereby no one node of the pair is a direct ancestor of the other. We performed hierarchical clustering on both rows and columns, using the R package pheatmap. the resulting dendrogram is displayed (top and left). White cells represent NAs = non-unique pathways.

### Supplementary Results 2 – Instance-level feature-contribution

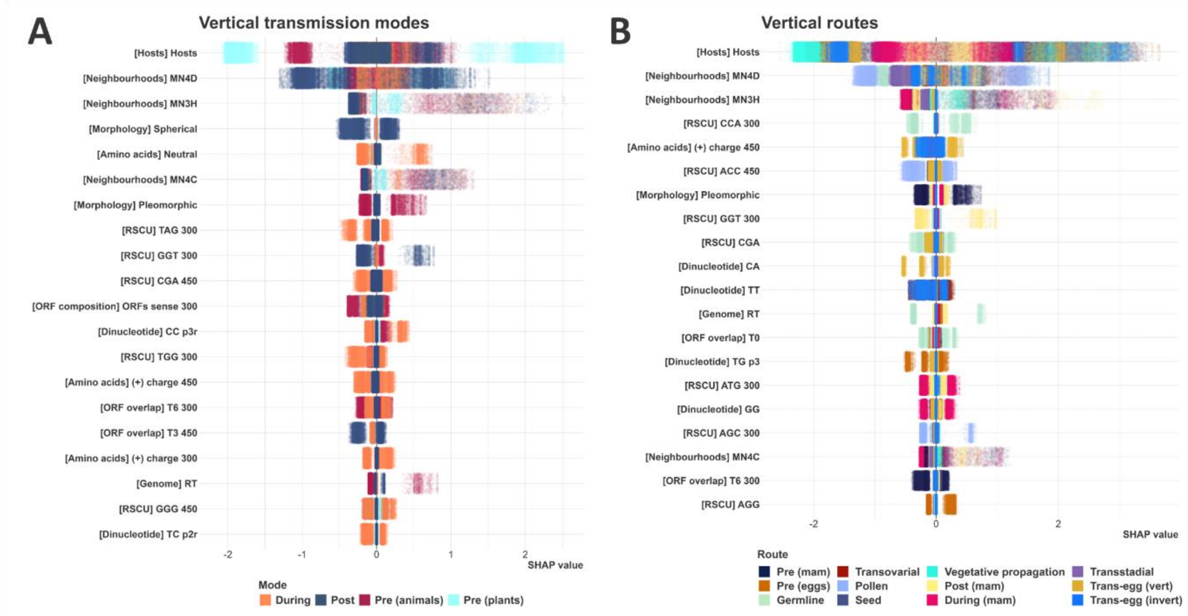


**Supplementary Figure 9 – Instance-level feature-contribution to vertical transmission routes/modes.** We averaged instance-level SHAP values generated by all constituent models of each of our top-10 ensembles (50 models per each included route/mode). In each sub-plot, features were ordered by the spread of their variance (max(variance)-min(variance) across all routes/modes included in each sub-plot), and the top 20 features (from most to least spread) were selected. Points represent virus-host associations (instances), and are coloured by the underlying route/mode. The Y-axes represent the selected features (category of each feature between brackets). The X-axes represent SHAP values. Positive SHAP values indicate that the feature has contributed towards a positive prediction for the instance (the virus is transmitted to the host species via route/mode). Negative SHAP values indicate that the feature has contributed towards a negative prediction (the virus is not transmitted to the host via route/mode). Larger magnitudes indicate that the feature has a stronger influence on the prediction for the given instance.

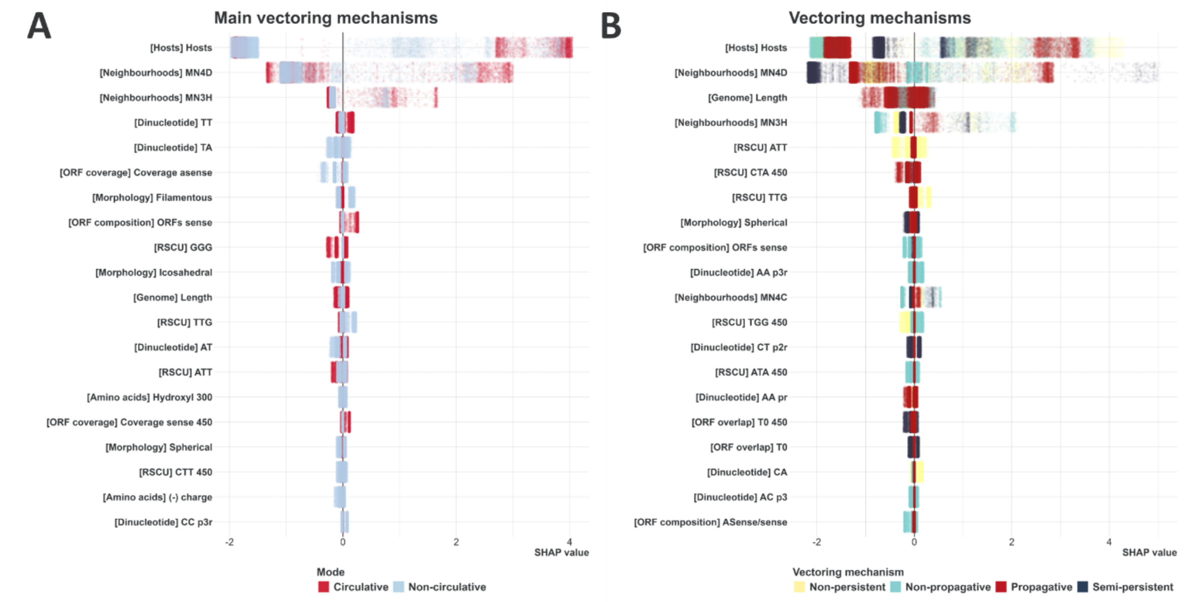


**Supplementary Figure 10 – Instance-level feature-contribution to vectoring-mechanisms.** We averaged instance-level SHAP values generated by all constituent models of each of our top-10 ensembles (50 models per each included vectoring mechanism). In each sub-plot, features were ordered by the spread of their variance (max(variance)-min(variance) across all vectoring mechanisms included in each sub-plot), and the top 20 features (from most to least spread) were selected. Points represent virus-host associations (instances), and are coloured by the underlying route/mode. The Y-axes represent the selected features (category of each feature between brackets). The X-axes represent SHAP values. Positive SHAP values indicate that the feature has contributed towards a positive prediction for the instance (the virus is transmitted to the host species via route/mode). Negative SHAP values indicate that the feature has contributed towards a negative prediction (the virus is not transmitted to the host via route/mode). Larger magnitudes indicate that the feature has a stronger influence on the prediction for the given instance.

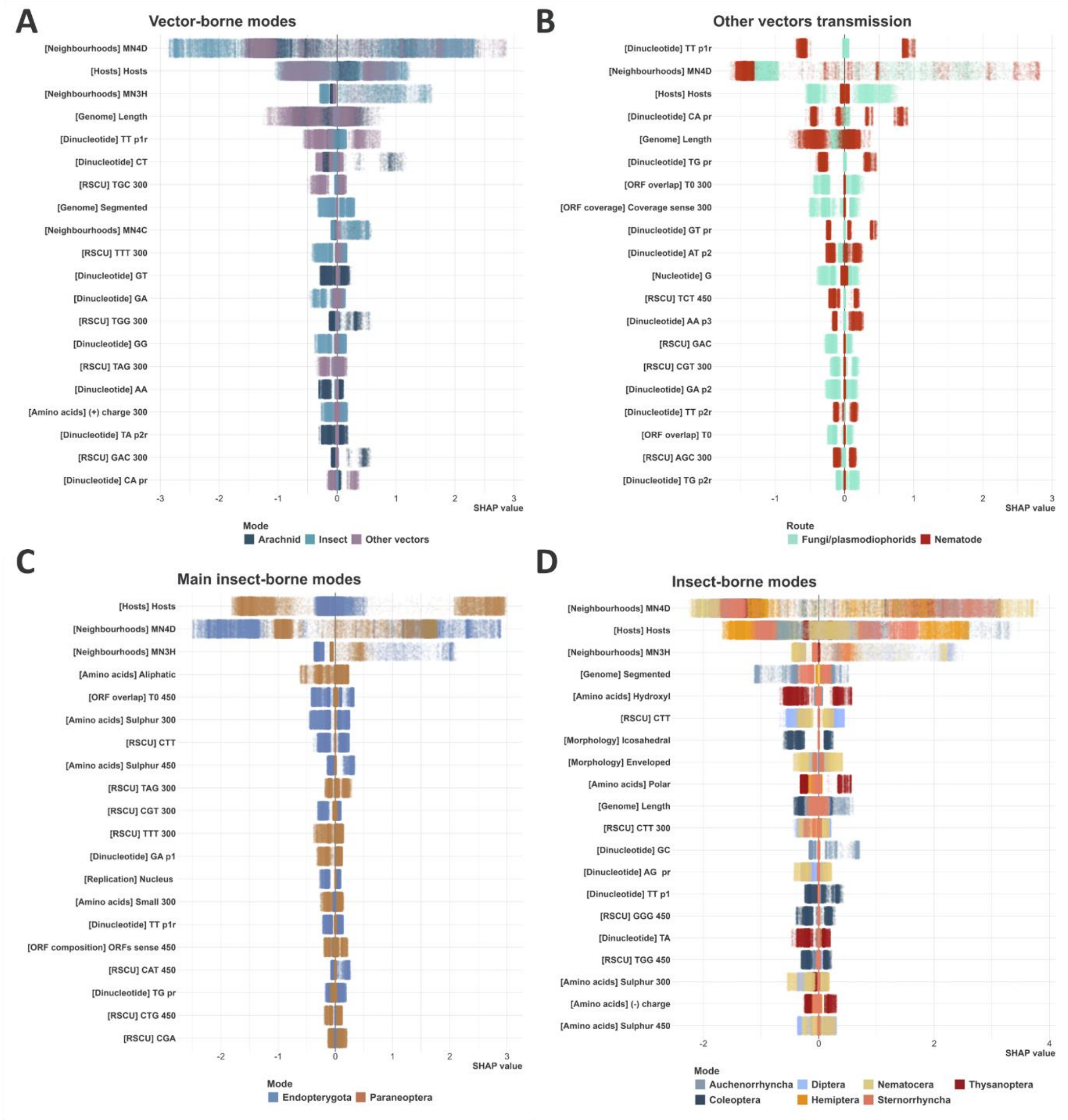


**Supplementary Figure 11 – Instance-level feature-contribution to vector-borne transmission routes/modes.** We averaged instance-level SHAP values generated by all constituent models of each of our top-10 ensembles (50 models per each included route/mode). In each sub-plot, features were ordered by the spread of their variance (max(variance)-min(variance) across all routes/modes included in each sub-plot), and the top 20 features (from most to least spread) were selected. Points represent virus-host associations (instances), and are coloured by the underlying route/mode. The Y-axes represent the selected features (category of each feature between brackets). The X-axes represent SHAP values. Positive SHAP values indicate that the feature has contributed towards a positive prediction for the instance (the virus is transmitted to the host species via route/mode). Negative SHAP values indicate that the feature has contributed towards a negative prediction (the virus is not transmitted to the host via route/mode). Larger magnitudes indicate that the feature has a stronger influence on the prediction for the given instance.

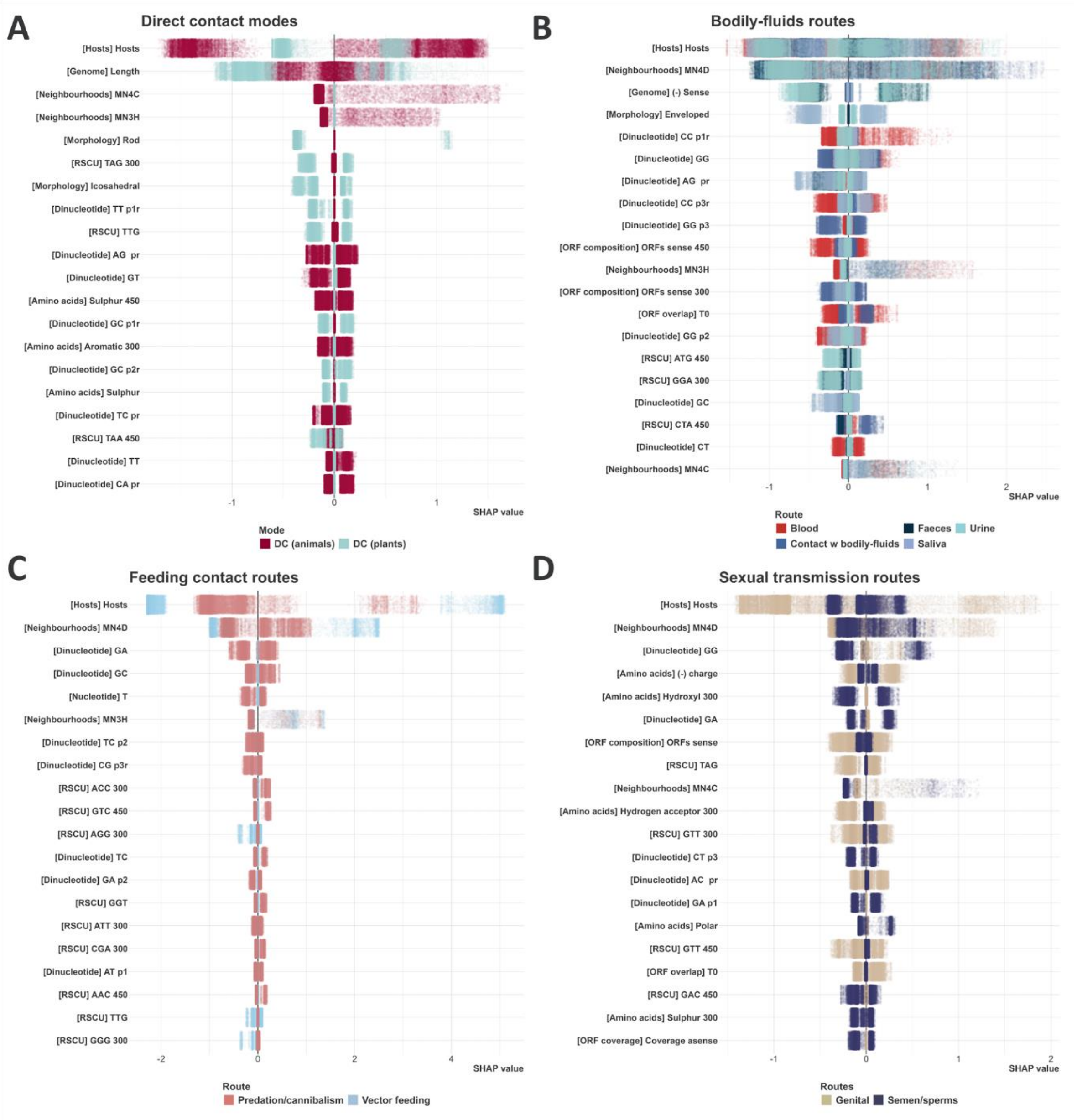


**Supplementary Figure 12 – Instance-level feature-contribution to remainder direct transmission routes/modes.** We averaged instance-level SHAP values generated by all constituent models of each of our top-10 ensembles (50 models per each included route/mode). In each sub-plot, features were ordered by the spread of their variance (max(variance)-min(variance) across all routes/modes included in each sub-plot), and the top 20 features (from most to least spread) were selected. Points represent virus-host associations (instances), and are coloured by the underlying route/mode. The Y-axes represent the selected features (category of each feature between brackets). The X-axes represent SHAP values. Positive SHAP values indicate that the feature has contributed towards a positive prediction for the instance (the virus is transmitted to the host species via route/mode). Negative SHAP values indicate that the feature has contributed towards a negative prediction (the virus is not transmitted to the host via route/mode). Larger magnitudes indicate that the feature has a stronger influence on the prediction for the given instance.

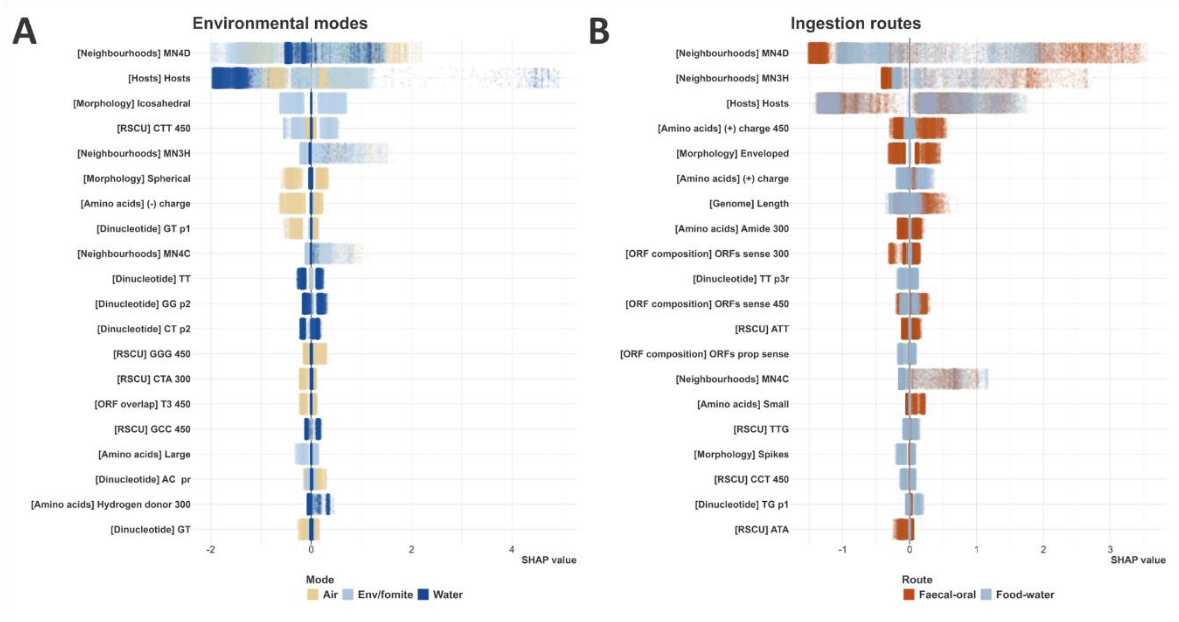


**Supplementary Figure 13 – Instance-level feature-contribution to remainder indirect transmission routes/modes.** We averaged instance-level SHAP values generated by all constituent models of each of our top-10 ensembles (50 models per each included route/mode). In each sub-plot, features were ordered by the spread of their variance (max(variance)-min(variance) across all routes/modes included in each sub-plot), and the top 20 features (from most to least spread) were selected. Points represent virus-host associations (instances), and are coloured by the underlying route/mode. The Y-axes represent the selected features (category of each feature between brackets). The X-axes represent SHAP values. Positive SHAP values indicate that the feature has contributed towards a positive prediction for the instance (the virus is transmitted to the host species via route/mode). Negative SHAP values indicate that the feature has contributed towards a negative prediction (the virus is not transmitted to the host via route/mode). Larger magnitudes indicate that the feature has a stronger influence on the prediction for the given instance.

### Supplementary Results 3 – Stability of SHAP Values

SHAP stability analysis examines the consistency of SHAP values across different subsets of data and/or model iterations, thus providing insights into the reliability of resulting interpretations, particularly in scenarios with highly correlated features. When features exhibit high correlation, there is some risk that SHAP values may vary significantly depending on the subset of data or model iteration, leading to inconsistent interpretations.

We quantified SHAP stability by computing the absolute differences between SHAP values for each feature across different subsets of training data (per each run of our top-10 ensembles, total runs = 10, total repeats per run = 10) and model iterations (5 class-balancing techniques). The absolute differences are then averaged across all features to compute the average absolute difference. A lower average absolute difference suggests higher stability and greater consistency in SHAP values, indicating more consistent model interpretations, whereas a higher value implies greater variability in SHAP values, and may highlight potential challenges in accurately attributing model predictions to individual features.

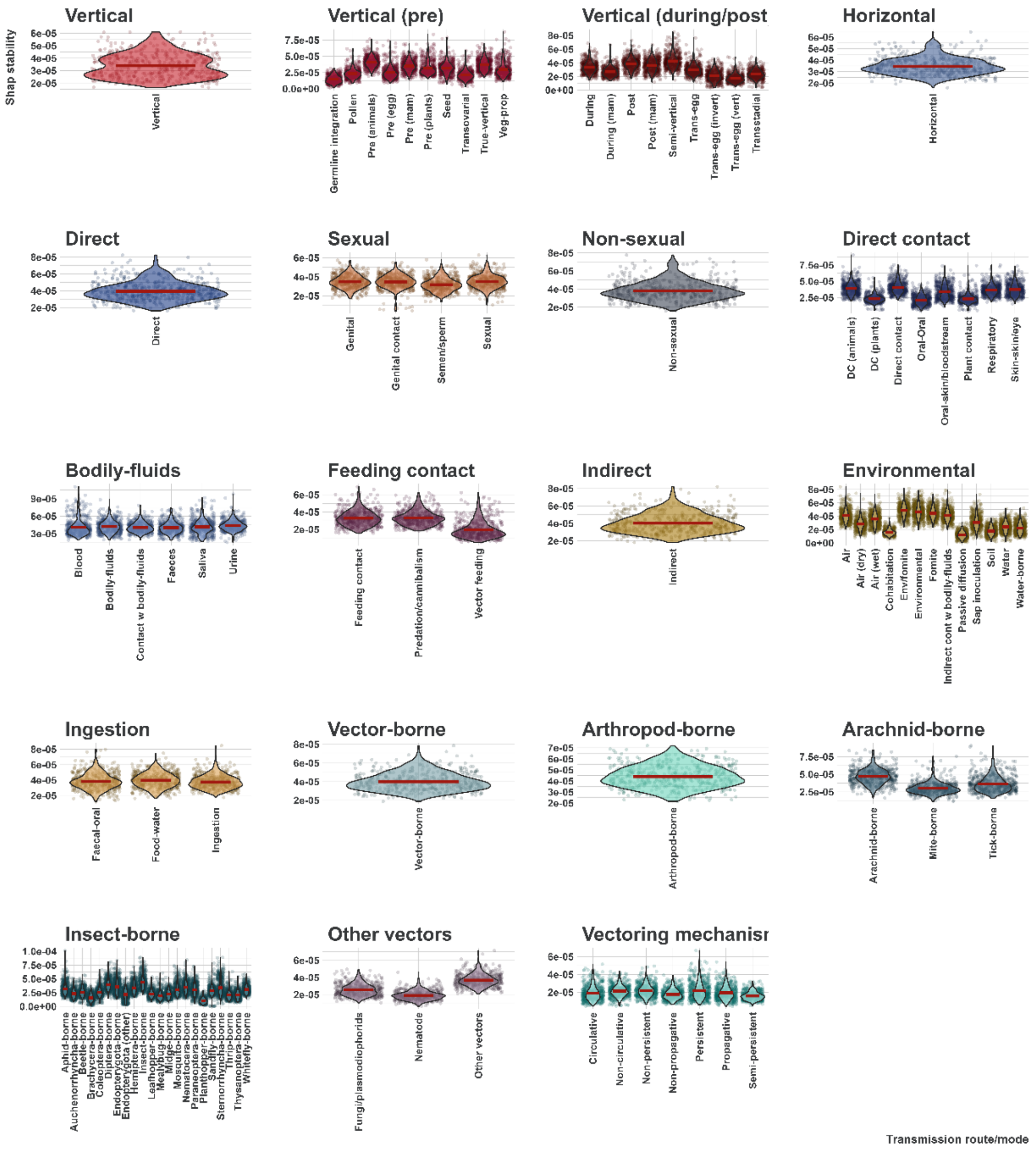


**Supplementary Figure 14 – Stability of SHAP values of our top-10 ensembles.** Points indicate average absolute differences in SHAP values (500 points per route/mode). Violin plots indicate the kernel probability density of the data at different values. Red horizontal lines indicate the mean average absolute difference in SHAP values per each route/mode. Routes/modes are categorised into 19 categories. Points and violin plots are coloured by category of transmission.

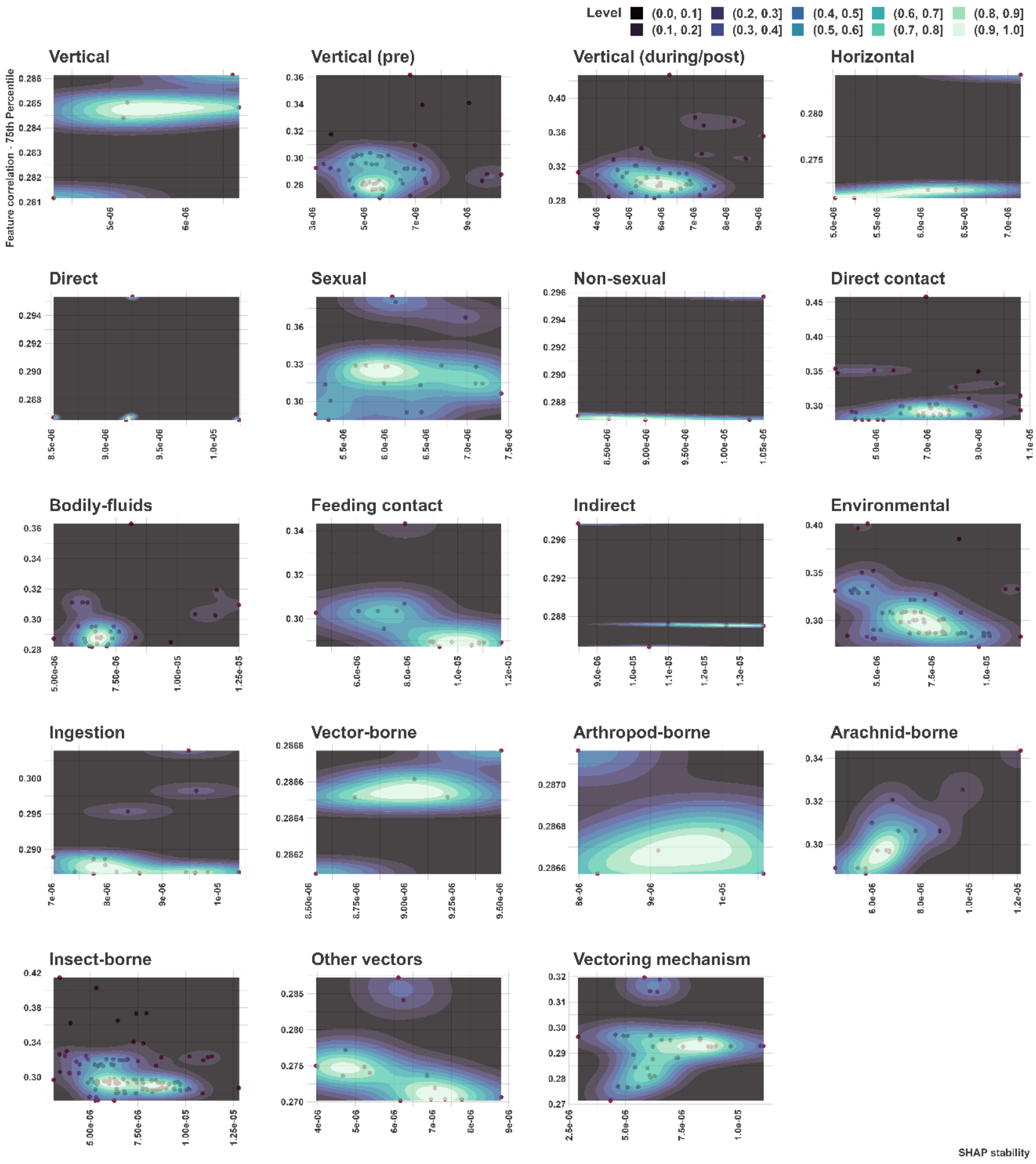


**Supplementary Figure 15 – Joint distribution of SHAP stability and third quartile of feature correlation values of our top-10 ensembles.** Density plots illustrates the joint distribution of mean SHAP stability (average absolute differences in SHAP values) and the third quantile of correlation values between features across different iterations (n=10), and models (n=5 per route/mode, 98 routes/modes in total). Routes/modes are categorised into 19 categories. Darker regions indicate lower density, while lighter regions suggest higher density. Dense regions in the lower left quadrant suggest robust interpretations. Dense regions in the upper right quadrant may suggest challenges in achieving stable SHAP values with high feature correlation.

### Supplementary Results 4 – Post-hoc analysis of prediction dependencies

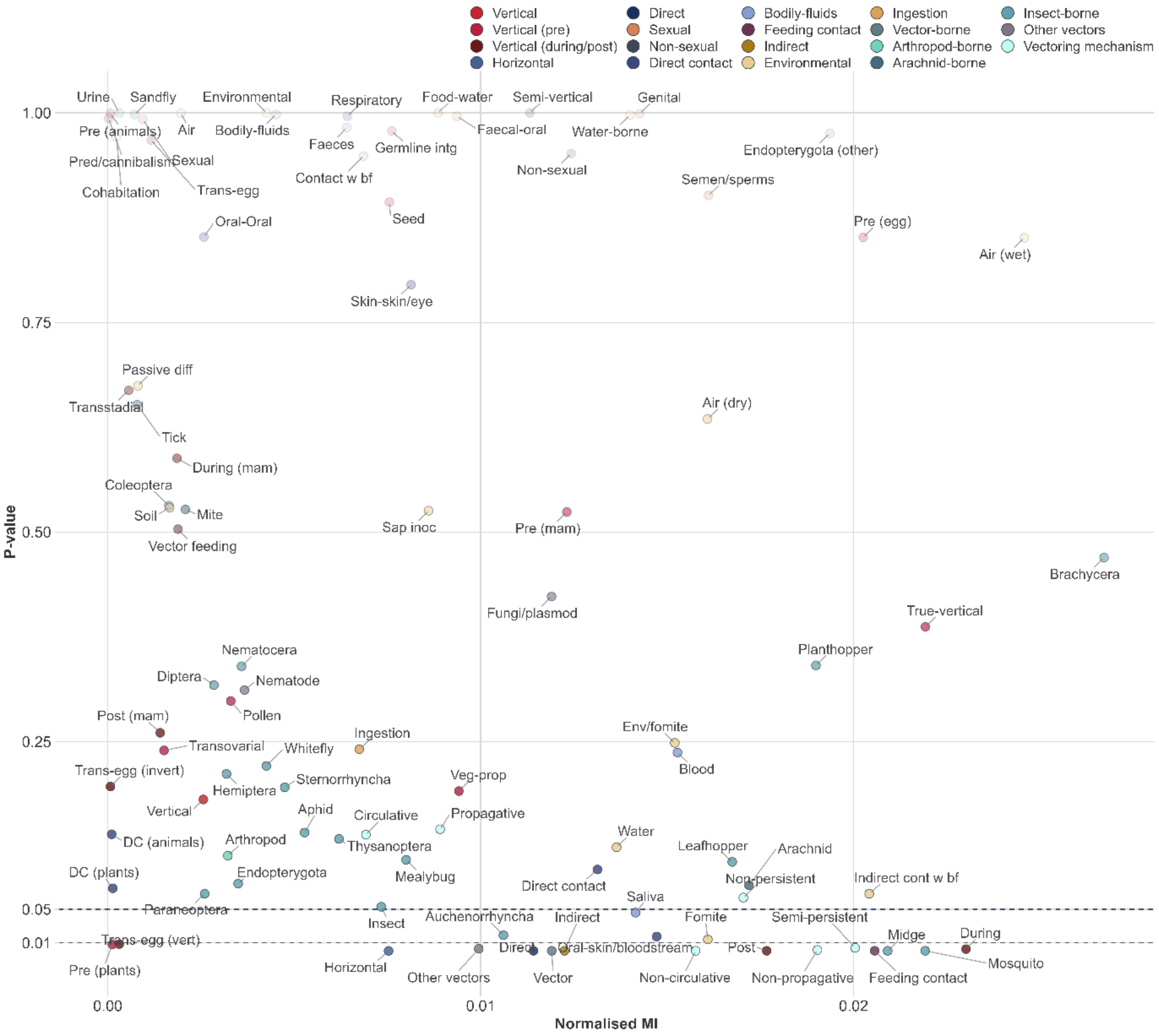


**Supplementary Figure 16 – Dependencies between predicted routes/modes and probabilities of their siblings.** Points represent virus transmission routes/modes modelled in this study (Supplementary Table 3). Points are coloured by transmission mode. X axis represents the normalised MI estimates computed between the mean probabilities (top-10 ensembles) of instances predicted for the focal route/mode (instances with mean probability >0.5), and the corresponding mean probabilities obtained for the corresponding siblings. Y axis represents the p-value obtained for each MI estimate via bootstrapping (n=2,000).

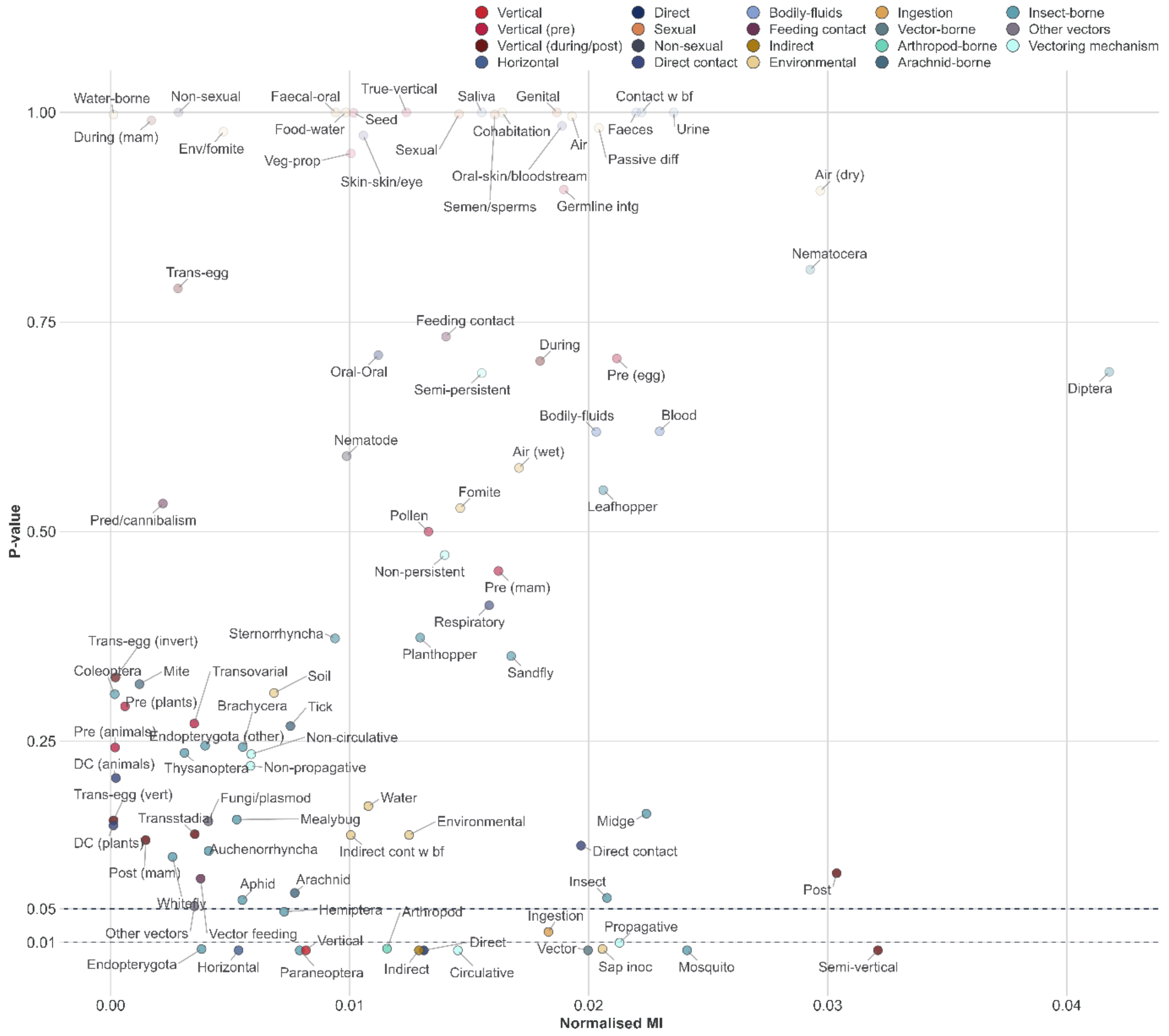


**Supplementary Figure 17 – Dependencies between probabilities for routes/modes and knowledge of their siblings.** Points represent virus transmission routes/modes modelled in this study (Supplementary Table 3). Points are coloured by transmission mode. X axis represents the normalised MI estimates computed between the mean probabilities (top-10 ensembles) produced for the focal route/mode, and the corresponding mean probabilities obtained for the siblings, for virus-host instances observed for at least one of the siblings. Y axis represents the p-value obtained for each MI estimate via bootstrapping (n=2,000).

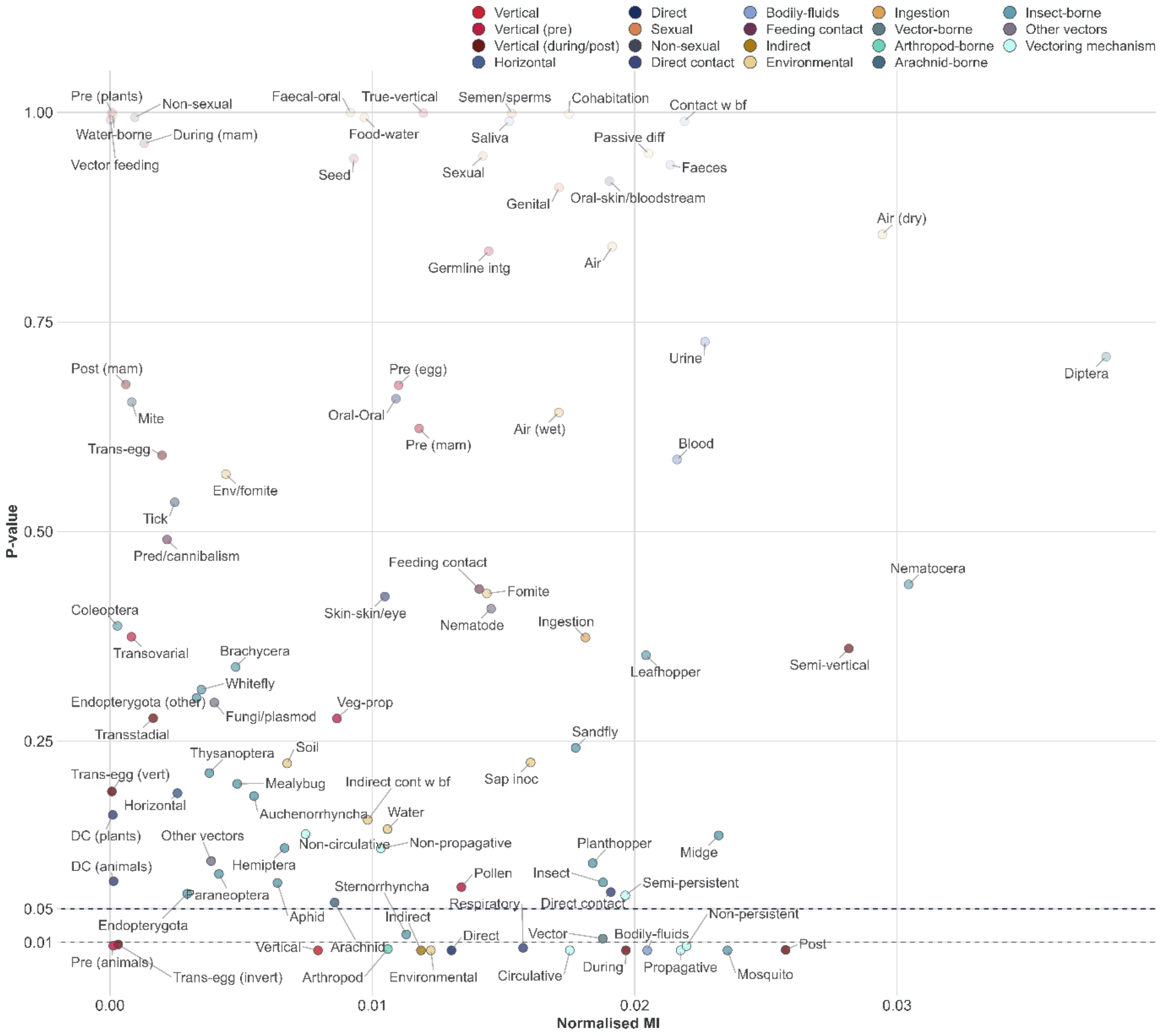


**Supplementary Figure 18 – Dependencies between probabilities for routes/modes and predictions of their siblings.** Points represent virus transmission routes/modes modelled in this study (Supplementary Table 3). Points are coloured by transmission mode. X axis represents the normalised MI estimates computed between the mean probabilities (top-10 ensembles) produced for the focal route/mode, and the corresponding mean probabilities obtained for the siblings, for virus-host instances predicted (mean probability >0.5) for at least one of the siblings. Y axis represents the p-value obtained for each MI estimate via bootstrapping (n=2,000).

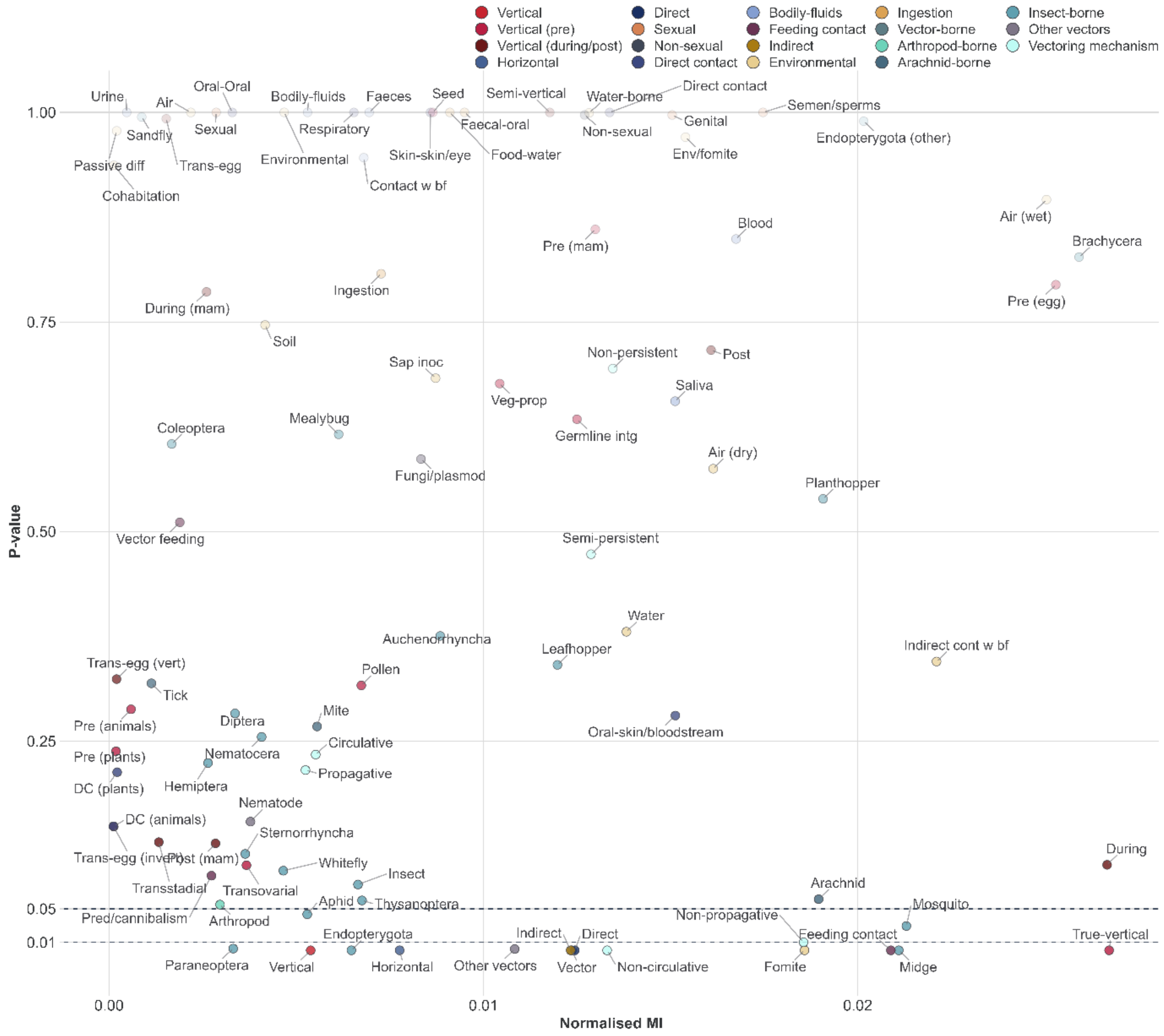


**Supplementary Figure 19 – Dependencies between knowledge of routes/modes and resulting probabilities of their siblings.** Points represent virus transmission routes/modes modelled in this study (Supplementary Table 3). Points are coloured by transmission mode. X axis represents the normalised MI estimates computed between the mean probabilities (top-10 ensembles) of instances produced for the focal route/mode, for virus-host instances observed to be transmitted via the focal route/mode, and the corresponding mean probabilities obtained from the siblings. Y axis represents the p-value obtained for each MI estimate via bootstrapping (n=2,000).

### Supplementary Results 5 – Performance assessment

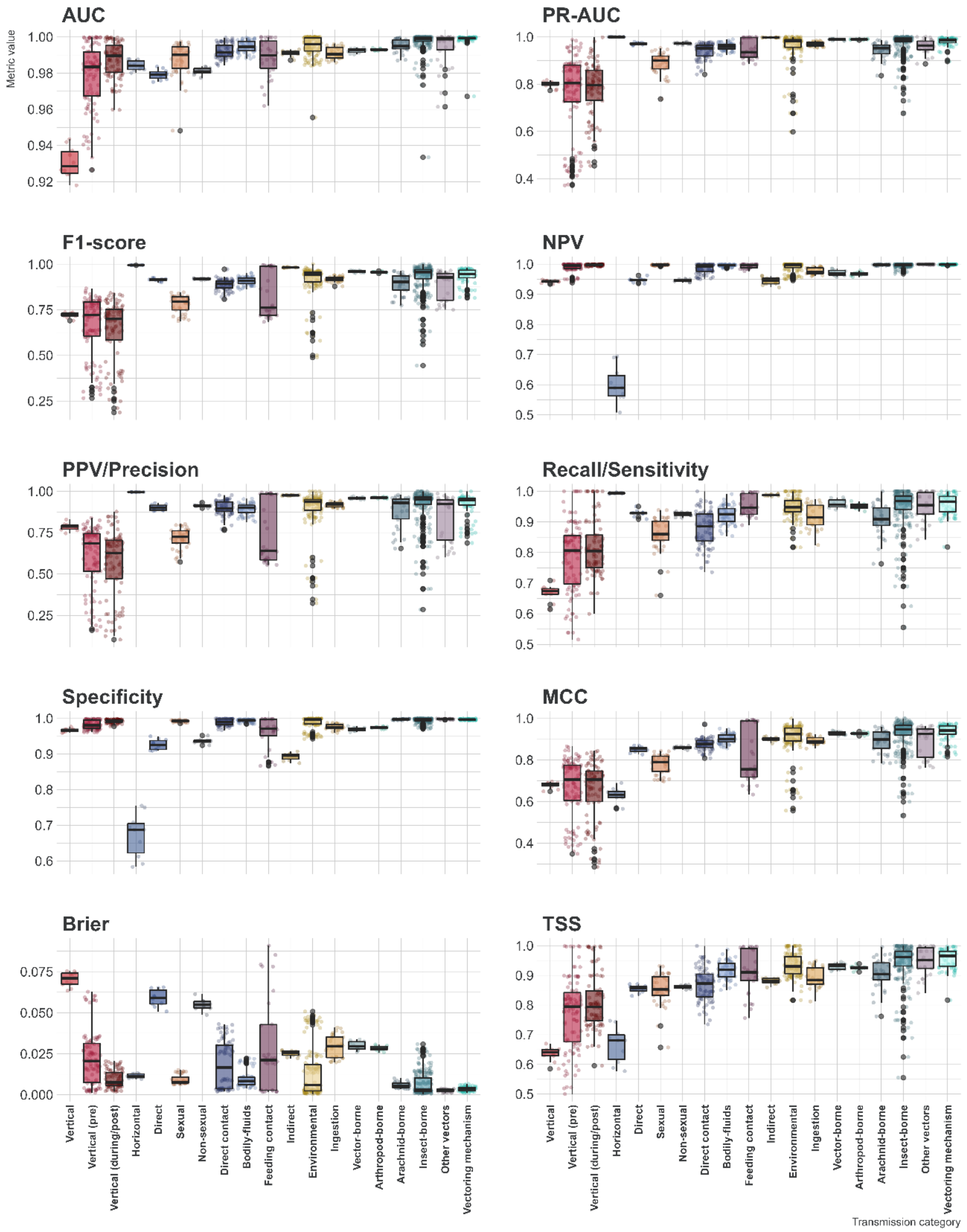


**Supplementary Figure 20 – Performance assessment of constituent class-balancing ensembles of against 50 held-out test sets at >0.5 probability threshold.** Points represent the class-balancing ensemble mean values for each performance metric (50 points per route/mode, 98 routes/modes in total). Boxplots represent the interquartile range (IQR), of the data distribution per each category of transmission route/mode. Horizontal lines within the box represent the median of the data distribution. Whiskers extend from the edges of the box to the minimum and maximum values within a distance of 1.5 times the IQR from the nearest quartile, individual data points that fall outside the range covered by the whiskers are plotted as outliers. Supplementary Table 9 provides full definitions of included performance metrics. For Brier score values closer to 0 indicate better performance, and those closer to 1 indicate worse performance.

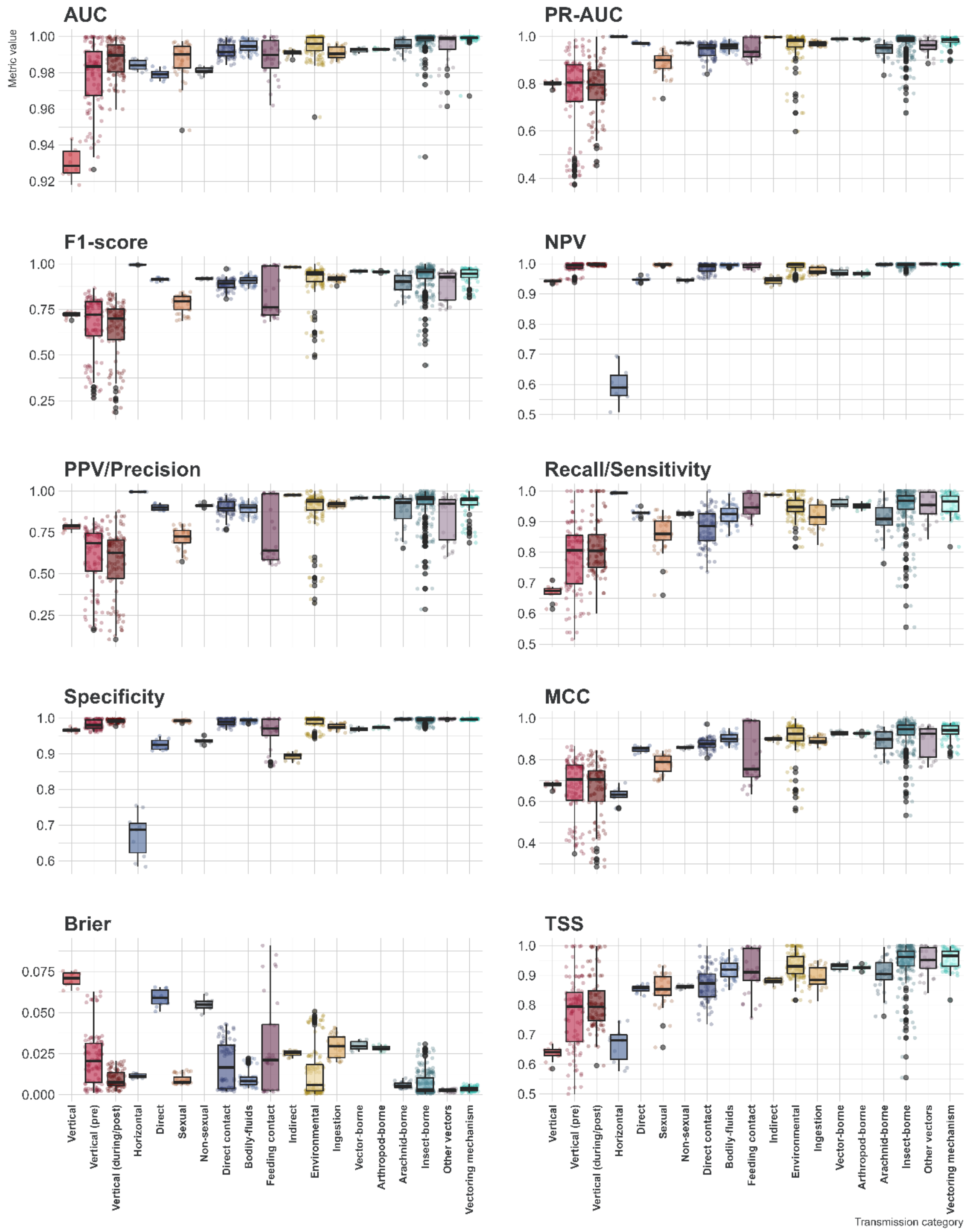


**Supplementary Figure 21 – Performance assessment of constituent class-balancing ensembles of our top-10 selected ensembles against 10 held-out test sets at >0.5 probability threshold.** Points represent the class-balancing ensemble mean values (5 models per each run) for each performance metric (20 points per route/mode, 98 routes/modes in total). Boxplots represent the interquartile range (IQR), of the data distribution per each category of transmission route/mode. Horizontal lines within the box represent the median of the data distribution. Whiskers extend from the edges of the box to the minimum and maximum values within a distance of 1.5 times the IQR from the nearest quartile, individual data points that fall outside the range covered by the whiskers are plotted as outliers. Supplementary Table 9 provides full definitions of included performance metrics. For Brier score values closer to 0 indicate better performance, and those closer to 1 indicate worse performance.

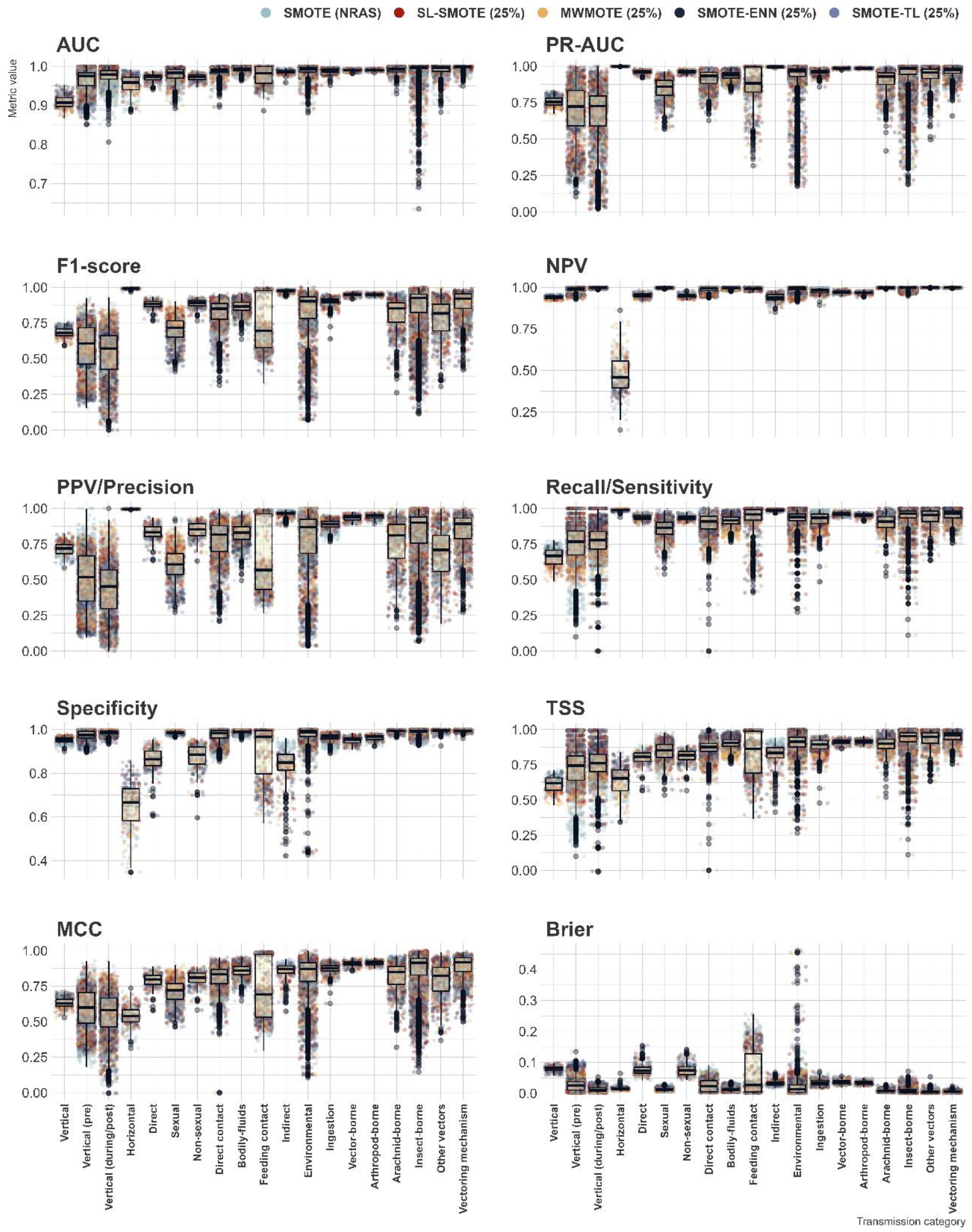


**Supplementary Figure 22 – Performance assessment of all trained models against 50 held-out test sets (at >0.5 probability threshold).** Points represent performance measured for constituent models per metric (250 points per route/mode, 98 routes/modes in total), and are coloured by the underlying class-balancing technique. Boxplots represent the interquartile range (IQR), of the data distribution per each category of transmission route/mode. Horizontal lines within the box represent the median of the data distribution. Whiskers extend from the edges of the box to the minimum and maximum values within a distance of 1.5 times the IQR from the nearest quartile, individual data points that fall outside the range covered by the whiskers are plotted as outliers. Supplementary Table 9 provides full definitions of included performance metrics. For Brier score values closer to 0 indicate better performance, and those closer to 1 indicate worse performance.

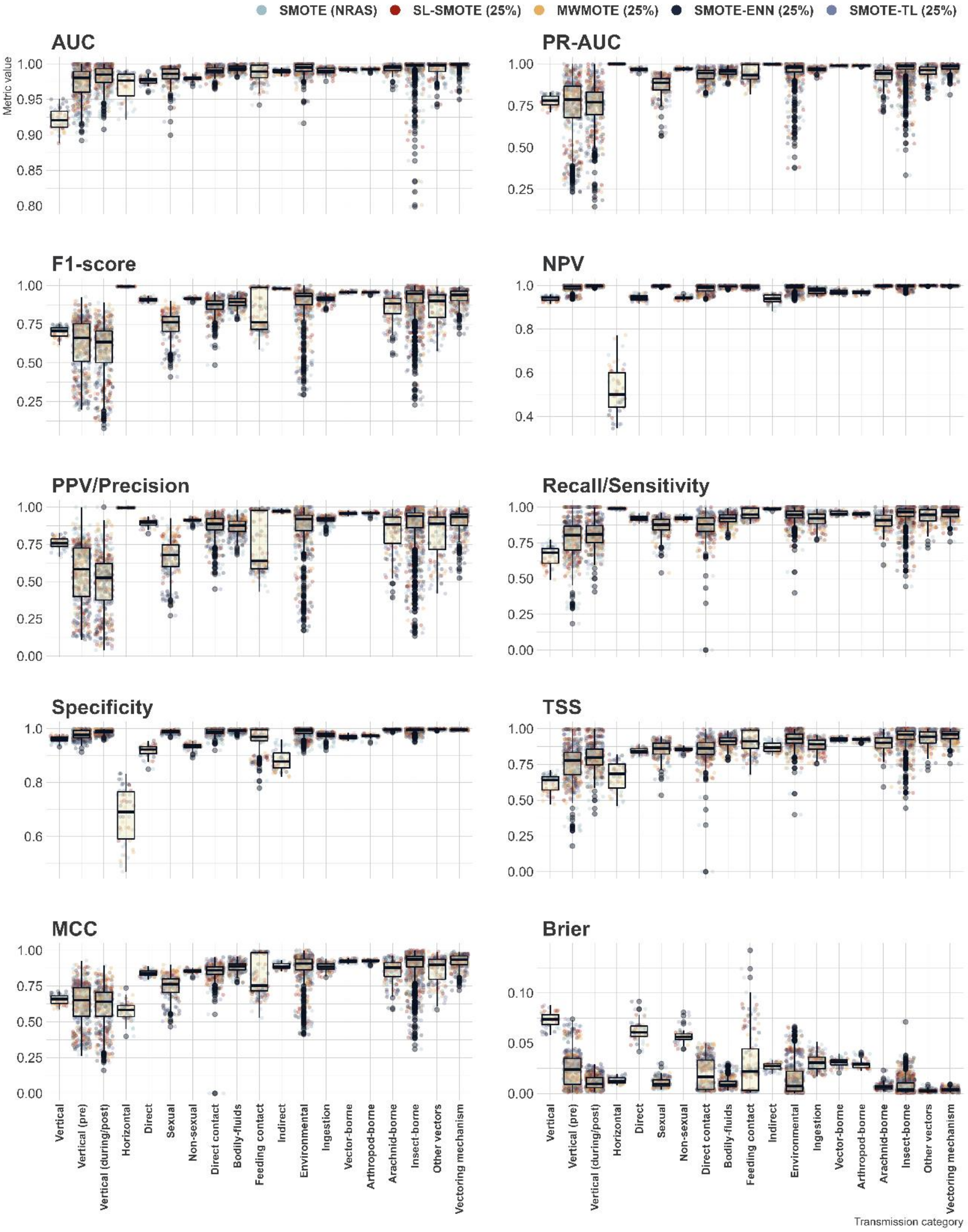


**Supplementary Figure 23 – Performance assessment of constituent models of our top-10 ensembles against 10 held-out test sets (at >0.5 probability threshold).** Points represent performance measured for constituent models per metric (50 points per route/mode, 98 routes/modes in total), and are coloured by the underlying class-balancing technique. Boxplots represent the interquartile range (IQR), of the data distribution per each category of transmission route/mode. Horizontal lines within the box represent the median of the data distribution. Whiskers extend from the edges of the box to the minimum and maximum values within a distance of 1.5 times the IQR from the nearest quartile, individual data points that fall outside the range covered by the whiskers are plotted as outliers. Supplementary Table 9 provides full definitions of included performance metrics. For Brier score values closer to 0 indicate better performance, and those closer to 1 indicate worse performance.

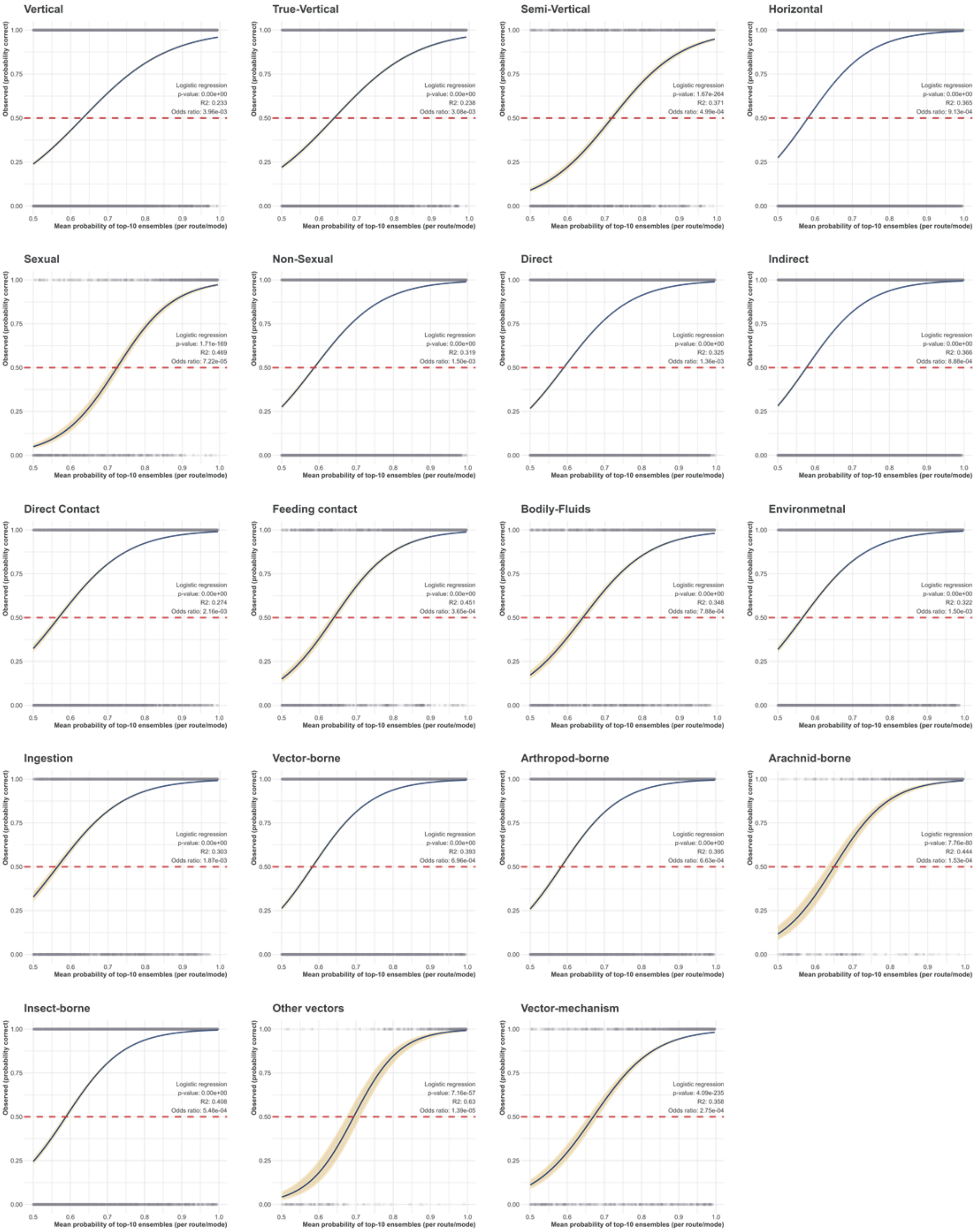


**Supplementary Figure 24 – Post-hoc assessment of in-sample predictions of our top-10 selected ensembles of five class balancing techniques at >0.5 mean probability threshold.** Logistic regression models for each mode (as per Supplementary Figures 18 and 19), relating the strength of mean prediction (probability averaged across the 10 ensembles per each constituent mode/route) to the prediction outcome. The blue line is the model prediction with standard error (yellow shading). Points are resulting outcomes (1 = observed, 0 = not observed), for all modelled routes/modes (as per Figure1 – Horizontal figure contains all predictions with probability>0.5, for all horizontal modes (e.g. indirect, direct) and routes (e.g. oral-faecal, mosquito-borne)). The dashed line is the null accuracy, defined here as 0.5 (random).
